# Thermally-induced neuronal plasticity that mediates heat tolerance

**DOI:** 10.1101/2024.08.06.606929

**Authors:** Wojciech Ambroziak, Sara Nencini, Jörg Pohle, Kristina Zuza, Gabriela Pino, Sofia Lundh, Carolina Araujo-Sousa, Larissa I. L. Goetz, Katrin Schrenk-Siemens, Gokul Manoj, Mildred A. Herrera, Claudio Acuna, Jan Siemens

## Abstract

Heat acclimation is an adaptive process that improves physiological performance and supports survival in the face of increasing environmental temperatures. Understanding the underlying mechanisms holds potential to mitigate health risks and reduces the steadily increasing number of heat-related casualties associated with global warming. Here we report the identification of a discrete group of hypothalamic preoptic neurons that transform to rheostatically increase their activity over the course of heat acclimation, a property required for mice to become heat tolerant. Peripheral thermo-afferent pathways via the parabrachial nucleus activate preoptic neurons and mediate acute heat-defense mechanisms in non-acclimated animals. However, long-term heat exposure promotes the preoptic neurons to gain intrinsically warm-sensitive activity, independent of thermo-afferent parabrachial input. Our data shows that their newly gained cell-autonomous warm-sensitivity is required to recruit peripheral heat tolerance mechanisms in acclimated animals. Mechanistically, we find a combination of increased sodium leak current and enhanced utilization of the Na_v_1.3 ion channel to drive their pacemaker-like, warm-sensitive activity. We propose a salient neuronal plasticity mechanism, adaptively driving acclimation to promote heat tolerance.

**Highlights:**

- Heat acclimation induces tonic, warm-sensitive firing in hypothalamic VMPO neurons
- Tonic activity in VMPO neurons primes peripheral organs to gain heat tolerance capacity
- Warm-sensitive tonic firing recruits heat tolerance mechanisms in acclimated animals
- Na_V_1.3 persistent sodium currents drive tonic, warm-sensitive firing in VMPO neurons

## Introduction

Prolonged exposure to hot (but non-lethal) temperatures enhances thermoregulatory responses in peripheral organ systems to rheostatically maintain body temperature within physiological limits, an adaptive phenomenon commonly referred to as heat acclimation. It has been proposed that the central nervous system (CNS) regulates these adaptive changes^1–4^.

While hypothalamic thermoregulatory pathways orchestrating long-term acclimation and heat tolerance are unknown, several hypothalamic cell populations have been described that mediate acute heat loss responses. These neurons reside in the rostral part of the hypothalamic preoptic area (POA) with the median preoptic nucleus (MnPO) at its center, an area that from here on we refer to as anterior ventromedial preoptic area (VMPO). A subset of VMPO neurons has been shown to respond to acute heat exposure and, in accordance with their predicted homeostatic function, acute optogenetic and chemogenetic stimulation of these ––largely glutamatergic–– neurons triggers prompt heat loss responses and body cooling^5–12^. However, it is not known whether POA neurons also control long-lasting rheostatic adaptations, to promote heat tolerance as a consequence of acclimation.

We here tested the hypothesis that long-term heat exposure during acclimation triggers plastic changes in the hypothalamic thermoregulatory area to regulate heat tolerance in mice.

## Results

### VMPO^LepR^ neurons gain intrinsic warm-sensitivity upon heat acclimation

Acute exposure to hot environmental temperatures activates a subset of VMPO neurons to express the activity marker cFos^5,9,10,13–17^. We hypothesized that long-term heat exposure would alter the activity profile of these VMPO <u>W</u>arm-<u>R</u>esponsive <u>N</u>eurons (VMPO^WRN^), based on the premise that long-lasting thermo-afferent input could induce plastic changes and cellular adaptation.

To assess whether exposure to warm/hot ambient temperatures (36°C) over an extended time period would change VMPO^WRN^ activity, we used a cFos-based genetic mouse model, the so-called FosTRAP2 mouse line, that allows unbiased labelling of activated neurons^18^. We captured VMPO^WRN^ by exposing FosTRAP2 mice to 36°C for 4 or 8 hours. The pattern of “warm- TRAPped” neurons, visualized by the expression of nuclear GFP (nGFP) under the control of the FosTRAP2 mice (FosTRAP2;HTB), recapitulated the previously described cFos expression pattern of VMPO^WRN^ ^8^ (Extended Data Fig. 1a), demonstrating that FosTRAP2;HTB mice allow permanent labelling of WRN. Moreover, longer heat exposure (8 vs. 4 hours) resulted in increased TRAPping of neurons, suggesting that progressively more neurons within the preoptic network are recruited upon longer heat exposure (Extended Data Fig. 1a).

Next, we “warm-TRAPped” FosTRAP2;HTB animals for either 4 or 8 hours and subsequently acclimated them at 36°C for ≥ 4 weeks, a time period required to reach full heat acclimation in rodents^19^. Finally, we prepared acute brain slices for electrophysiological recordings (Fig. 1a).

**Fig. 1.**
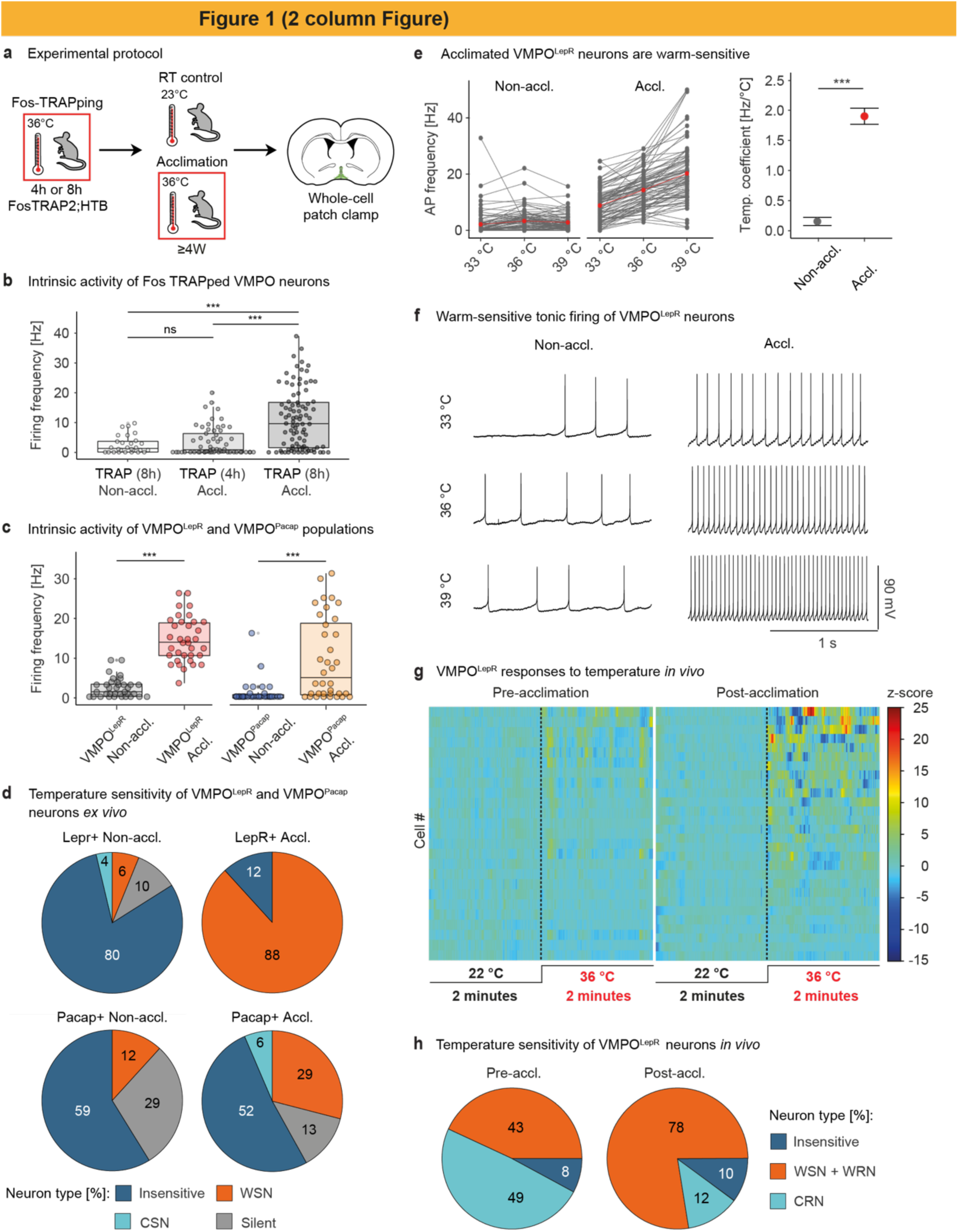
Heat acclimation increases warm-sensitive tonic AP firing of VMPO^LepR^. a, Cartoon depicting the protocol of acclimation to heat (36°C) for ≥ 4 weeks, following FosTRAPping at 36°C for 4 or 8 hours. Acclimated (Accl., 36°C) and non-acclimated (Non-accl., 22°/23°C) FosTRAP animals were used for acute brain slice electrophysiological recordings. b, Spontaneous action potential frequency in neurons of short (4h) and long (8h) warm-TRAPped mice subsequent to heat-acclimation (Accl.). Neuronal activity was recoded at 36°C bath temperature. Boxplots show median and interquartile range. One-way ANOVA, P < 0.0001; Tukey’s multiple comparison test, P = 0.8364 (TRAP (8h) Non- accl.: TRAP (4h) Accl.), ***P < 0.0001 (TRAP (8h) Non-accl.: TRAP (8h) Accl.), ***P < 0.0001 (TRAP (4h) Accl.: TRAP (8h) Accl.). n = 28/3 (TRAP (8h) Non-accl.), n = 22/2 (TRAP (4h) Accl.) and n = 33/3 (TRAP (8h) Accl.) cells. c, Comparison of spontaneous firing AP frequency in non-acclimated vs heat- acclimated VMPO^LepR^ (n = 35/6 for non-acclimated and n = 35/6 for acclimated) and VMPO^Pacap^ (n = 30/3 for non-acclimated and n = 37/3 for acclimated) neurons. Neuronal activity was recoded at 36°C bath temperature. Boxplots show median and interquartile range. Unpaired two-tailed t-test, ***P ≤ 0.0001 (VMPO^LepR^) and ***P ≤ 0.0001 (VMPO^Pacap^). d, Distribution of temperature-insensitive, cold-sensitive (CSN, temperature coefficient < -0.6 Hz/°C), warm-sensitive (WSN, temperature coefficient ≥ 0.75 Hz/°C) and silent neurons within the non-acclimated and acclimated VMPO^LepR^ (n = 81/9 for non-acclimated and n = 85/10 for acclimated) and VMPO^Pacap^ (n = 17/3 for non-acclimated and n = 31/3 for acclimated) neuron populations *ex vivo* recorded at 33°C, 36°C and 39°C bath temperature. e, Left: firing frequencies of non-acclimated (n = 81/9) and acclimated (n = 85/10) VMPO^LepR^ neurons at three bath temperatures recorded under synaptic blockade; individual cells are plotted in grey, red points represent group averages. Right: temperature coefficient (Hz/°C; mean ± s.e.m.) comparison between the non-acclimated and acclimated VMPO^LepR^ neurons. Unpaired two-tailed t-test, ***P ≤ 0.0001. f, Example traces of a non-acclimated and acclimated VMPO^LepR^ neuron activity recorded at 33°C, 36°C and 39°C. g, Heatmaps displaying *in vivo* single cell VMPO^LepR^ responses at 22°C and 36°C, before (left) and after (right) 30 days of heat acclimation. h, Pie charts showing fractions of neurons either increasing (warm-sensitive and warm-responsive neurons, WSN + WRN), decreasing (cold-responsive neurons, CRN) or not changing (Insensitive) activity of VMPO^LepR^ neurons upon increasing ambient temperature acutely from 22°C to 36°C, before and after acclimation to heat. Number of detected cells Pre-accl.: WSN + WRN: 22, CRN: 25, Insensitive: 4; Post- accl.: WSN + WRN: 38, CRN: 6, Insensitive: 5. N=4 mice. Neuronal activity *ex vivo* was recoded under fast synaptic transmission blockade. Acclimation was performed with mice housed at 36°C for ≥ 4 weeks with *ad libitum* access to water and food (see also Extended Data Figure 1-4).

Strikingly, longer-TRAPped neurons –but not shorter-TRAPped neurons– showed increased action potential (AP) firing when FosTRAP2;HTB mice were acclimated (Fig. 1b).

Neuronal activity *ex vivo* was recoded under fast synaptic transmission blockade. Acclimation was performed with mice housed at 36°C for ≥ 4 weeks with *ad libitum* access to water and food (see also Extended Data Figure 1-4).

Leptin receptor- (LepR-) and PACAP/BDNF- expressing VMPO neurons (VMPO^LepR^ and VMPO^PACAP^) have been found to partially overlap with the VMPO^WRN^ population, with subfractions of them co-expressing the activity marker cFos when mice are placed at warm temperatures^8,10,20^, a finding that we confirmed (Extended Data Fig 1b, c).

Moreover, VMPO^LepR^ and VMPO^PACAP^ can drive heat loss responses when activated chemogenetically and optogenetically^10,8^ (Extended Data Fig. 1d), consistent with a role in thermoregulation during heat exposure. We therefore wondered whether VMPO^LepR^ and/or VMPO^PACAP^ neurons would also change their activity profile upon long-term heat acclimation. We heat acclimated animals expressing a green fluorescent reporter under the control of the leptin receptor gene (LepR-Cre;HTB) or the PACAP gene (PACAP;EGFP) for ≥ 4 weeks at 36°C. Indeed, we also found VMPO^PACAP^ and VMPO^LepR^ neurons to increase AP firing upon long-term heat acclimation, with the smaller LepR-positive population appearing to plastically transform more robustly (Fig 1c).

We noted that 8-hour warm-TRAPping labelled neurons with a greater potential to subsequently become acclimation-activated compared to shorter (4-hour) TRAPping (Fig 1b). Interestingly, this result mirrored warm-induced cFos-labelling of VMPO^LepR^ neurons: although native cFos expression follows an overall faster kinetic than cFos-TRAPping, a significant fraction of cFos-positive cells coincided with VMPO^LepR^ neurons only after 4 hours but not yet after 2 hours (Extended Data Fig. 1e), suggesting that those VMPO neurons that slowly respond to prolonged thermal stimuli transform into acclimation-activated neurons rather than rapid responders.

To further evaluate the specificity of the observed acclimation-induced plasticity, we randomly sampled unlabeled VMPO neurons of similar size compared to VMPO^LepR^, assessed by cellular capacitance measurements (Extended Data Fig. 1f and 2a), to find that acclimation-induced plasticity is not a general phenomenon of all (randomly selected) VMPO neurons (Extended Data Fig. 2b).

Several recent studies suggest that heat loss responses are largely mediated by glutamatergic (Vglut2-positive) rather than GABAergic (Vgat-positive) VMPO neurons^6,9,10,21,22^. In line with these observations, we found Vglut2-positive (but not Vgat-positive) VMPO neurons to be enriched in the heat acclimation-induced population. However, their acclimation-induced response profile appeared more heterogeneous compared to VMPO^LepR^, with a considerable subset of VMPO^Vglut2^ being silent or near-silent (Extended Data Fig 2b). The observed VMPO^Vglut2^ (and VMPO^PACAP^) response heterogeneity correlates with the presumed larger cell-molecular diversity of these two populations compared to the smaller VMPO^LepR^ population^9^.

In contrast, cold-responsive LepR-positive neurons residing in the dorsal medial hypothalamus (DMH^LepR^)^23–25^ did not increase their firing rates upon heat acclimation (Extended Data Fig. 2c). Importantly, in both TRAPped WRNs (Fig. 1b) and in VMPO^LepR^ neurons (Extended Data Fig. 2d, e), inhibiting fast synaptic transmission did not affect the increased AP firing, indicating induction of a cell-autonomous, tonic pacemaker-like mechanism by heat acclimation.

Intriguingly, tonic activity is a characteristic feature of the so-called warm-sensitive neurons (WSN) that increase their activity (spontaneous action potential firing rate / fAP) upon temperature (T_core_) increase, presumably to mount appropriate heat loss responses. Traditionally, WSN are identified *ex vivo* in brain slice preparations by monitoring their fAP while warming the temperature of the perfusion fluid^26^. However, their physiological role and significance is not fully understood, largely because specific molecular markers for this cellular population have not been _found13,15,16,27,28._

We hypothesized that VMPO^LepR^ might be the long sought-after WSN. However, non-acclimated VMPO^LepR^ showed little to no warm-sensitivity. Strikingly, heat acclimation transformed the majority of VMPO^LepR^ into robust, cell-autonomous WSNs (Fig. 1d-f).

We wondered whether acclimated and non-acclimated neurons would become indistinguishable at a bath temperature of around 29.1°C, which was predicted by regression analysis (Fig. 1e and Extended Data Fig. 2f). Indeed, at recording temperatures of 30°C and lower, the firing rates became indistinguishable (Extended Data Fig. 2f), suggesting that the decisive difference of non-acclimated vs acclimated VMPO^LepR^ is their acquired warm-sensitivity in the physiological temperature range (36°C-39°C). Acclimation-induced warm-sensitivity was lower in PACAP- and VGlut2-positive VMPO neurons (Fig. 1d and Extended Data Fig. 2g, h). Moreover, tonic temperature-dependent firing of VMPO^LepR^ became highly regular as a consequence of acclimation. Again, this feature, assessed by determining the coefficient of variation of the interspike interval (ISI_Cov_), was most pronounced in acclimated VMPO^LepR^ neurons compared to any other population analyzed (Extended data Fig. 2i,j).

**Fig 2.**
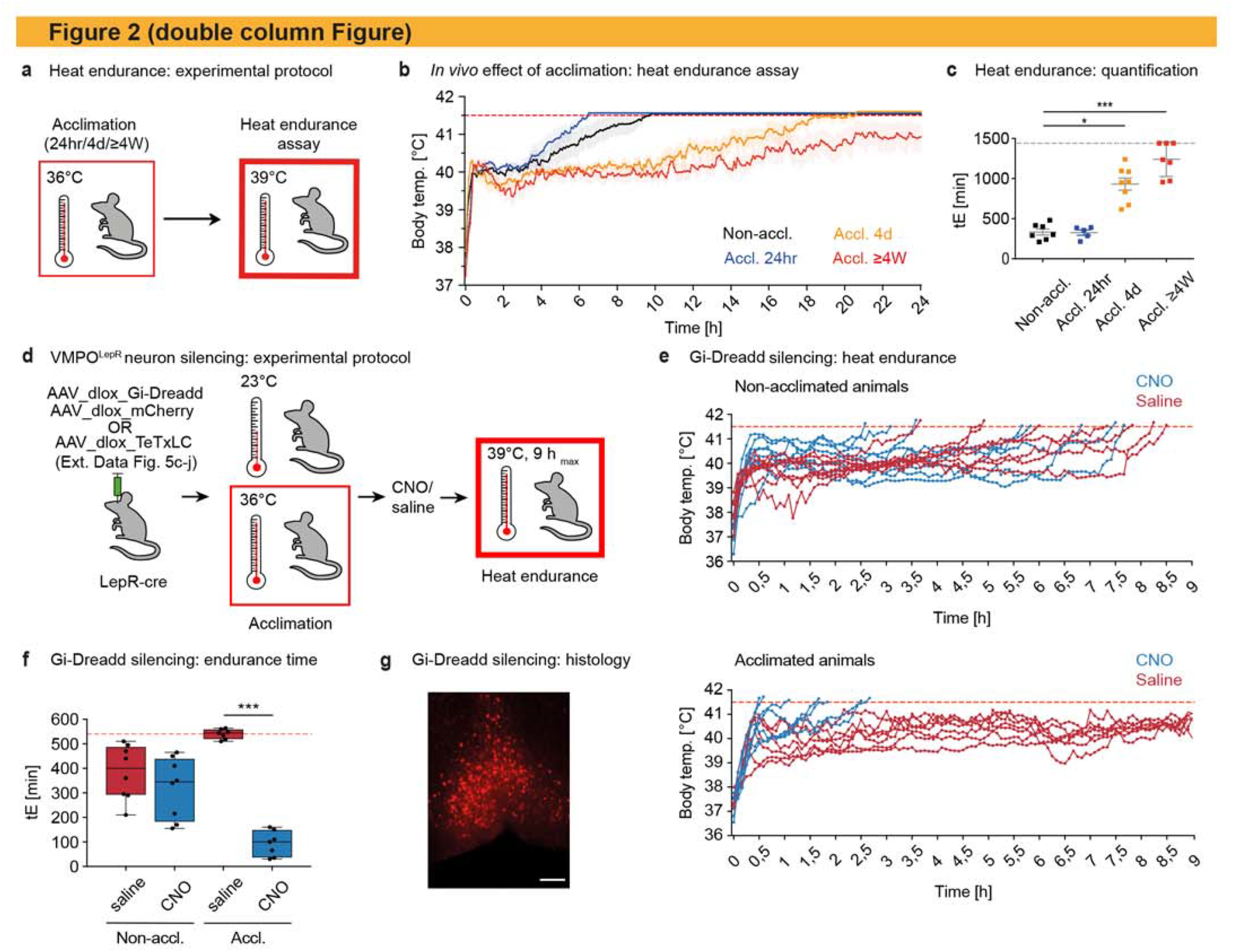
Enhanced heat tolerance after heat acclimation depends on VMPO^LepR^ neuron activity. a, Heat endurance assay design following heat acclimation for variable times. b, Average body temperature (mean ± s.e.m.) of non-acclimated (black, N = 7), 24 h (blue; N = 5), 4 days (orange; N = 8) and 4-5 weeks (red; N = 7) acclimated animals in the heat endurance assay monitored for a maximum of 24 hours or until the animal reached the cut-off temperature of 41.5°C (dashed red line); animals that reached the cut-off time were removed from the assay. c, Endurance time (tE, min) of mice shown on the left. Cut-off time = 24 hours (dashed grey line). Kruskal-Wallis test, P < 0.0001; Dunn’s multiple comparisons test, *P = 0.0262 (Non-accl : Accl. 4d), ***P = 0.0006 (Non-accl. : Accl. ≥ 4 weeks). Error bars represent mean ± s.e.m. d, Schematic showing the two experimental strategies used to interfere with VMPO^LepR^ activity. LepR-Cre animals were POA-injected with Cre-dependent TeTxLC- or Gi-DREADD viral particles to permanently eliminate synaptic output from VMPO^LepR^ neurons or to temporarily silence VMPO^LepR^ neuron activity, respectively. e, Heat endurance assay of Gi-DREADD expressing mice. Non-acclimated (top panel) or acclimated (bottom panel) animals were injected either with CNO (0.3 mg/kg i.p.) or saline 10 min prior to the assay and the body temperature was continuously monitored. Non-acclimated animals endured for similar short times, independently whether they received CNO or vehicle (saline). In acclimated mice, CNO injection (but not saline injection) eliminated acquired heat tolerance and the animals quickly reached the cut-off temperature (41.5°C). f, Endurance time (tE) for the groups shown in (e). Boxplots show median and interquartile range. Kruskal-Wallis test, P < 0.0001; Dunn’s multiple comparisons test, ***P < 0.0001 (Accl. saline : CNO); N = 8 animals for Non-accl. groups and N = 7 for Accl. groups. Note that due to the assay cut-off time of 9 h, the heat tolerance capacity (tE) of the Accl. saline treated group is underestimated. g, Representative image of VMPO^LepR^ showing mCherry labelling of the Gi-DREADD-mCherry fusion protein; scale bar: 250 μm. Boxplots show median and interquartile range (see also Extended Data Figure 5 and 6).

Taken together, we found expression of the LepR gene in the VMPO to circumscribe a population of heat acclimation-activated neurons, with most VMPO^LepR^ acquiring warm-sensitive pacemaker activity.

### Heat acclimation enhanced warm-responsive VMPO^LepR^ activity *in vivo*

Next, we assessed whether heat acclimation also induced activity changes of VMPO^LepR^ neurons *in vivo*. To this end, we stereotactically delivered the Cre-dependent calcium sensor GCaMP6f into the VMPO of LepR-Cre mice and performed micro endoscopic (Miniscope-) imaging in freely moving mice^29^ before and after heat acclimation (Extended Data Fig. 3a,b). Indeed, acclimation increased the heat responsiveness of VMPO^LepR^ *in vivo*. Not only did we find more VMPO^LepR^ respond to a heat challenge subsequent to acclimation, but the neurons also responded more robustly (Fig. 1g, h, Extended Data Fig. 3c-g and Supplementary Videos 1, 2). This observation agrees with findings showing that increases in body temperature (upon a heat challenge) are directly transferred to POA neurons^27,30^.

Although it is technically challenging to register and follow individual neurons by Miniscope imaging over the extended acclimation period, such an analysis did not reveal an increase in acclimation-induced baseline tonic activity at 22°C ambient temperature (Extended Data Fig. 3h).

### Enhanced VMPO^LepR^ activity mediates heat tolerance

Heat acclimation-induced activity increases in VMPO^LepR^ were first detectable *ex vivo* after 4 days of heat acclimation, further increasing until reaching a maximum at about 4 weeks of acclimation (Extended Data Fig. 4a), a similar time frame is required to reach a fully heat acclimated state in rodents, resulting in their increased heat tolerance^31^.

When fully acclimated mice were returned to 23°C ambient temperature, AP firing in VMPO^LepR^ subsided to baseline levels within 7 days (Extended Data Fig. 4b). However, an “adaptive memory” remained: subsequent to a 7-day de-acclimation phase at 23°C, high AP firing rates in VMPO^LepR^ neurons were quickly retrieved when animals were placed again at 36°C for only 2 days, reaching significantly higher AP firing rates compared to naïve mice subjected to a 2-day acclimation period for the first time (Extended Data Fig. 4b). This property is reminiscent of acclimation-induced adaptations observed in peripheral organs promoting heat tolerance, which are also quickly recalled after primed acclimation^31,32^. We therefore wondered if heat acclimation-induced tonic activity in VMPO^LepR^ mediates this adaptive response and conveys heat tolerance.

Heat tolerance expands the limit of tolerable temperatures^33–35^. To assess beneficial autonomic effects of acclimatization *in vivo* and probe the tolerance to heat, we utilized a heat endurance assay during which the animal is challenged with hot ambient temperatures (39°C) while body temperature (T_core_) is monitored telemetrically (Fig. 2a and b)^36^. Non-acclimated mice were able to keep their T_core_ below 41.5°C ––demarcating the maximal T_core_ mice are able to tolerate^33,37^–– for an average endurance time (t_E_) of only 333.6 ±37.6 minutes (mean ± s.e.m.). Oppositely, animals acclimated at 36°C for ≥ 4 weeks were able to sustain T_core_ within the physiological range for long time periods (t_E_ = 1235 ±81.3) with some animals even exceeding a full circadian cycle (Fig. 2b and c and Extended Data Fig. 5a), attesting to the high heat tolerance level they had gained following acclimation. We found longer acclimation periods to enhance heat tolerance more robustly than shorter acclimation periods and, interestingly, increased heat endurance correlated with increased average AP firing frequencies of VMPO^LepR^ neurons (Extended Data Fig. 5b).

To address whether acclimation-induced activity in ––and resulting synaptic output of–– VMPO^LepR^ is required for gaining heat tolerance, we silenced the cells by virally delivering Cre- dependent tetanus toxin light chain (TeTxLC)^38^ into the POA of LepR-Cre mice prior to acclimation. We verified effectiveness of TeTxLC silencing (Extended Data Fig. 5c). While T_core_ and overall behavior of TeTxLC-silenced animals was normal at ambient temperatures of 23°C (data not shown), the mice were compromised during the 36°C-acclimation phase and presented with higher T_core_ temperatures than littermate controls (Extended Data Fig. 5d-f); several animals reached 41.5°C during the first 2 days of acclimation and thus could not be tested in the heat endurance assay. Presumably, strong and permanent TeTxLC-mediated inhibition revealed that output of the fraction of rapidly heat-responsive VMPO^LepR^ in non-acclimated mice (Fig. 1g, h and Extended Data Fig. 1a, b) has a role in acute heat defense of the animals.

The remaining TeTxLC-silenced animals were able to complete the full 30-day acclimation cycle but, subsequently, failed the heat endurance assay and performed similar to non-acclimated control animals (Extended Data Fig. 5h, i). These data concur with the hypothesis that continuous neuronal output from VMPO^LepR^ during acclimation is necessary to acquire heat tolerance.

However, this experiment did not allow us to conclude whether heightened acclimation-induced warm-sensitive activity in VMPO^LepR^ is mediating increased heat tolerance following acclimatization and during the heat endurance assay. To inhibit acclimation-induced AP firing in VMPO^LepR^ acutely, we made use of chemogenetic interference using the inhibitory hM4Di (Gi-DREADD) receptor^39^ in LepR-Cre mice (Fig. 2d and Extended Data Fig. 6a). Different from the tetanus toxin approach, virally mediated Gi-DREADD expression in VMPO^LepR^ does not hinder acclimation-relevant adaptive changes to occur but only inhibits neuronal activity in the presence of the DREADD agonist clozapine N-oxide (CNO), which we verified in brain slice recordings (Extended Data Fig. 6b).

When VMPO^LepR^ neurons were chemogenetically silenced during the heat challenge period, acclimated animals failed to maintain T_core_ within physiological boundaries in the heat endurance assay (Fig. 2e-g and Extended Data Fig. 6c). Strikingly, Gi-DREADD mediated inhibition resulted in rapid hyperthermia and short endurance times in acclimated animals, while it did not accelerate hyperthermia in non-acclimated controls, demonstrating that acclimation-induced warm-sensitive AP firing of VMPO^LepR^ triggers the utilization of gained heat tolerance capacity.

Collectively, these results suggest that acclimation-induced VMPO^LepR^ warm-sensitive activity is necessary for both, building up heat tolerance capacity over the course of the acclimation period and recruiting heat tolerance mechanisms upon an acute heat challenge.

### Excitatory LPBN**→**POA pathway is critical for the induction of heat acclimation

In line with a reduction in body weight (Extended Data Fig. 7a), a decline in blood plasma leptin levels paralleled the increase in warm-sensitive firing when mice were heat acclimated (Extended Data Fig. 7b). Leptin signalling has been implicated in POA-orchestrated thermoregulation and body temperature adaption^40–42^. We therefore wondered if a reduction in leptin levels during acclimation is a prerequisite for ––or permissive to–– the induction of AP firing increases in VMPO^LepR^ neurons. We found that modulating leptin levels *in vivo*, either by food deprivation (which naturally lowers leptin levels) or by supplementing leptin by i.p. injections during acclimation only had a small or negligible effect on VMPO^LepR^ activity or the performance of acclimated animals in the heat endurance assay, respectively (Extended Data Fig. 7c-h).

To assess whether the absence of leptin signalling may promote heat tolerance, we also tested if leptin receptor-deficient Db/Db mice^43^ would be better equipped to cope with 39°C heat without prior heat acclimation. However, we found that Db/Db mice did not perform longer in the heat endurance assay compared to their pair-fed and weight-matched littermate controls (Extended Data Fig 7i). We thus concluded that the reduction in leptin levels has a minor role in shaping VMPO^LepR^ activity and heat acclimation.

Given the results, we hypothesized that synaptic transmission could serve as an initial trigger of the observed neuronal plasticity mechanism. Intriguingly, at early stages (∼17 hours after placing animals at 36°C) but not at late stages of heat acclimation we found VMPO^LepR^ neurons to receive a higher frequency of excitatory synaptic inputs compared to non-acclimated animals (Extended Data Fig. 8a).

These findings suggested that heat-driven, thermo-afferent excitatory synaptic inputs to VMPO^LepR^ could be involved in triggering their plasticity and warm-sensitive tonic firing. Previously, the lateral parabrachial nucleus (LPBN) has been shown to constitute a major hub for thermo-afferent pathways that are relayed to the rostral POA^13,14,17^. We therefore wondered whether synaptic LPBN→VMPO transmission is important for acclimation. Thermoregulatory LPBN neurons innervating the POA are Vglut2-positive^17^. This allowed us to use Vglut2-Cre mice in combination with a dual viral-delivery strategy to selectively silence those LPBN projections reaching the VMPO: first we stereotactically supplied Cre-dependent FlpO retroAAV particles (designed to infect axonal nerve terminals^44^) into the POA. Subsequently, we injected AAV particles expressing FlpO-dependent TeTxLC into the LPBN (Fig. 3a, b). While Silencing LPBN→POA transmission did not alter baseline T_core_ of mice kept at normal (23°C) ambient temperatures, it prevented acclimation, and mice were unable to maintain T_core_ within the physiological range when placed at 36°C (Fig. 3c), a result similar to that observed when silencing VMPO^LepR^ directly (Extended Data Fig. 5g).

**Fig. 3.**
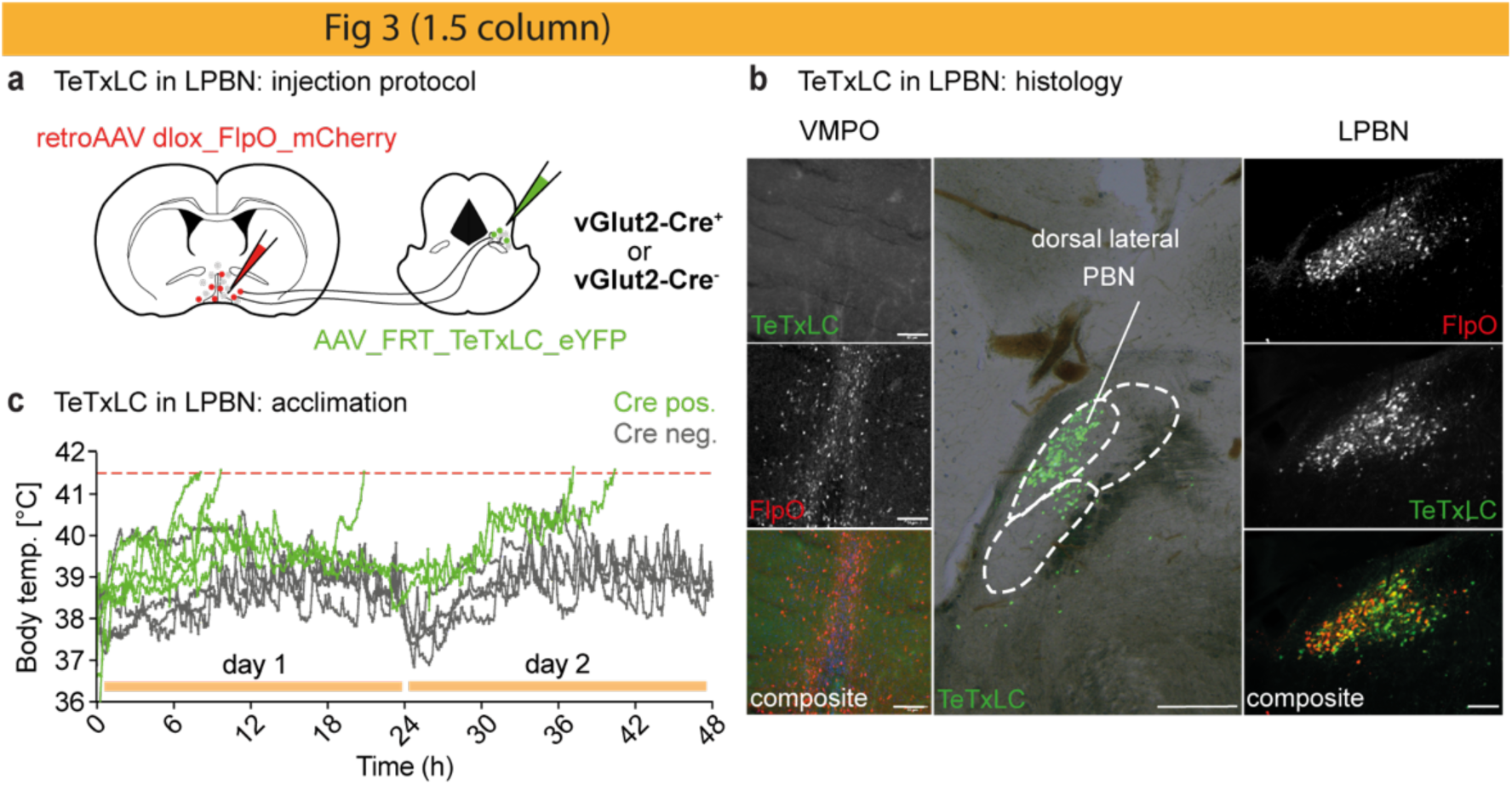
Thermo-afferent LPBN pathway is required to trigger heat acclimation. a, Left: Schematic showing the viral injection strategy for TeTxLC-mediated silencing of excitatory (Vglut2-positive) presynaptic neurons located in the LPBN and innervating VMPO. b, Example images showing the expression of AAV-FRT-TeTxLC-EGFP (green) and retroAAV-dlox-FlpO-mCherry (red) in the VMPO (left panel) and in VMPO-projecting LPBN neurons (right panel), scale bar: 250 μm. The histological labelling confirmed double infection of glutamatergic LPBN neurons in Vglut2-Cre mice expressing the recombinase FlpO (red, derived from the retro-AAV injected into VMPO) and TeTxLC (green, derived from Cre- and FlpO-dependent AAV particles injected into the LPBN) (middle panel), scale bar: 100 μm. Note that labelled neurons are mainly located in the dorsal lateral part of the LPBN; no TeTxLC is detectable in the POA (top left image), assuring that inhibition happened at the level of the LPBN but not the POA. c, Body temperature traces of individual LPBN→VMPO silenced (Cre pos., green, N = 5) and non-silenced control (Cre neg., grey, N = 5) animals during the initial 48 hours of heat acclimation. In contrast to Cre-negative animals, all animals expressing TeTxLC failed to maintain their body temperature below 41.5 °C during the first two days of acclimation (see also Extended Data Figure 7 and 8).

However, unlike transiently blocking VMPO^LepR^ neurons at the end of the acclimation period, transiently blocking VMPO-projecting LPBN neurons after a 30-day heat acclimation period did not abrogate heat tolerance: we again used a dual viral delivery strategy (Extended Data Fig. 8b) but in this case to temporarily silence LPBN→POA projection neurons via Gi-DREADD. Injecting CNO into Vglut2-Cre mice at the end of their long-term acclimation phase slightly (and fairly briefly) increased T_core_ (Extended Data Fig. 8c), but did not interfere with their performance in the heat endurance assay (Extended Data Fig. 8d). Subsequently, we verified the effectiveness of Gi-DREADD receptors in silencing LPBN→POA projection neurons *ex vivo* (Extended Data Fig. 8e).

These results are consistent with a role of LPBN→VMPO projections in the initial induction of heat acclimation, but this pathway plays only a minor role, if any, in driving and sustaining long-term heat acclimation.

Congruent with this hypothesis, we found that Trpv1-Cre;DTA mice, lacking most ––but not all– – peripheral thermosensory neurons due to the genetically-controlled expression of the diphtheria toxin^45,46^, were slightly but significantly hyperthermic at the beginning of the acclimation phase (Extended Data Fig. 8f and g), in agreement with previous acute heat-challenge results^45^. However, body temperatures recovered to normal levels by days 2 to 3 of acclimation, indicating that, with a delay, another (peripheral or central) mechanism was able to compensate for reduced primary afferent thermosensory signals. Similarly, the TRPM2 ion channel, which previously has been implicated in the acute detection of warm/hot temperatures in the peripheral and central nervous systems^27,47,48^, appeared largely dispensable for long-term heat acclimation and both groups (TRPM2 KO and Ctrl mice) performed similar in the heat endurance assay (Extended Data Fig. 8h-k).

Together, these results suggest that thermo-afferent excitatory synaptic pathways via the LPBN are largely important at the beginning of heat acclimation presumably to trigger adaptive plasticity in VMPO^LepR^, which thereby become autonomous warm-sensitive pacemaker neurons.

### Long-lasting, continuous VMPO^LepR^ activity is sufficient to increase heat tolerance

We wondered if we could mimic this process by continued, long-term activation of VMPO^LepR^ neurons in the absence of a warming stimulus. To this end, we implemented a chemogenetic gain of function approach. Stereotactic viral delivery and Cre-dependent expression of the chemogenetic activator hM3Dq (Gq-DREADD)^39^ allowed us to stimulate the neurons repetitively by injecting CNO every 24 hours for 1, 5 and 10 days (Extended Data Fig. 9a, b). We found that “chemogenetic conditioning” of the animals by increasing the activity of VMPO^LepR^ neurons for 10 days (but not for 5 days or shorter) before the heat endurance assay was sufficient to induce increased heat tolerance and, somewhat surprisingly, also slightly increased tonic activity in VMPO^LepR^ neurons assessed in brain slice recordings (Extended Data Fig. 9c-e).

Gq-DREADD-mediated chemogenetic stimulation of VMPO^LepR^ neurons induces pronounced hypothermia^10^ (Extended Data Fig. 9b), presumably by acutely triggering excessive neuronal activation, thereby potentially also explaining the initial dip in heat tolerance capacity after 5 days (Extended Data Fig. 9d). To have a more accurate control over firing rates of VMPO^LepR^, we next opted for long-term optogenetic stimulation ––optogenetic conditioning–– by expressing Cre- dependent channelrhodopsin (ChR2) in the POA of LepR-Cre animals (Fig. 4a). We optically stimulated VMPO^LepR^ neurons with a low stimulation frequency of 1Hz, which still triggered hypothermia albeit of lower magnitude compared to chemogenetic stimulation (Extended Data Fig. 9f). Optical stimulation of control mice absent of ChR2 did not have any measurable effect on T_core_ (Extended Data Fig. 9g), demonstrating that light-induced heating was minimal and did not affect this thermosensitive brain area^27^. Similar to chemogenetic conditioning, continuous optic stimulation for 3 days (but not for shorter periods) also resulted in increased thermotolerance and enhanced performance in the heat endurance assay (Fig. 4b, c and Extended Data Fig. 9h).

**Fig. 4.**
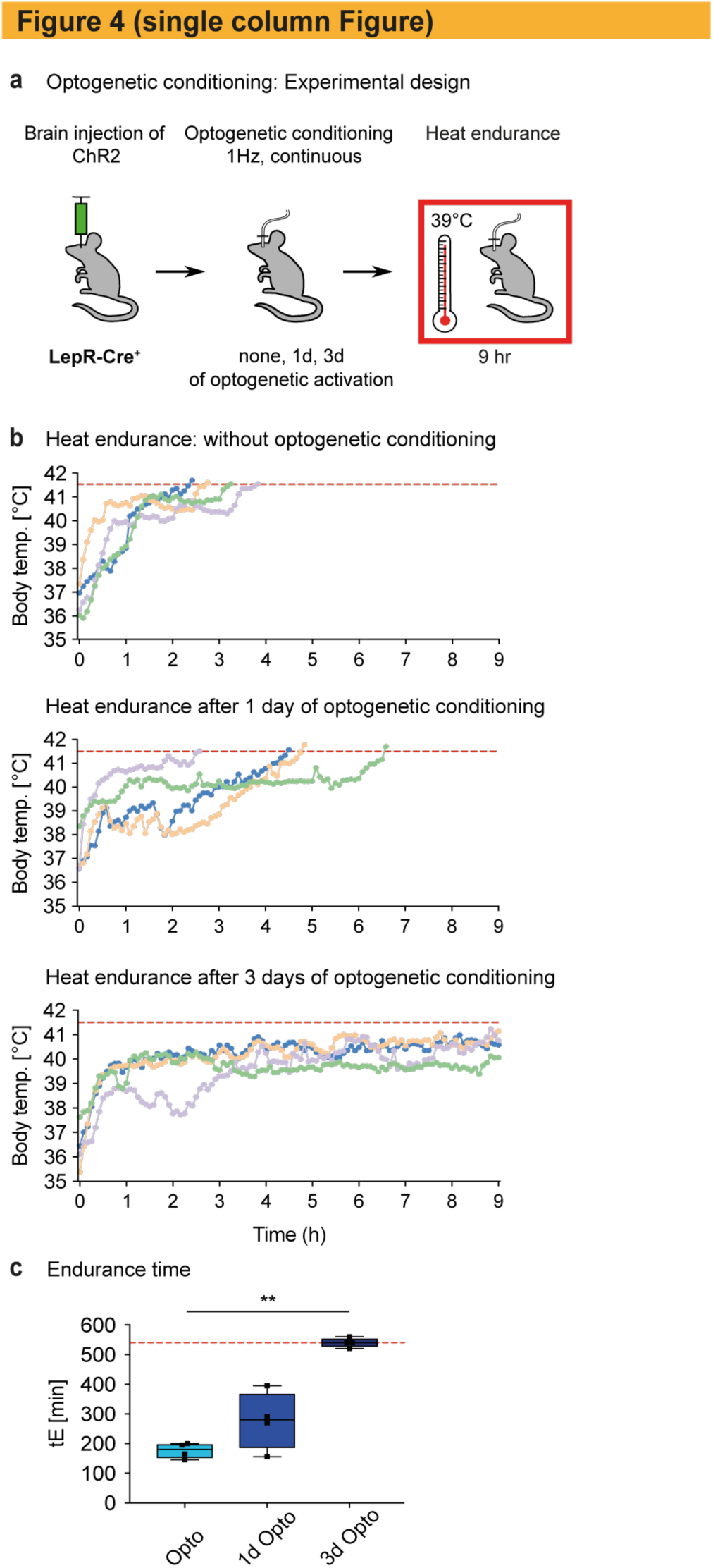
Optogenetic conditioning of VMPO^LepR^ neurons induces heat tolerance. a, Experimental paradigm used for continuous optogenetic activation of VMPO^LepR^ neurons prior to the heat endurance assay. LepR-Cre animals were injected with Cre-dependent ChR2 AAV particles into the rostral POA and either not stimulated or stimulated for 1 or 3 days by blue light at a low frequency (1 Hz) prior to the heat endurance assay. All animals were optogenetically stimulated during the heat endurance assay. b, Body temperature of individual mice subjected to optogenetic conditioning. Only those animals conditioned for 3 days had acquired heat tolerance and performed robustly in the heat endurance assay. Animals that reached the cut-off temperature of 41.5°C were removed from the assay; assay duration was limited to 9 hours. c, Endurance time (tE) of the differently conditioned groups shown in (b). Boxplots show median and interquartile range. Kruskal-Wallis test, P < 0.05; Dunn’s multiple comparisons test, **P = 0.0088 (Opto : 3d Opto). N = 4 per group (see also Extended Data Figure 9).

Collectively, these data demonstrate that long-term increases in VMPO^LepR^ activity is required and sufficient for the expression of heat tolerance.

### Ionic basis for acclimation-induced activity of VMPO^LepR^

To shed light on the molecular underpinnings of the acquired cell-autonomous, warm-sensitive VMPO^LepR^ activity, we analyzed electrophysiological changes that occurred in the face of acclimation.

We found the average resting membrane potential (RMP) to be depolarized by approximately 10mV in heat-acclimated VMPO^LepR^ compared to non-acclimated controls (Fig. 5a; Vm = -44.31 ±0.85 vs Vm = -54.00 ±1.98, P = 0.0001), possibly contributing to higher firing rates. In principle, a reduction of background K^+^ current can yield a more depolarized membrane potential. Instead of a decrease, we rather found a slight increase in overall K^+^ current in acclimated VMPO^LepR^ (Extended Data Fig. 10a), suggesting that leak K^+^ currents are not majorly contributing to acclimation-induced RMP depolarization. This conclusion is further supported by indistinguishable membrane input resistances for the acclimated and non-acclimated groups (Fig. 5b).

**Fig. 5.**
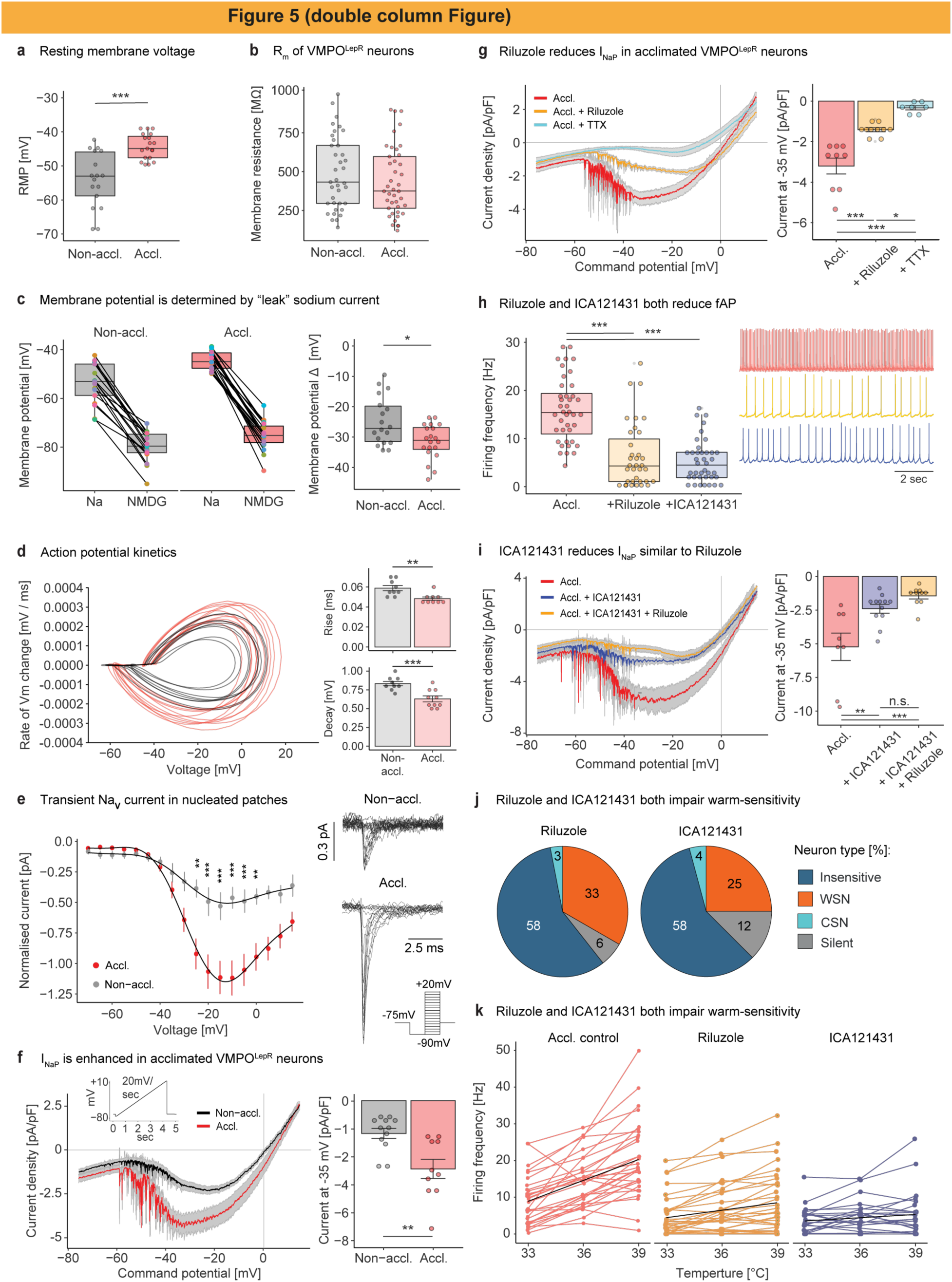
Electrophysiological characterization of VMPO^LepR^. a, Resting membrane potential (RMP) in acclimated VMPO^LepR^ neurons (n = 19/3) is depolarised compared to the RMP of non-acclimated VMPO^LepR^ (n = 17/3) cells. Unpaired two-tailed t-test, ***P = 0.0001. b, Membrane input resistance (Rm) is comparable between non-acclimated (n = 37/9) and acclimated (n = 41/10) VMPO^LepR^ neurons. c, Left: Membrane hyperpolarization in non-acclimated (n = 17/2) and acclimated (n = 19/2) VMPO^LepR^ neurons caused by replacement Na^+^ for NMDG^+^ in aCSF. Right: Difference in membrane potential (Δ) between Na^+^-based aCSF and NMDG^+^-based aCSF is larger in acclimated VMPO^LepR^ neurons. Unpaired two-tailed t-test, *P = 0.0201. d, Left: Action potential phase plot of non-acclimated (grey, n = 9/4) and acclimated (red, n = 10/5) VMPO^LepR^ neurons. Right: Both action potential 10% - 90% rise time (Wilcoxon-Test, **P = 0.0046) and 90% - 10% decay time (Unpaired two-tailed t-test, ***P = 0.0006) are significantly faster in VMPO^LepR^ neurons after acclimation. e, Left: Current-voltage relationship for VMPO^LepR^ neuron peak transient NaV currents recorded in nucleated patches. Two-way ANOVA (effect of acclimation * voltage), P < 0.0001; Tukey’s multiple comparison test, **P = 0.0016 (-25 mV), ***P = 0.0003 (-20 mV), ***P = 0.0002 (-15 mV), ***P < 0.0001 (-10 mV), ***P = 0.0005 (-5 mV) and **P = 0.0072 (0 mV). n = 6/2 (Non-accl.) and n = 6/2 (Accl.) cells. Right: Example transient NaV current recordings from VMPO^LepR^ nucleated patches obtained from acclimated Lepr-Cre;HTB mice or non-acclimated controls. Inset: voltage step protocol used. f, Left: Average INaP, revealed by slow depolarizing voltage ramp, is enhanced after heat acclimation. n = 12/4 (Non-accl.) and n = 10/4 (Accl.) cells. Inset: ramp protocol used to record INaP. Right: Quantification of INaP at -35mV based on data shown in the left panel. Unpaired two-tailed t-test, *P = 0.0055. g, Left: INaP in acclimated VMPO^LepR^ neurons is reduced by Riluzole (10 µM) and completely blocked by TTX (1 µM). Right: quantification of INaP at - 35mV based on data shown in the left panel. One-way ANOVA, P < 0.0001; Tukey’s multiple comparison test, ***P < 0.0001 (Accl. : Accl.+Riluzole), ***P < 0.0001 (Accl. : Accl.+TTX), *P = 0.0170 (Accl.+Riluzole : Accl.+TTX). n = 9/2 (accl.), n = 10/2 (Riluzole) and n = 7/2 (TTX) cells. h, Firing frequency (fAP) of acclimated VMPO^LepR^ is reduced by both, Riluzole (10 µM) and ICA121431 (200 nM). One-way ANOVA, P < 0.0001; Tukey’s multiple comparison test, ***P < 0.0001 (Accl. : Accl.+Riluzole), ***P < 0.0001 (Accl. : Accl.+ICA121431). n = 40/10 (Accl.), n = 35/4 (Riluzole) and n = 39/6 (ICA121431) cells. Right: Example traces of the 3 conditions are shown. i, Left: NaV1.3 antagonist ICA121341 blocks INaP in acclimated VMPO^LepR^ neurons to a similar extent as Riluzole. Right: quantification of INaP at -35 mV based on data shown in the left panel. One-way ANOVA, P = 0.0002; Tukey’s multiple comparison test, **P = 0.0029 (Accl. : Accl.+ICA121431), ***P = 0.0001 (Accl. : Accl.+ICA121431+Riluzole). n = 8/3 (Accl.), n = 12/4 (ICA121431) and n = 10/2 (ICA121431+Riluzole) cells. Part of the Accl. INaP data shown in (g) was repurposed for comparisons shown in (i). j, Distribution of temperature-insensitive, cold-sensitive (CSN), warm-sensitive (WSN) and silent neurons within acclimated VMPO^LepR^ population recorded either with Riluzole (10 µM) or ICA121431 (200 nM) in perfusion fluid. n = 33/4 for Riluzole and n = 24/4 for ICA121431. k, Firing frequencies of acclimated VMPO^LepR^ control cells (n = 30/5), acclimated VMPO^LepR^ cells recorded with Riluzole (n = 33/4) and acclimated VMPO^LepR^ cells recorded with ICA121431 (n = 24/4) at three bath temperatures. Individual cells are plotted in color; black lines represent linear regression for each group Tc (slope / temperature coefficient) = 1.9 for Accl. control, Tc = 0.68 for Riluzole and Tc = 0.29 for ICA121431). Accl. control cells were randomly sampled from the acclimated VMPO^LepR^ cells plotted in Fig. 1e. In panel (e), current amplitudes were normalized to nucleated patch circumference. In all other panels, currents were normalized to cell capacitance to avoid variability due to cell size. All ionic current recordings were conducted at 36°C bath temperature; action potentials were recoded under fast synaptic transmission blockade and using “high-K^+^ aCSF” (see methods for details). Data presented as median and interquartile range in (a), (b), (c) and (h); elsewhere data represents mean ± s.e.m (see also Extended Data Figure 10- 12).

Theoretically, cation-selective TRP ion channels could pass tonic depolarizing current to increase AP firing in VMPO^LepR^. Ruthenium Red, 2-Aminoethyl diphenylborinate (2-APB), and ML204 – ––broad spectrum inhibitors of heat-activatable TRPV1, TRPM2, TRPM3 and TRPC channels^6,30,49,50^–– had little or no effect on cation currents in acclimated VMPO^LepR^ (Extended Data Fig. 10b). Importantly, none of the substances had a significant impact on tonic AP firing (Extended Data Fig. 10c, d). Next to TRPM2, TRPC4 channels have recently been implicated in warm-sensitive AP firing of POA neurons^30^. ML204 and Pico145, two potent inhibitors of TRPC4 channels^51^, did not attenuate tonic AP firing and warm-sensitivity of heat-acclimated VMPO^LepR^, respectively (Extended Data Fig. 10d, e), suggesting that heat acclimation induces a TRPC4- independent molecular mechanism of warm-sensitivity.

The sodium “leak” channel NALCN^52^ has been described to modulate autonomous firing of other tonically active neurons, including suprachiasmatic nucleus (SCN) neurons that are neighboring the POA and that regulate the circadian cycle^53^. We found Na^+^ “leak” currents to contribute to the RMP in both non-acclimated and acclimated neurons with a significantly larger contribution in acclimated VMPO^LepR^ (Fig. 5c). To determine whether the difference in RMP could explain the AP firing increase, we depolarized the non-acclimated cells to a similar membrane potential observed in acclimated VMPO^LepR^. We found that depolarization of non-acclimated cells did not have a major impact on either frequency or regularity of action potential firing (Extended Data Fig. 10f), suggesting that mimicking a depolarized state is ––on its own–– insufficient to recapitulate their acclimation-induced firing pattern. Nevertheless, injecting a hyperpolarizing current into acclimated VMPO^LepR^ reduced their firing rate (Extended Data Fig. 10g), showing that a depolarization bias supports tonic VMPO^LepR^ activity.

Since passive conductances –– either K^+^ or Na^+^ –– did not fully explain the increased activity of acclimated VMPO^LepR^, we investigated the contribution of voltage-gated ion channels to tonic firing.

In panel (e), current amplitudes were normalized to nucleated patch circumference. In all other panels, currents were normalized to cell capacitance to avoid variability due to cell size. All ionic current recordings were conducted at 36°C bath temperature; action potentials were recoded under fast synaptic transmission blockade and using “high-K^+^ aCSF” (see methods for details). Data presented as median and interquartile range in (a), (b), (c) and (h); elsewhere data represents mean ± s.e.m (see also Extended Data Figure 10- 12).

Despite the observation that fast afterhyperpolarization was changed upon acclimation (Extended Data Fig. 11a), we found that ion channels typically carrying or being activated by the underlying current, such as Ca^2+^-activated large conductance K^+^ (BK) channels and HCN channels, respectively, did not appear to contribute to acclimation-induced firing (Extended Data Fig. 11b and c). Also, removing intracellular Ca^2+^ by including BAPTA in the patch pipette only slightly reduced the firing frequency of acclimated VMPO^LepR^ neurons (Extended Data Fig. 11d). Similarly, neither nifedipine nor mibefradil, blockers of L- and T-type voltage-gated Ca^2+^ (Ca_V_) channels, majorly affected the firing frequency of acclimated VMPO^LepR^ (Extended Data Fig. 11e), and we rather observed the overall Ca_V_-mediated current to be larger in *non*-acclimated VMPO^LepR^ compared to acclimated cells (Extended Data Fig. 11f), arguing for a minor role of Ca_V_s, if any, in VMPO^LepR^ pacemaking.

Voltage-gated Na^+^ (Na_v_) channels are at the core of AP initiation and upstroke. We therefore tested whether changes in Na_v_s could explain acclimation-induced spiking. We found the kinetic parameters of APs, such as the AP rise time and half-width, to be more rapid in acclimated VMPO^LepR^ compared to controls (Fig. 5d and Extended Data Fig. 11a) and transient Na_v_ currents to be of larger amplitude (Fig. 5e), suggesting that acclimation had changed Na_v_ composition and functionality to support faster firing.

Persistent (I_NaP_) and resurgent (I_NaR_) Na^+^ currents, both of which are carried by Na_v_s, have been associated with higher excitability and tonic pacemaker activity in several different central and peripheral neuronal populations^54–58^. Specific molecular rearrangements and the presence of certain auxiliary subunits permit TTX-sensitive Na_v_ channels to inject depolarizing currents during interspike intervals to drive neurons to threshold voltages, thereby inducing repetitive firing. We found I_NaP_ and I_NaR_ in acclimated VMPO^LepR^ to be significantly larger than in non-acclimated controls (Fig. 5f and Extended Data Fig. 12a). Riluzole is a compound that preferentially blocks I_NaP_ and I_NaR_ but, unlike TTX, does not inhibit the transient Na_v_ current at low concentrations^59,60^. Indeed, Riluzole inhibited TTX-sensitive I_NaP_ and I_NaR_ present in acclimated VMPO^LepR^; in contrast, the compound only had minimal effects on non-acclimated neurons (Fig 5g and Extended Data Fig. 12b, c). In agreement with its reported selectivity for I_NaP_ and I_NaR60_, Riluzole, at the concentration used, did not reduce transient Na_v_ currents (Extended Data Fig. 12d). Importantly, Riluzole significantly reduced tonic AP firing in acclimated VMPO^LepR^ but had no significant effect on slowly firing non-acclimated neurons (Fig. 5h and Extended Data Fig. 12e), demonstrating that I_NaP_ and/or I_NaR_ contribute majorly to acclimation-induced pacemaking.

Among the different TTX-sensitive Na_v_s that could generate I_NaP_ and I_NaR_, we found 5 out of the 6 corresponding alpha subunits, Na_v_1.1-1.3 and 1.6-1.7, to be expressed in VMPO^LepR^ (Extended Data Fig. 12f). Pharmacological profiling, using semi-selective inhibitors targeting Na_V_1.7 (ProToxin-II and PF-05089771), Na_V_1.6 (4,9-Anhydro-tetrodotoxin, a TTX derivative that displays some cross-inhibitory potential on Na_V_1.1^61^) and Na_V_1.2 (Phrixotoxin-3), ruled out these channels as major contributors to I_NaP_ (Extended Data Fig. 12g) and I_NaR_ (data not shown) that we observed in acclimated VMPO^LepR^. Na_V_1.7 has been implicated in plastic changes of hypothalamic neurons^62^. However, Na_V_1.7 inhibition did not significantly affect Na_V_ currents or acclimation-induced AP firing (Extended Data Fig. 12g, h). Additionally, RNA knock-down specifically in VMPO^LepR^ neurons using LepR-Cre mice in combination with previously published viral AAV-shRNA particles targeting Na_V_1.7^62^, also did not affect tonic firing of acclimated VMPO^LepR^ (Extended Data Fig. 12i). Only ICA-121431, an antagonist of Na_v_1.3 and Na_v_1.1 channels^63^ substantially reduced I_NaP_ in VMPO^LepR^ (Fig. 5i). Because Na_v_1.1 is also inhibited by 4,9-Anhydro- tetrodotoxin^61^, an antagonist that did not show any effect on I_NaP_ (Extended Data Fig. 12g), we concluded that Na_v_1.3 is the more likely candidate of the two Na_v_ subtypes inhibited by ICA- 121431 and relevant for generating acclimation-induced I_NaP_.

The effect of ICA-121431 on I_NaP_ was similar (and non-additive) to that observed for Riluzole (Fig. 5i). ICA-121431 also reduced tonic AP firing of VMPO^LepR^ (Fig. 5h) and, as opposed to Riluzole, had a negligible effect on I_NaR_ (Extended Data Fig. 12b). Importantly, both Riluzole and the Na_V_1.3 blocker robustly reduced acclimation-induced warm-sensitivity of VMPO^LepR^ (Fig. 5j, k).

Collectively, these pharmacological experiments suggest that Na_V_1.3 driven I_NaP_, but not I_NaR_, is a major contributor to VMPO^LepR^ warm-sensitive pacemaking.

### Na_V_1.3 drives tonic warm-sensitive activity of VMPO^LepR^ and contributes to heat acclimation

To further investigate the role of Na_v_1.3 in VMPO^LepR^ activity and heat tolerance, we used an RNA-interference-mediated “knock-down” strategy, similar to that used for Na_V_1.7 above and we developed AAV vectors for Cre-dependent, cell-type-specific knock-down of Na_V_1.3 in VMPO^LepR^ (Fig. 6a and b). We confirmed Na_V_1.3 knock-down by qPCR (Extended Data Fig. 13a). Indeed, the amplitude of I_NaP_ was reduced in acclimated VMPO^LepR^ neurons when Na_v_1.3 was knocked-down, but not when scrambled control shRNA was used (Fig. 6c). Moreover, warm- sensitivity was also strongly and significantly reduced by Na_v_1.3 knock-down, while baseline excitability of non-acclimated neurons was not affected (Fig. 6d, e and Extended Data Fig. 13b- e), further strengthening the association between the I_NaP_ and acclimation-induced VMPO^LepR^ firing properties.

**Fig. 6.**
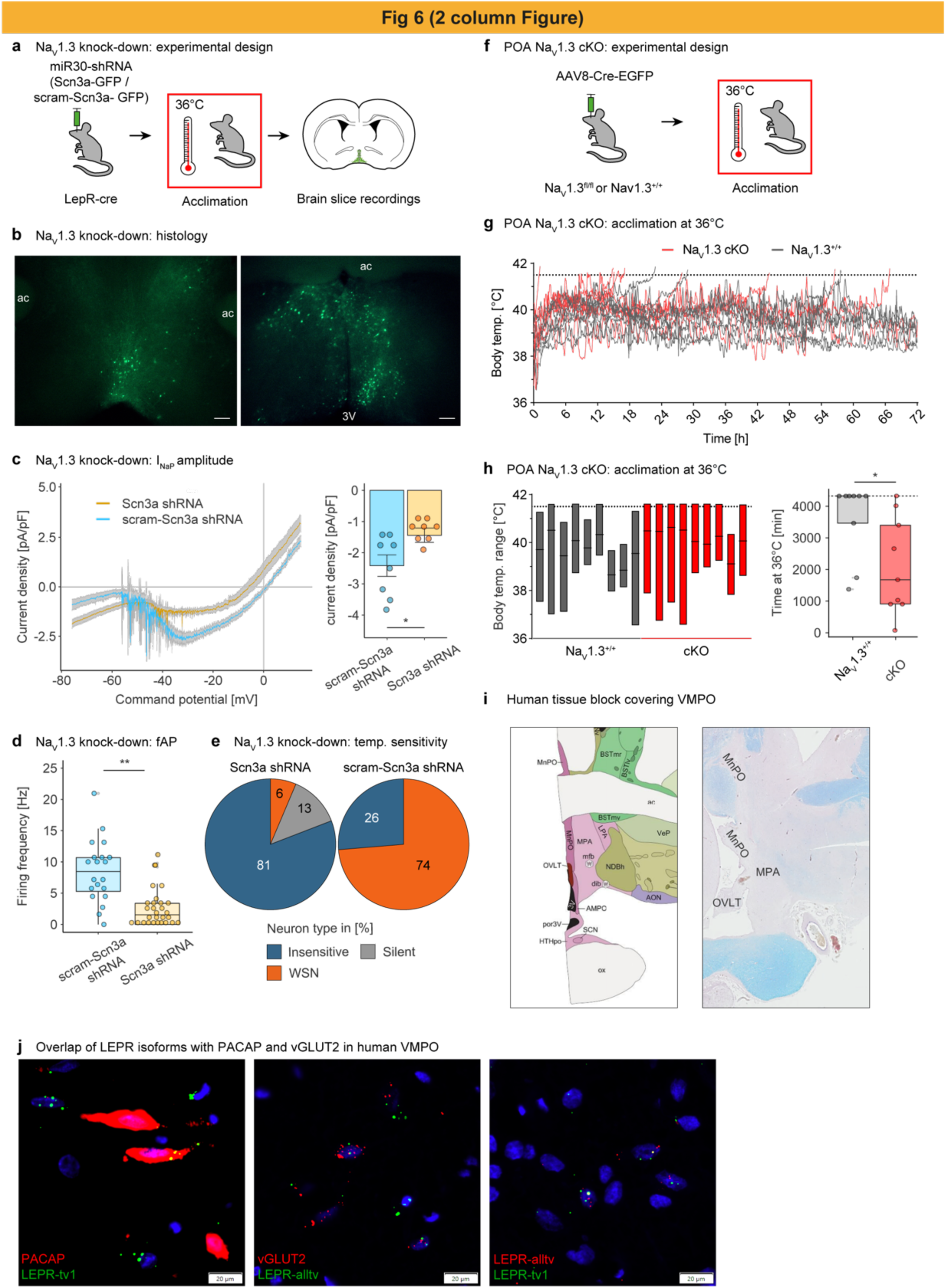
NaV1.3 is required for acclimation-induced tonic warm-sensitive firing and heat tolerance. a, Schematic showing the experimental protocol for NaV1.3 mRNA knock-down in LepR-Cre mice. The animals were POA-injected with the mix of AAVs carrying Cre-dependent constructs encoding shRNAs against Scn3a or scrambled control. A minimum of 3 weeks was allowed for recovery and viral DNA expression after which animals were acclimated at 36°C (≥ 4weeks) followed by brain slice recordings. b, Representative image of acclimated VMPO^LepR^ neurons labeled with GFP encoded within the shRNA constructs; scale bar: 250 μm. Note that shRNA-GFP expression is Cre-dependent and thus only the small population of VMPO^LepR^ neurons are infected. c, Left: Average INaP in VMPO^LepR^ neurons expressing functional shRNAs against NaV1.3 mRNA (Scn3a shRNA) and a scrambled control shRNA (scram-Scn3a shRNA). Traces presented as mean ± s.e.m. Right: Quantification (mean ± s.e.m.) of INaP at -35mV based on data shown left, showing a reduction of INaP in Scn3a shRNA expressing acclimated LepR+ neurons. Unpaired two-tailed t-test, *P = 0.0174. n = 8/4 (Scn3a shRNA) and n = 8/3 (scram-Scn3a shRNA) cells. d, Firing frequency of acclimated VMPO^LepR^ neurons expressing either Scn3a shRNAs or scram-Scn3a shRNA (data presented as median and interquartile range). Frequency of spontaneous APs was significantly reduced by the functional shRNAs. Unpaired two-tailed t-test, **P = 0.0044. n = 30/5 (Scn3a shRNA) and n = 20/3 (scram-Scn3a shRNA) cells. e, Distribution of temperature-insensitive, cold-sensitive (CSN), warm-sensitive (WSN) and silent neurons within the acclimated VMPO^LepR^ neurons expressing either Scn3a shRNAs or scram-Scn3a shRNA. n = 47/5 (Scn3a shRNA) and n = 19/3 (scram-Scn3a shRNA) cells. f, Schematic showing the experimental protocol for conditional NaV1.3 knock-out (NaV1.3 cKO). NaV1.3^fl/fl^ and WT controls were injected with an AAV encoding the Cre recombinase under the control of a general CAG promoter into the POA. After 4 weeks of recovery and viral DNA expression, animals were at 36°C (≥ 4weeks) and Tcore was recorded telemetrically. g, Body temperature traces of individual NaV1.3 cKO and WT control animals (N = 9 each) during heat acclimation. Most of the WT animals were able to defend their body temperature within physiological range. In contrast, all but one of the NaV1.3 cKO animals were unable to maintain their body temperature below 41.5 °C. Acclimation was discontinued after 72 hours. h, Left: Range of body temperatures of each animal shown in (g). Horizontal lines represent median body temperatures. Animals were discontinued from acclimation chamber as soon as they reached 41.5 °C. Right: quantification of endurance time at 36°C acclimation temperature of mice shown (g). Cut-off time: 72 hr (dashed line). Mann Whitney test, *P = 0.0155 (N = 9 each). Data presented as median and interquartile range. i, Left: Allen Brain Atlas annotation of human preoptic areas. Right: Human tissue block covering preoptic areas MnPO/MPA/OVLT (LFB/HE stain). j, LEPR co-expression in human VMPO with RNAscope ISH. Left: PACAP + LEPR-tv1 (long isoform; co-expression in yellow). Middle: vGLUT2 + LEPR-alltv (all isoforms; co-expression in yellow). Right: LEPR-alltv + LEPR-tv1 (co-expression in yellow). Neuronal activity and currents were recoded under fast synaptic transmission blockade and at 36°C (see also Extended Data Figure 13-16).

In order to assess the role of Na_V_1.3’s enhanced functionality in acclimation and heat tolerance *in vivo*, we employed Na_V_1.3^fl/fl^ mice to conditionally delete the Scn3a gene^64^. Homozygous floxed mice and wildtype controls were preoptically injected with Cre recombinase-encoding AAVs (Fig. 6f and Extended data Fig 14a). After verification of Cre-mediated Scn3a gene deletion in the POA (Extended data Fig. 15a, b) we first assessed baseline T_core_ of the animals at room temperature (23°C) and found it to be indistinguishable between the conditional POA-Na_V_1.3^-/-^ mice and the control group (Extended Data Fig. 15c). Strikingly, all but one of the POA-conditional Na_v_1.3 knock-out mice failed already during heat acclimation at 36°C and the animals reached the critical cut-off T_core_ of 41.5°C approximately twice as fast as AAV-Cre injected wildtype controls (Fig. 6g, h). It appeared that AAV-Cre infection, broadly covering the mouse POA, had some effect in wildtype animals, slightly hampering their ability to acclimate, albeit to a lesser extent than in mice lacking the Na_V_1.3 channel. These data provide further genetic evidence that preoptic Na_V_1.3 plays an important role in heat acclimation and tolerance.

Given the critical role of VMPO^LeprR^ in heat acclimation and heat tolerance in mice we wondered whether this population of neurons, that are part of so-called “QPLOT” neurons mediating heat loss responses^9^, also exists in humans. To this end, we carried out multiplex in situ hybridizations (RNAscope) on postmortem human brain tissue encompassing the human VMPO (based on anatomical annotations from the adult human Allen Brain Atlas of MnPO, <u>m</u>edial <u>p</u>reoptic <u>a</u>rea (MPA) and the closely neighboring <u>o</u>rganum <u>v</u>asculosum laminae terminalis (OVLT) region - https://atlas.brain-map.org). Indeed, we were able to detect a subset of neurons within the human VMPO to express overlapping “QPLOT” marker genes including the leptin receptor, PACAP, OPN5 and PTGER3 (Fig. 6i, j and Extended Data Fig. 16).

In summary, our work emphasizes an acclimation-induced plasticity mechanism involving a persistent Na^+^ current ––carried largely by Na_V_1.3–– that drives warm-sensitive AP firing in VMPO^LepR^ to promote heat tolerance.

## Discussion

Intrinsic warm-sensitivity has been used as a defining functional parameter for a subset of POA neurons for many decades^65,66^. To what extent this feature is physiologically relevant has been a matter of debate^16^. Recent studies suggests that in mice living under normal ––coolish–– housing conditions^67^, experimental POA heating has only a small effect on body temperature regulation^27,30^, arguing for modest relevance of POA heat sensitivity in rodents under these conditions. Here we show that long-term heat acclimation plastically transforms VMPO neurons to become spontaneously active, highly warm-sensitive neurons. Their gained activity is critically important to drive heat tolerance and to trigger heat loss responses in hot environments.

It is interesting to note that, *in vivo*, the increase in acclimation-induced warm responsiveness is recruited from the cold-responsive (CRN) population, whereas in brain slices, it is recruited from temperature-insensitive neurons. Cold sensitivity has been suggested to largely stem from synaptic connectivity rather than constituting a neuron-intrinsic property^26^. Given the high degree of reciprocal (local and long-range) POA connectivity, ––which is largely absent in *ex vivo* slice preparations–– it is therefore conceivable that heat acclimation additionally induces changes in the strength of synaptic connections, thereby contributing to the robust induction of warm-responsive neurons from the pool of CRNs.

Heat acclimation modulates energy metabolism and promotes loss of body weight^33^. On the other hand, perturbed energy metabolism and obesity negatively affect heat acclimation^68^. Leptin, a major signaling indicator of energy metabolism status, has been implicated in thermoregulation and body temperature increases^41,69,70^. Parallel to a reduction in body weight, we observed a drop in leptin levels in heat acclimated animals. In line with a subtle role of leptin to modulate heat acclimation, we found that leptin supplementation during heat acclimation slightly reduced AP firing frequencies of VMPO^LepR^. However, leptin supplementation neither affected VMPO^LepR^ warm-sensitivity nor the animals’ performance in the heat endurance assay. This, along with our *in vivo* imaging data, indicates that the acquired warm-sensitivity––rather than a general tonic activity increase–– likely mediates enhanced heat tolerance in acclimated animals upon a heat challenge.

Our study suggests the LPBN, which processes and relays peripheral temperature information to thermoregulatory POA neurons^13,14,71^, to be critical at the beginning of heat acclimation but to be dispensable at later stages. These two phases ––Phase 1: thermo-afferent/LPBN driven; Phase II: driven by spontaneous, temperature-sensitive VMPO^LepR^ activity–– may very well coincide with the two phases of heat acclimation that have been described based on transcriptional profiling studies in rodents^31^.

After heat acclimation, chemogenetic inhibition of VMPO^LepR^ had a robust effect and dramatically reduced heat tolerance, which contrasts with the effect observed when inhibiting LPBN (Fig. 2e, f and Extended Data Fig. 8d). These data suggest that the low activity of VMPO^LepR^ prior to heat acclimation is not majorly contributing to baseline heat tolerance, a task possibly performed by other/parallel thermo-afferent pathways. However, upon heat acclimation, VMPO^LepR^ gain dominance and peripheral thermo-afferent pathways have a reduced influence on heat tolerance.

This peripheral→central shift in thermoregulatory control can be explained by a transfer of intrinsic warm-sensitivity to VMPO^LepR^ upon heat acclimation, that we not only find in *ex vivo* brain slice recordings but also *in vivo*. Given that (i) synaptic blockers don’t affect warm- sensitivity in slice recordings, (ii) inhibition of LPBN→POA thermo-afferent pathways appear to have little impact subsequent to heat acclimation and (iii) considering that an elevation in T_core_ caused by environmental heat challenges is directly detectable in the POA^27,30^, suggests that VMPO^LepR^ gain intrinsic cell-autonomous warm-sensitivity that is independent of thermo-afferent synaptic drive. We presume that the newly gained warm-sensitivity allows robust detection of T_core_ and permits VMPO^LepR^ to keep T_core_ in check at dangerously high ambient temperatures. This transformation may reduce VMPO receptivity to peripheral ––anticipatory^16^–– heat detection, which presumably is of lesser importance when ambient temperatures are permanently high and affect T_core_.

We hypothesize that VMPO^LepR^ tonic warm-sensitivity, which we find to develop over many days during heat acclimation, drives the adaptation of peripheral organs involved in thermoregulation. Whether acclimation-relevant parameters of multiple organs ––including heart rate/weight, basal metabolic rate (BMR), brown adipose tissue (BAT) activity/capacity, cutaneous heat dissipation/insulation, water balance and others^33–35^–– are orchestrated by VMPO^LepR^ activity or whether only a subset of adaptations are VMPO^LepR^-dependent is currently unknown.

The ion channels TRPM2 and TRPC4 have previously been proposed to detect acute temperature changes in the POA^27,30,48^. However, pharmacologic blockade of either ion channel did not significantly inhibit spontaneous warm-sensitive AP firing of acclimated VMPO^LepR^. Because TRPM2 was shown to constitute a molecular temperature sensor in presynaptic terminals we did not expect the channel to drive cell-autonomous warm-sensitive AP firing in VMPO^LepR^. It is possible that acclimation induces a warm-sensitivity mechanism in VMPO^LepR^ that is distinct from the molecular mechanism(s) utilized by canonical WSNs.

We implicate background and voltage-gated Na^+^ currents in the acclimation-induced tonic pacemaker activity and warm-sensitivity. Induced pacemaker activity is a feature that is similar to hypothalamic neurons residing in the suprachiasmatic nucleus (SCN) important for circadian clock function: during daytime of the circadian cycle, background and voltage-gated Na^+^ currents are induced to increase tonic AP firing of SCN neurons relevant for circadian homeostasis^53,72^. Intriguingly, AP firing of some SCN neurons is also temperature sensitive^73^.

Genetic perturbation as well as pharmacologic inhibition of Na_V_1.3 not only reduced the tonic activity of acclimated VMPO^LepR^ neurons but also abrogated their warm-sensitivity, suggesting that the two features are mechanistically linked. However, it is unclear whether Na_V_1.3 conveys warm-sensitivity directly or whether another molecular mechanism modulates its pacemaker activity in a temperature dependent manner. Na_V_1.3 appears to be present in murine POA already at baseline conditions and its mRNA level does not seem to change following heat acclimation (Extended Data Fig. 12f and 15b). It is thus likely that Na_V_1.3’s interplay with additional channels and/or auxiliary channel subunits may render the neurons spontaneously active and warm-sensitive. Such a scenario would be similar to Na_V_1.7’s reported interaction with FGF13 that results in increased heat sensitivity in peripheral sensory neurons^74^.

Possibly, Na_V_1.3 is already required by preoptic neurons to transmit temperature information early on during acclimation, potentially explaining why mice lacking Na_V_1.3 in preoptic neurons already fail during the early phase of acclimation.

While our study emphasizes the role of Na_V_1.3 for acclimation-induced activity in VMPO neurons, it is important to note that the coordinated action of several different classes of channels is likely necessary to fully express the highly regular, warm-sensitive AP firing rate increases observed in acclimated VMPO^LepR^, akin to the electrical interaction of multiple conductances in SCN pacemaker neurons^75^.

Our findings provide a basic molecular and cellular framework governing the central regulation of heat acclimation (Fig. 7). We anticipate that this work will pave the way to further elucidate how homeostatic pathways adapt rheostatically, and whether the underlying plasticity can be utilized in medical settings, such as enhancing tolerance to hot environmental conditions.

**Fig. 7.**
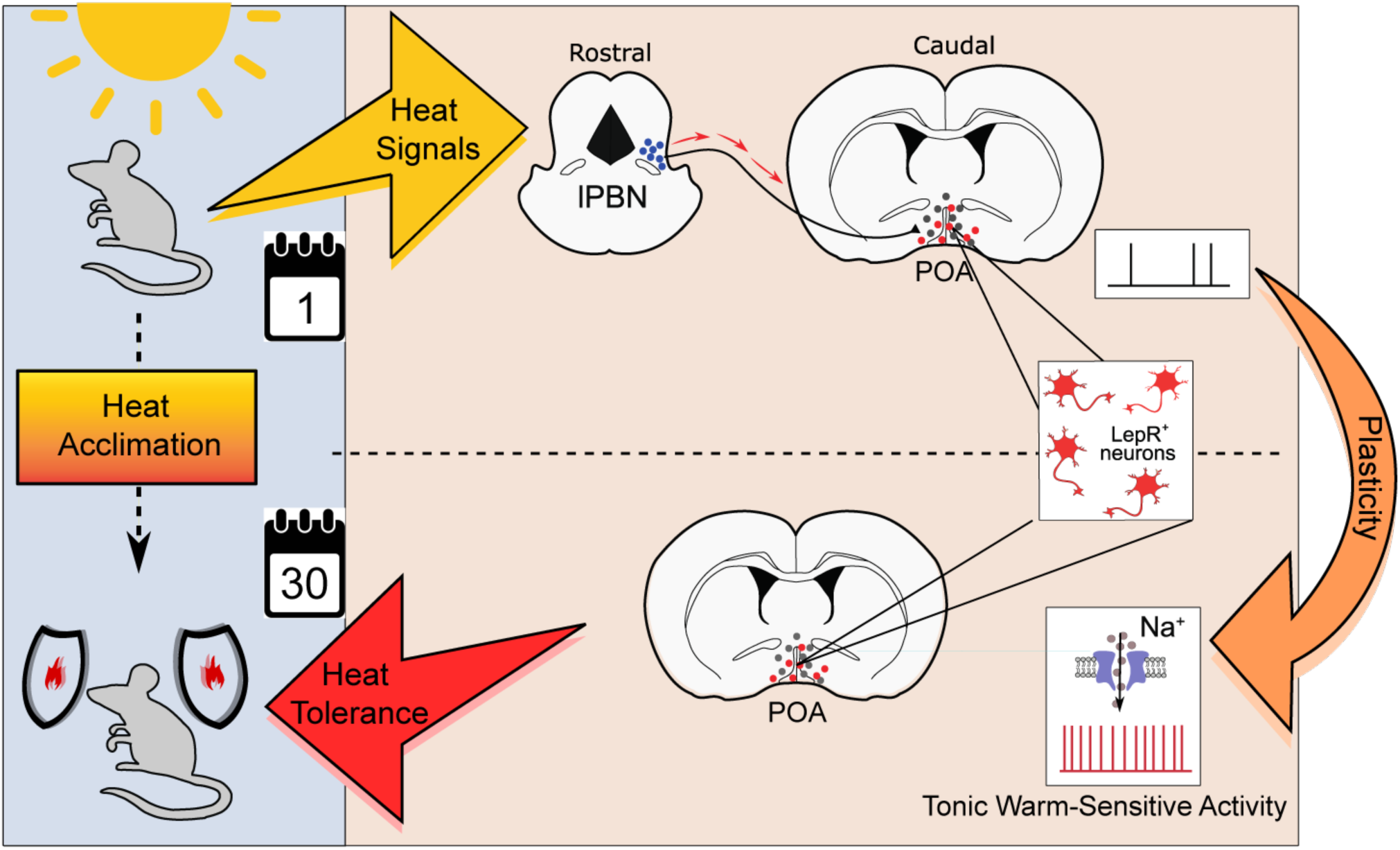
Summary Heat stimuli reach thermoregulatory neurons in the hypothalamic preoptic area (POA) via parabrachial thermo-afferent pathways (LPBN). Sustained, long-term heat exposure triggers an adaptive process that transforms LepR-expressing POA neurons to become tonically active and warm-sensitive. This form of cellular plasticity, which is mediated in part by the activity of a voltage-gated sodium channel, increases heat tolerance in mice to protect the animals from the detrimental effects of hot environments.

## Acknowledgements

We thank Amandine Cavaroc, Daniela Pimonov and Daniel Erdelyi for technical support; members of the Siemens lab for inspiring discussions and critical input; Silvia Calderazzo for help with statistical analysis of RNAseq data; Christine Fischer for help with statistical analysis of *in vivo* physiology experiments; Scott Sternson for making available AAV-shRNA particles targeting Na_V_1.7; John Wood for providing Na_V_1.3^flox/flox^ mice; Jeanette Bannebjerg Johansen, from Novo Nordisk A/S for technical support on human brain histology and The Edinburgh Brain and Tissue Bank for providing human brain tissue samples.

Additionally, the authors thank Angelika Lampert, Henning Fenselau, James Poulet and Christopher Bohlen for discussions, scientific input and valuable criticism of the manuscript; The Nikon Imaging Center at Heidelberg University for support with confocal microscopy. Funding: The authors gratefully acknowledge the data storage service SDS@hd supported by the Ministry of Science, Research and the Arts Baden-Württemberg (MWK) and the German Research Foundation (DFG) through grant INST 35/1314-1 FUGG. This work was supported by the European Research Council ERC-CoG-772395, the German research Foundation SFB/TRR 152 (to J.S.), and SFB1158 (to J.S. and C.A.), the International Human Frontier Science Program Organization postdoctoral fellowship LT000762/2019-L (to W.A.) and the European Molecular Biology Organization (EMBO) postdoctoral fellowship (to S.N.), and the Chica and Heinz Schaller Foundation, NARSAD, the Fritz Thyssen Foundation (to C.A.).

## Author Contributions

J.S. together with S.N., W.A., and J.P conceived the project. J.P. discovered the cellular VMPO^LepR^ neuron phenotype and initiated its characterization together with S.N. K.Z. performed and analyzed RNA_Seq_ data of FACS sorted VMPO^LepR^ neurons and carried out histological stainings, including cFos labeling. S.N. with help from C.A.S. carried out most *in vivo* manipulations including chemogenetic and optogenetic manipulation of VMPO^LepR^ activity and established the heat endurance assay. GP, GM, and CA designed, performed and analyzed *in vivo* Miniscope imaging. S.L. performed mRNA transcript analysis of human hypothalamic brain sections. L.I.L.G. performed qPCRs of Na_V_1.3 knock-out and knock-down mice. Detailed electrophysiological characterization of VMPO neuronal activity and its underlying ionic currents, including pharmacologic and genetic Na_v_1.3 manipulations, were performed by W.A. Python code for electrophysiology data analysis was provided by M.H. J.S. together with S.N. and W.A. wrote the paper. All authors commented on and approved the paper.

## Supplemental information

Document S1. Extended Data Figures 1-16, including corresponding legends.

Supplementary Video 1, Title: *In vivo* GCaMP6 imaging of VMPO^LepR^ neurons before heat acclimation. Related to Figure 1. Representative LepR-Cre animal injected to express GCaMP6 in the VMPO. The mouse was not heat-acclimated but kept at normal housing conditions (22°C-23°C). During the GCaMP6 signal recording the mouse was first kept at a temperature of 22°C and then exposed to 36°C as indicated in the upper part of the video. Blue ovals indicate registered cells that respond to the heat stimulus.

Supplementary Video 2, Title: *In vivo* GCaMP6 imaging of VMPO^LepR^ neurons after heat acclimation. Related to Figure 1 The same animal as shown in Supplementary Video 1 but after 30 days of heat acclimation at 36°C. Shortly before and during the first part of the GCaMP signal recording session the mouse was first exposed to a temperature of 22°C and then exposed to 36°C as indicated in the upper part of the video. Blue ovals indicate registered cells that had also responded to the heat stimulus before heat acclimation (as shown in Supplementary Video 1); red ovals indicate additional cells that responded to the heat stimulus only after heat acclimation and that were not detectable in the non-acclimated mouse.

## Methods

### Lead contact

Further information and requests for resources and reagents should be directed to and will be fulfilled by the Lead Contact, Jan Siemens.

### Mice

The following mouse lines were used in this study: LepR-cre (B6.129-Leprtm3(cre)Mgmj/J; The Jackson Laboratory, IMSR Cat# JAX:032457), PACAP-EGFP (Tg(Adcyap1- EGFP)FB22Gsat/Mmucd; MGI ID: MGI:4846839)^76^, Rosa26Lox-stop-LoxHTB (The Salk Institute for Biological Studies)^77^, Vglut2-cre (Slc17a6tm2(cre)Lowl/J; The Jackson Laboratory, IMSR Cat# JAX:016963), Vgat-cre (Slc32a1tm2(cre)Lowl/J; The Jackson Laboratory, IMSR Cat# JAX: 016962), TrpV1-cre (B6.129-Trpv1tm1(cre)Bbm/J; The Jackson Laboratory, IMSR Cat# JAX:017769), Rosa_DTA (Gt(ROSA)26Sortm1(DTA)Jpmb/J; The Jackson Laboratory, IMSR Cat# JAX:006331), FosTRAP2 (Fos2A-iCreER/+(FosTRAP2); The Jackson Laboratory, IMSR Cat# JAX:030323), Nav1.3-floxed (B6.129S6-Scn3atm1.1Jwo/H; EMMA Strain ID: EM:02214). Heterozygous mice were used for experiments with the exception of Nav1.3-floxed line where homozygous mice were used to create a conditional knock-out.

All animal experiments were in accordance with the local ethics committee and governing body (Regierungspräsidium Karlsruhe, Germany) and were approved under protocol numbers: G- 111/14, G-168/15, G-169/18, G-223/18 and G-181/21. Mice were housed at room temperature (RT, 23 ± 1°C, unless specified otherwise) in air-conditioned lab space / animal vivarium with a standard 12-h light/dark cycle and *ad libitum* access to food and water. All genetically modified mice in this study were on the C57BL/6N background. All studies employed a mixture of male and female mice.

### Acclimation protocol

Mice were divided according to their acclimation status. Acclimated animals were attained by continuous exposure to 36 ± 1°C for either 24 hrs, 4 days or ≥4 weeks, at a humidity level of 45 ± 5%. Full heat acclimation in rodents is reached after around 4 weeks of habituating the animals to warm temperatures^19^. We therefore generally ––and if not stated otherwise–– used mice acclimated for 4 to 5 weeks, which in the manuscript we denote as “≥ 4 weeks”. Mice held at RT served as control (non-acclimated) group. For heat acclimation, mice were placed in a climate chamber (Binder, KB720) with *ad libitum* access to food and water. All mice were kept at the standard 12- h light/dark cycle. Mice of age between 7 and 14 weeks were used for heat acclimation.

### Heat endurance assay

Previous studies on the dynamics of acclimation reported that the acclimatory homeostasis is reached after 25-30 days whereas short-term acclimation occurs after 2-3 days of acclimation^36^. At the end of acclimation period, animals were evaluated in a heat endurance assay. The heat endurance assay took place in a similar climate chamber as used for acclimation where the ambient temperature was set to 39°C ± 0.5°C. Animals (11-16 weeks old following full acclimation) were transferred immediately from acclimation chamber to the 39°C chamber (always in the morning, between 9 AM and 11 AM) where they took part in heat endurance assay lasting for up to a maximum of 24 h. Similar to the acclimation period, mice had access to food and water *ad libitum*.

Body temperature of the mice was constantly monitored for the entire period. A body temperature of 41.5°C was used as cut-off criterion^37,78^. At the end of the heat endurance test, animals were shortly placed back to 36°C in order to avoid prolonged hypothermia and monitored until the animals were sacrificed. Mice were tested only once in the heat endurance assay and not multiple times.

In experiments where mice where supplemented with Leptin during heat acclimation (and prior to heat endurance assay), animals were administered with Leptin (Peprotech, Cat# 450-31, diluted in phosphate buffer solution) at a dose of 1.25 mg/kg twice daily i.p.^79^

### General immunohistochemistry procedures

Animals were deeply anaesthetised with isoflurane and transcardially perfused with a phosphate buffer solution (PBS; 3.85 g of NaOH and 16.83 g of NaH_2_PO_4_ in 1L of distilled water) followed by a 4% paraformaldehyde (PFA) solution. Brains were dissected out and left overnight (O/N) in 4% PFA at 4 °C. Over the following 2 days brains were immersed into PBS/sucrose solutions (24 hours in 10% sucrose followed by 30% sucrose, until brains sank to the bottom of container tube). Brains were sectioned with a cryo-microtome at 30 μm thickness and sections (free-floating) were kept in cryo-protectant solution (250ml Glycerol; 250ml ethylene glycol filled up with PBS to 1 L) at 4 °C until stained.

For antibody staining (see primary antibody list below), sections were washed once in PBS and left overnight at 4 °C in 0.2% TritonX-100 (PBX0.2). On the following day, sections were blocked with 5% goat serum in PBS containing 0.1% TritonX-100 (PBX0.1) for 2 hours at RT. Sections were then incubated with primary antibodies, diluted in 1% goat serum in PBX0.1, for 3 days at 4°C. On the fifth day, sections were washed extensively with PBX0.1 and were then incubated with secondary antibodies and DAPI for 4 hours at RT. Finally, tissue was washed extensively with PBSX0.1 and once with PBS after which sections were mounted using Immu-Mount (Cat# 9990402, Fisher Scientific, UK) onto glass slides.

Confical images were taken at the Nikon imaging center at Heidelberg University, with the Nikon A1R confocal microscope under Nikon Plan Apo λ 10x magnification NA 0.45 (working distance 4mm, the field of view 1.27 x 1.27 mm) objective. Cell counting of cells expressing markers of interest was performed with NIS-Elements software (Nikon Instruments, Inc) using an automatic cell counting method. The same thresholding of fluorescence signal was used for each of the color channels in all the quantified images. Images presented were processed with ImageJ.

### Antibodies

- chicken anti-GFP (1:1000, Novus Biotechne Cat# NB100-1614)
- rabbit anti-c-Fos (1:1000, Synaptic Systems Cat# 226 003)
- rabbit anti-mCherry (1:1000, Abcam Cat# ab167453)
- rabbit anti-SCN3A (1:700, Abcam Cat# ab65164)
- goat anti-chicken Alexa Fluor 488 (1:750, Thermo Fischer, Cat# A-11039),
- goat anti-rabbit Alexa Fluor 555 (1:750, Thermo Fischer, Cat# A-21430
- DAPI (1:10000, Sigma, Cat# 10236276001)

### TRAPping of WRNs using FosTrap2 mice

“TRAPping” of warm-responsive neurons (WRNs): heterozygous FosTRAP2;HTB mice (resulting from crossing FosTRAP2 mice with the Rosa26Lox-stop-LoxHTB reporter line) mice were habituated in their home cages in a climate chamber (Binder, KB720) at 23°C and injected with saline solution on 5 consecutive days to reduce stress responses. On the day of the experiment the climate chamber was warmed to 36°C; 2 hours into warmth exposure 4-OHT (see preparation procedure below) was delivered by i.p. injection at a dose of 50 mg/kg. Mice were kept at 36°C for another 2 or 6 hours to reach a total of 4h and 8h “TRAPping” duration, respectively. Control FosTRAP2;HTB mice kept at RT (and not warmed to 36°C) were treated in the same way (5 consecutive days of saline injections before 4-OHT injection). After the corresponding warmth exposure, both groups of animals were left at thermoneutrality (31°C) for 48 hours in order to prevent secondary trapping of cold responsive cells and expecting the 4-OHT to be completely metabolized. For electrophysiology, mice were subsequently either kept at RT or acclimated at 36°C.

Drug preparation: z-4-hydroxytamoxifen (4-OHT, Sigma H7904) was prepared for intraperitoneal (i.p.) delivery essentially as described before^80^ with some modifications: 4-OHT was dissolved at 20 mg/mL in ethanol by vigorous shaking at RT for 5 min plus 1 min of sonication in a bath sonicator and was then aliquoted in 50 microliter (1 mg) aliquots and stored at –80°C for up to several weeks. Before use, 4-OHT was redissolved by vigorous shaking at RT for 5 min plus 1 min of sonication in a bath sonicator; subsequently, 200 microliter of 1:4 mixture of castor oil:sunflower seed oil (Sigma, Cat #s 259853 and S5007) was added per 50 microliter aliquot containing 1mg of 4-OHT and the Ethanol-oil suspension was vigorously mixed; subsequently the ethanol was evaporated by vacuum under centrifugation (without heating). The final 5 mg/mL 4- OHT solution was always used on the day of preparation. All injections were delivered i.p.

For immunohistochemistry, animals following warmth exposure were left at 31°C until the following day. After this, all three groups (4h “TRAPped”, 7h “TRAPped”, and control group were transferred to their home cages for the next 2.5 weeks. After this period all mice were placed in a climate chamber (Binder, KB720) for 24 hours at 23°C to get accustomed once more to the chamber’s environment. On the following day, the temperature in the climate chamber was adjusted to reach 36°C to perform the classical warming challenge for hours. After the 4-hour exposure to warmth, animals were sacrificed using isoflurane and transcardially perfused. POA- containing brain sections were cut at 30 μm thickness as described above. Tissue was stained for GFP and cFos in order to quantify the overlap of the TRAP-positive neurons (HTB/GFP positive) with neurons expressing endogenous cFos after the 36°C warming stimulus.

### cFos expression in VMPO^LepR^ neurons after exposure to 36°C ambient temperature

To elucidate the role of VMPO^LepR^ neurons in thermoregulatory responses, we investigated whether LepR+ neurons are activated by acute warmth exposure. To do this, LepR-cre mice crossed to the Rosa26Lox-stop-LoxHTB reporter line^81^ (here referred to as LepR-Cre;HTB mice) got accustomed to the climate chamber for 24 hours. On the 2^nd^ day control animals were taken out, anaesthetized with isoflurane and transcardially perfused with PBS followed by 4% PFA. The temperature of the climate chamber was switched to 36°C and the experimental animals were kept at this temperature for 4h, immediately followed by anaesthesia and perfusion. Brains were dissected out and left O/N in PFA at 4°C. Brains were immersed in sucrose solutions and sliced as described above. α-GFP and α-cFos primary antibodies were applied to amplify HTB/GFP reporter and label endogenous cFos proteins.

### Nav1.3 staining

C57BL/6 and Nav1.3^flox/flox^ mice were injected with AAV-Cre-GFP. After 4 weeks, to allow AAV expression and protein turnover, animals were transcardially perfused with PFA and brain tissue was processed for immunohistochemistry as described above. 30 μm free-floating brain sections containing POA and cortex were stained with primary antibodies against Na_V_1.3 and GFP.

### Constructs for Scn3a knockdown

Short-hairpin (sh) RNA constructs for Scn3a were developed according to the method described in Matveeva et al.^82^, with the murine Scn3a canonical cDNA sequence as template. AAV2-based CAG::FLEX-rev-hrGFP:mir30(Scn9a) vector, used previously by Branco et al.^62^, was used as backbone following the excision of the shRNA sequence targeting Nav1.7 using EcoRI and XhoI restrictases (New England Biolabs). Using the miR_Scan tool (https://www.ncbi.nlm.nih.gov/staff/ogurtsov/projects/mi30/), we have selected three sequences, binding to the 5’ region (encoding the extracellular loop between segments 5 and 6 of domain I of the channel; sense strand sequence: GAAGGACTATATCGCAGATGA), a central region (encoding the intracellular loop connecting domains II and III; sense strand sequence: GTGGAGAAATACGTAATTGAT) and the 3’ region (encoding the segment 2 of domain IV; sense strand sequence: GTCCCGAATCAACCTGGTATTT), to construct shRNAs against. Sense strands and guide strands, separated by the loop sequence TACATCTGTGGCTTCACTA and supplemented with restriction site overhangs, were synthesized as oligonucleotides and together with complementary oligonucleotides aligned and cloned into the recipient vector. Such AAV vectors, where the shRNA sequences were placed between a FLEX switch sequence together with a GFP reporter gene, were packaged into AAV1/2 particles by the Viral Vector Facility,University of Zurich (Zurich, Switzerland).

As a negative control for these shRNA Scn3a shRNA constructs, we produced an AAV containing a scrambled sequence (ACTGTAGTCGTCGACTTACCAT) that was subcloned into the same vector backbone as functional shRNAs.

### AAV brain injections

All surgical procedures were performed under aseptic conditions and deep anesthesia. Adult mice (7 to 18 weeks) were anesthetized using an intraperitoneal (i.p) injection of anesthesia mix (Medetomidine 0.5 mg/kg, Midazolam 5 mg/kg and Fentanyl 0.05 mg/kg). Mice were placed on a stereotaxic apparatus (Model 1900; Kopf, USA) and kept warm using a heating pad at 33.5°C. Fur of the head was removed, skin disinfected (Braunol; Braun, Germany) and cornea moisture was preserved during surgery by the application of eye ointment (Bepanthen; Bayer, Germany). Craniotomies of approximately 0.5 mm diameter were drilled on the skull with a hand drill (OS40; Osada Electric, Japan). A pulled-glass capillary with 20–40 µm tip diameter was lowered into the brain and specific recombinant adeno-associated virus (rAAV) carrying the functional construct or a fluorescent protein was injected using a manual air pressure system.

The following AAVs and titers were used:

- ssAAV-DJ/2-hSyn1-chI-dlox-hChR2(H134R)_mCherry(rev)-dlox-WPRE-hGHp(A) (Zurich Vector Core, 5.3 x 10E12 vg/ml)
- ssAAV-DJ/2-hSyn1-chI-dlox-mCherry(rev)-dlox-WPRE-hGHp(A) (Zurich Vector Core, 7.2 x 10E12 vg/ml)
- ssAAV-5/2-hSyn1-chI-dlox-EGFP_2A_FLAG_TeTxLC(rev)-dlox-WPRE-SV40p(A) (Zurich Vector Core, 7.7 x 10E12 vg/ml)
- ssAAV-1/2-hEF1α-dlox-hM4D(Gi)_mCherry(rev)-dlox-WPRE-hGHp(A) (Zurich Vector Core, 4.5 x 10E12 vg/ml)
- ssAAV-1/2-hSyn1-chI-dFRT-EGFP_2A_FLAG_TeTxLC(rev)-dFRT-WPRE-hGHp(A) (Zurich Vector Core, 5.0 x 10E12 vg/ml)
- ssAAV-1/2-hSyn1-dlox-EGFP(rev)-dlox-WPRE-hGHp(A) (Zurich Vector Core, 6.7 x 10E12 vg/ml)
- rAAV2/9-CAG::FLEX-rev-hrGFP:mir30(Scn9a) (a gift from Scott Sternson, 1.5-1.7 10E13 GC/ml)
- rAAV2/9-CAG::FLEX-rev-hrGFP:mir30(Scn9a-scrambled) (a gift from Scott Sternson, 1.5-1.7 10E13 GC/ml)
- ssAAV-1/2-hEF1α-dlox-hM3D(Gq)_mCherry(rev)-dlox-WPRE-hGHp(A) (Zurich Vector Core, 4.0 x 10E12 vg/ml)
- ssAAV-retro/2-hSyn1-chI-dlox-mCherry_2A_FLPo(rev)-dlox-WPRE-SV40p(A) (Zurich Vector Core, 6.3 x 10E12 vg/ml)
- ssAAV-retro/2-hSyn1-chI-dlox-EGFP_2A_FLPo(rev)-dlox-WPRE-SV40p(A) (Zurich Vector Core, 9.9 x 10E12 vg/ml)
- ssAAV-1/2-hSyn1-dFRT-hM4D(Gi)_mCherry(rev)-dFRT-WPRE-hGHp(A) Zurich Vector Core, 8.4 x 10E12 vg/ml)
- ssAAV-1/2-shortCAG-dlox-miR(Nav1.3-v1)(rev)-hrGFP(rev)-dlox-WPRE-SV40op(A) (Zurich Vector Core, 1.0 x 10E13 vg/ml)
- ssAAV-1/2-shortCAG-dlox-miR(Nav1.3-v2)(rev)-hrGFP(rev)-dlox-WPRE-SV40op(A) (Zurich Vector Core, 8.9 x 10E12 vg/ml)
- ssAAV-1/2-shortCAG-dlox-miR(Nav1.3-v3)(rev)-hrGFP(rev)-dlox-WPRE-SV40op(A) (Zurich Vector Core, 7.8 x 10E12 vg/ml)
- ssAAV-1/2- shortCAG-dlox-miR(Nav1.3-scrambled)(rev)-hrGFP(rev)-dlox-WPRE- SV40op(A) (Dirk Grimm laboratory, Heidelberg University, 1.9 x 10E12 vg/ml)
- ssAAV-8/2-CAG-EGFP_Cre-WPRE-SV40p(A) (Zurich Vector Core, 2.1 x 10E12 vg/ml)
- ssAAV-1/2-hSyn1-chI-iCre-WPRE-SV40p(A) (Zurich Vector Core, 5.2 x 10E12 vg/ml)
- AAV.DJ/2.hEF1α.dlox.GCaMP6f(rev).WPRE.bGHp(A) (Zurich Vector Core, 4.8x 10E12 vg/ml vg/ml)

Skin was sutured with sterile absorbable-needled sutures (Marlin 17241041; Catgut, Germany) and mice were injected subcutaneously with Carprofen at 5 mg/Kg (Rimadyl; Zoetis, USA). Finally, anesthesia was antagonized using a subcutaneous injection of Atipamezole 2.5 mg/kg, Flumazenil 0.5 mg/kg and Naloxon 1.2 mg/kg and mice were transferred to their home cages. For postoperative care, a second dose of Caprofen was injected after 24 hours and mice cages were kept on a veterinary heating pad at 37°C for 12 hours and monitored closely. A minimum of 3 weeks of viral expression was allowed before any experiments were conducted.

### Telemetry transmitter implantation

All animals (with the exclusion of those used for electrophysiological recordings) were implanted with a telemetry transmitter (TA11TA-F10; Data Sciences International, USA) to monitor body temperature during acclimation procedure and behavioral testing. Animals were injected with anesthesia mix i.p. as described above, fur of the abdomen was removed, skin disinfected with Braunol (Cat# 3864065, Braun, Germany) and cornea protected with Bepanthen ointment (Bayer, Germany). A sterile telemetric transmitter was implanted in the abdominal cavity. Thereafter, muscle and skin layers were separately sutured with absorbable surgical threads (Marlin Cat# 17241041, Catgut, Germany). After the surgery, the anesthesia was antagonized and animals were monitored as described above; recovery for at least one week was allowed before any further procedures were undertaken.

### Tail, iBAT, and core body temperature measurement

In ChR2-encoding AAV-injected mice (and respective control animals), tail temperatures and brown adipose tissue temperatures were measured using an infrared thermal camera (VarioCAMhr, InfraTec, Germany). Snapshot images were taken every 5 min using IRBIS 3 software (InfraTec, Germany). The average temperature was calculated in the middle of the tail (segment length of 1 cm) and at the center of the interscapular region, which was shaved 3 – 5 days before measurement. Core body temperature of was sampled every 5 min via receiver plates (RSC-1, DSI, USA) placed underneath the cages. Telemetry data were registered using Ponemah (DSI, USA). All measurements were conducted during the light phase.

### Optogenetic stimulation of LepR cells

Stereotactic surgeries were performed in adult LepR^Cre^. Animals were injected bilaterally with 250 nL of AAV encoding the Cre-dependent ChR2 or mCherry (control group) (ssAAV-DJ/2-hSyn1- chI-dlox-hChR2(H134R)_mCherry(rev)-dlox-WPRE-hGHp(A) or ssAAV-DJ/2-hSyn1-chI-dlox-mCherry(rev)-dlox-WPRE-hGHp(A)) at coordinates targeting VMPO: Bregma: ML: ±0.400 mm, AP: 0.800 mm, DV: -4.850 mm (VMPO). A 200 μm diameter fiber optic probe (Cat# FT200UMT, ThorLabs) was lowered to target preoptic LepR cell population (coordinates: Bregma: ML: 0.400 mm, AP: 0.800 mm, DV: -4.700 mm (VMPO)). The probe was anchored to the skull with dental acrylic. After the surgery, the anesthesia was antagonized and mice were transferred to their home cages. Postoperative care and telemetry implantation was performed as described above. At least four weeks were allowed for recovery and full expression of ChR2 before the start of optogenetic stimulation.

To activate ChR2- expressing LepR neurons, fiber optic probe was attached through an FC/PC adaptor to a 473-nm blue LED (Optogenetics-LED-Blue, Prizmatix). All experiments were conducted unilaterally, and the fiber optic cable was connected at least 2 hr before the experiments to allow for habituation. For the optogenetic probing, mice received light pulses of 4–6 mW power, 10 ms width and delivered at 20 Hz stimulation frequency using a Prizmatix Pulser software and pulse train generator (Prizmatix). In each optogenetic probing experiment, the light stimulation period was of 1 min followed by an inter-stimulation interval of 3 min.

### TeTxLC and Gi-DREADD silencing of LepR cells

For these experiments, stereotactic surgeries were performed in adult LepR^Cre^ mice as described in previous sections. 250 nL of rAAV encoding the Cre-dependent tetanus toxin light chain (TeTxLC) (ssAAV-5/2-hSyn1-chI-dlox-EGFP_2A_FLAG_TeTxLC(rev)-dlox-WPRE- SV40p(A)) or the inhibitory Gi-DREADDs (ssAAV-1/2-hEF1α-dlox-hM4D(Gi)_mCherry(rev)- dlox-WPRE-hGHp(A)) was injected bilaterally in VMPO. AAVs encoding a Cre-dependent mCherry/EGFP were used as controls. Following brain injection, the anesthesia was antagonized, and mice were transferred to their home cages. Postoperative care was performed as described before. After telemetry implantation, heat (or RT) acclimation and heat endurance assay were performed as detailed in previous sections.

Acute chemogenetic silencing of LepR cells was performed by injecting CNO (or saline) 0.3 mg/kg i.p. (Enzo, diluted in saline) 10 min before transferring the animals to the heat endurance assay. Body temperature was constantly monitored as mentioned above. In order to validate CNO effects on the firing frequency of acclimated LepR cells, a group of chemogenetically silenced animals were used for *in vitro* electrophysiological recordings. Slice preparation and electrophysiological recording procedure are described below.

### Gq-DREADD- and ChR2- stimulation and long-term activation (optogenetic- and chemogenetic conditioning) of LepR cells

For these experiments, stereotactic surgeries were performed in adult LepR-cre mice as described in previous sections. 250 nL of AAV encoding for Cre-dependent excitatory Gq-DREADDs (ssAAV-1/2-hEF1α-dlox-hM3D(Gq)_mCherry(rev)-dlox-WPRE-hGHp(A)) or Cre-dependent ChR2 (ssAAV-DJ/2-hSyn1-chI-dlox-hChR2(H134R)mCherry(rev)-dlox-WPRE-hGHp) was injected bilaterally into the VMPO.

To mimic acclimation by optogenetic activation of VMPO^LepR^ cells, ChR2-expressing mice received light pulses via the fiber optic probe of 10 ms duration / pulse (4–6 mW), triggered at 1 Hz frequency. The stimulation was protracted continuously for the duration of heat challenge (max: 9hr) or was started 1 day or 3 days prior to heat endurance. The degree of hypothermia produced by this continuous optogenetic stimulation was tested in a different cohort of mice at RT.

For chemogenetic activation of VMPO^LepR^ cells to mimic acclimation, animals were injected daily with CNO (0.3 mg/kg i.p., Enzo, diluted in saline) for 1, 5 or 10 consecutive days. CNO effect on body temperature was monitored constantly. At the end of the injection period, and after 24 hrs from the last injection, animals were tested in the heat endurance assay while the temperature was recorded telemetrically as described above. A group of long-term chemogenetically stimulated animals were used for electrophysiological recordings.

Repeated administration of CNO to control mice (in the absence of Gq-DREADD) over a period of 10 days also had a small but discernible effect, in particular on the kinetics of Tcore at the initial phase of the heat endurance assay, possibly reflecting the activity of CNO metabolites such as clozapine, known to modulate several neuronal receptor systems^83,84^. Nevertheless, this effect was considerably smaller than that observed in mice carrying the chemogenetic actuator Gq-DREADD. Of note, all mice carrying Gq-DREADD and chemogenetically conditioned for 10 days reached the cut-off time in the heat endurance assay, suggesting that their gained heat tolerance ––and different to that of any of the other experimental groups––was underestimated in this assay (Extended Data Fig. 9c and d).

### Silencing of LPBN neurons

Vglut2-cre mice were injected 250 nL of retro-AAV encoding the Cre-dependent FlpO recombinase (ssAAV-retro/2-hSyn1-chI-dlox-mCherry_2A_FLPo(rev)-dlox-WPRE-SV40p(A) or ssAAV-retro/2-hSyn1-chI-dlox-EGFP_2A_FLPo(rev)-dlox-WPRE-SV40p(A)) bilaterally in VMPO. After 3 weeks, the same animals received 250 nL bilateral injection of AAV encoding the FlpO-dependent TeTxLC or the inhibitory Gi-DREADD (ssAAV-1/2-hSyn1-chI-dFRT- EGFP_2A_FLAG_TeTxLC(rev)-dFRT-WPRE-hGHp(A) or ssAAV-1/2-hSyn1-chI-dFRT- EGFP_2A_FLAG_ hM4D(Gi)(rev)-dFRT-WPRE-hGHp(A)) at Bregma: ML: ±1.25 mm, AP: - 4.900 mm, DV: -2.7 mm (lateral parabrachial nucleus / lPBN). After telemetry implantation, heat acclimation was performed as detailed in previous sections.

Acute chemogenetic silencing of LPBN presynaptic partner cells was performed by injecting CNO (or saline) 0.3 mg/kg i.p. (Enzo, diluted in saline) at the end of the acclimation protocol, and 10 min before transferring the animals to the heat endurance assay. Cre-negative animals were subjected to the same injection procedure and served as controls. Body temperature was constantly monitored for all animals during the acclimation period and/or the heat endurance assay.

### Nav1.7 / Nav1.3 knock-down in VMPO^LepR^ cells

Animals were anaesthetized as described above and shRNA virus against Scn9a / Nav1.7 (rAAV2/9-CAG::FLEX-rev-hrGFP:mir30(Scn9a) or scrambled control (rAAV2/9-CAG::FLEX-rev-hrGFP:mir30 (Scn9a-scrambled))^62^ or against Scn3a / Nav1.3 (ssAAV-1/2-shortCAG-dlox- miR(Nav1.3-v1/v2/v3)(rev)-hrGFP(rev)-dlox or scrambled control ssAAV-1/2- shortCAG-dlox- miR(Nav1.3-scrambled)(rev)-hrGFP(rev)-dlox) were injected into POA to target LepR^+^ neurons (250 nL, bilaterally). The three Nav1.3-targeting shRNA AAVs (v1, v2 and v3) were mixed at 1:1:1 proportions prior to injections. After recovery and acclimation, animals were used for *in vitro* electrophysiological recordings.

### Conditional Nav1.3 knock-out

To create conditional Nav1.3 knock-out, Nav1.3 floxed mice were brought to homozygosity (Nav1.3^fl/fl^) and injected with Cre-encoding AAV (AAV8 CAG EGFP-Cre) into VMPO. Wildtype mice (Na_V_1.3^+/+^) were injected with Cre-encoding AAV to serve as controls. At least 3 weeks of virus expression / protein turnover was allowed before subjecting the cKO mice and controls to heat acclimation. A mix of Cre-AAV injected WT littermates and WT C57BL/6 mice was used as control group.

### Histology of AAV-injected mouse brains

Mice were anesthetized, transcardically perfused with PFA and decapitated. The entire heads were left in 4% PFA for at least one day at 4°C. Subsequentially, the brains were removed from the skull and transferred to a PBS solution containing sucrose. Coronal sections of 30 µm were cut at the microtome and stored at -20°C in cryo-protectant solution. Subsequently, brain sections were stained for GFP / mCherry as described previously.

### LepR cell RNA-seq: LepR cell dissociation

Adult mice LepR-Cre;HTB, acclimated and non-acclimated, (P77 ± 5) were anesthetized with isoflurane and decapitated. The brain was immediately removed and submerged in ice-cold artificial cerebrospinal fluid (ACSF). 3 brains were sectioned at the same time on a vibratome (Leica VT1200S) in a slicing chamber containing ice cold aCSF: NaH2PO4 (1.2 mM), KCl (1.2 mM), HEPES (20 mM), Glucose (25 mM), NaHCO3 (30 mM), NMDG (93 mM), Na-Ascorbate (5 mM), Na-Pyruvate (3 mM), N-acetylcysteine (12 mM), CaCl2 (0.5 mM), MgSO4x7H2O (10 mM), constantly bubbled with carbogen. Brain slices of 250 μm thickness, containing the rostral POA, parts of the Orgnaum Vasculosum Laminae Terminalis (OVLT) and Medial Preoptic Nucleus (MnPO), were transferred to a Petri dish containing ACSF. We implemented the neuron isolation protocol described in^20^. The regions of interest were micro dissected under a dissecting microscope and transferred to a small Petri dish containing 3 ml of Papain mix consisted of Hibernate mix (Hibernate A medium (Invitrogen A1247501), 1xGlutamax (Gibco 35050-038), 0.8 mM Kynurenic acid (Sigma K3375-5G), 0.05 mM AP-V (HelloBio HB0225), 0.01 mM Rock inhibitor Y-27632 (HelloBio, HB2297), 1 mM B27 (Invitrogen 17504001), 5% Threhalose (Sigma, T9531-10G)) and 8 U/ml papain (Sigma P4762), 100 U/ml DNAseI (Worthington über Cell-Systems, LK003172), 0.005 U/ml Chondroitinase ABC (Sigma, C3667-5UN), 0.07% Hyaluronidase(Sigma, H2126), 0.001 mM NaOH. The tissue was cut in smaller pieces and transferred together with the Papain mix into a 2 ml tube at 37 °C to incubate while shaking (700 rpm) for 2 h. After incubation the papain solution was pipetted out of the tube and exchanged by hibernate mix containing 0.1mg/ml ovalbumin and centrifuged for 1 min at 300 g. Supernatant was removed and hibernate mix was added to the tissue pieces which were further dissociated into single cells by gentle trituration through Pasteur pipettes with fire polished tip openings of 600 μm, 300 μm, and 150 μm diameter. Cell suspension was centrifuged at RT at 300 g, for 10 minutes, supernatant was removed and exchanged with 500μl of Hibernate A medium. Resuspended cell material was passed through a 20 µm filter. Cell suspension was stained with propidium iodide (BD Pharmingen, 5166211E) to exclude the dead cells before the FACS analysis. FACS sorting was performed on a BD FACS Aria II using the purity sorting mode. FACS populations were chosen to select cells with low PI and high GFP fluorescence.

Cells were FACS sorted into bulks of GFP+ and GFP- directly into the RLT buffer (QIAGEN RNeasy Micro Kit, 74004), according to the arbitrary levels of GFP fluorescence, immediately frozen on the dry ice and stored at -80°C. Samples were processed in maximum one month from the isolation by using column purification method according to manufacturer’s instructions (QIAGEN RNeasy Micro Kit, 74004) and samples were stored at -80 °C until further processing.

### LepR cell RNA-seq: cDNA library preparation

RNA integrity and concentration of each sample were assessed by Agilent Bioanalyzer Nano 6000 chip (Agilent technologies) and QUBIT (Invitrogen QUBIT2) measurement. We used Smart seq2 protocol^85^ for the cDNA library preparation (all processing performed at Gene Core EMBL, Heidelberg). 200 pg of each RNA bulk sample was processed for the reverse transcription (Superscript IV) followed by 18 cycles of PCR amplification, library tagmentation (Tn5 transposase produced in house, PEP Core, EMBL Heidelberg), sample barcoding and final 12 PCR cycle enrichment. All samples were sequenced on Illumina NextSeq 500 High sequencer, single end with 75 base pairs long reads (Gene Core EMBL, Heidelberg).

### POA slice preparation for electrophysiology

For *in vitro* electrophysiology, 8 to 15 week old mice were deeply anaesthetized using a Ketamine/Xylaxine mixture (ketamine: 220 mg/kg, Ketavet; Zoetis, USA and Xylazine 16 mg/kg, Rompun; Bayer, Germany), decapitated, and brains transferred to ice-cold (4°C) oxygenated (95% O2, 5% CO2) slicing artificial cerebrospinal fluid (aCSF; in mM): NaCl, 85; KCl, 2.5; glucose, 10; sucrose, 75; NaH2PO4, 1.25; NaHCO3, 25; MgCl, 3; CaCl2, 0.1; myo-inositol, 3; sodium pyruvate, 2; ascorbic acid, 0.4. Coronal (250 μm thick) POA slices were prepared with a vibratome (Leica VT1200S, Germany) and then incubated at 32°C in a bath containing oxygenated holding aCSF (in mM): NaCl, 109; KCl, 4; glucose, 35; NaH2PO4, 1.25; NaHCO3, 25; MgCl2, 1.3; CaCl2, 1.5. After a recovery period of 30 minutes, individual slices were transferred to the recording chamber where they were continuously superfused with oxygenated recording aCSF (for recipes see below) at ∼2 ml/min.

In some experiments, brain slices were prepared using carbogen-bubbled NMDG-HEPES solution (at 4°C) containing (in mM): NMDG, 93; KCl, 2.5; NaH2PO4, 1.2; L(+) ascorbic acid, 5; Thiourea, 2; sodium pyruvate, 3; MgSO4.7H2O, 10; CaCl2.2H2O, 0.5; HEPES, 20; NaHCO3, 30; Glucose, 25 and N-acetyl-L-cysteine, 10 (pH = 7.37-7.38, 295-305 mOsm/kg). After slicing, POA coronal slices were incubated for 15 min in the same NMDG-HEPES solution at 32°C and subsequently transferred to a chamber containing holding aCSF composed of (in mM): NaCl, 118; KCl, 2.5; NaHCO3, 24; NaH2PO4, 1.2; sodium pyruvate, 2.4; L(+) ascorbic acid, 4; N-acetyl-L-cysteine, 2; HEPES, 5; MgSO4, 1; CaCl2, 2 and glucose, 7 (pH = 7.3-7.5, 295-305 mOsm/kg).

Cells in acute POA slices were visualized using a SliceScope upright microscope (Scientifica, UK) equipped with a 40X water immersion objective (U-TV1X-2, Olympus, Japan). Images were acquired by a digital CCD camera (ORCA-R2 C10600-10B, Hamamatsu Photonics K.K., Japan) using MicroManager 1.4 software (Vale’s lab, UCSF, USA). Electrophysiological recordings were acquired using a MultiClamp 700B amplifier (Molecular Devices, USA), together with an Axon Digidata 1550B digitizer (Molecular Devices, USA) and Clampex 11.0.3 software (Molecular Devices, USA). All signals were sampled at 20 kHz and low pass filtered at 10 kHz. Borosilicate glass micropipettes used (O.D. 1.5 mm, I.D. 0.86 mm, Sutter Instrument, BF150-86-7.5) were pulled on a micropipette puller (P-97, Sutter Instrument, USA). Intracellular solution was passed through 0.22 µm filter before filling the electrode pipette. The open pipette resistance was between 4-8 MΩ.

### Electrophysiological measurement of warmth-sensitivity of VMPO neurons

In acute slice experiments where neuronal action potentials were recorded at varying temperatures, a bridge in a form of glass capillary filled with agar dissolved in 3 M KCl was placed between the bath chamber and the ground electrode in order to isolate the reference electrode from the temperature changes applied to the chamber^86^. Equipment for bath temperature control consisted of temperature-controlled microscope stage (TC07, Luigs & Neumann), an in-line heater (CL-100, Warner) and a liquid cooling system (LCS-1, Warner).

Neuronal action potentials were recorded with aCSF containing (in mM): NaCl, 125; KCl, 6.25; glucose, 15; NaH2PO4, 1.25; NaHCO3, 25; MgCl, 1.3; CaCl2, 2.4 (called “high-K^+^ aCSF”) as previously described^86^ and with an internal solution containing (in mM): K-gluconate, 138; KCl, 2; NaCl, 5; HEPES, 10; EGTA, 10 (or equimolar amount of BAPTA); CaCl2, 1; Mg-ATP, 1.

AP frequencies were analysed in traces where the bath temperature was either 33°C, 36°C or 39°C; deviation of maximum ± 0.5°C was tolerated. Neurons were classified as warm-sensitive when their temperature coefficient reached 0.75 Hz /°C and as cold-sensitive when their temperature coefficient was lower than -0.6 Hz /°C, thresholds traditionally used to define central temperature sensitive neurons^87^. Temperature-insensitive neurons had their temperature coefficient between ≥- 0.6 Hz /°C and <0.75 Hz /°C, and neurons were classified as silent when not a single spontaneous AP could be detected. Cells unable to produce AP even when stimulated with current injection were excluded from analysis. Probing VMPO neuronal populations for temperature sensitivity was done in the presence of synaptic blockers (gabazine 5 µM, CNQX 10 µM, AP-V 50 µM) added to the bath solution. In experiments where the effect of cholinergic transmission was tested, 10 μM tubocurarine and 10 μM scopolamine were included in perfusion fluid.

In some experiments, action potentials were measured without varying bath temperature (at 33°C) and with a “low-K^+^ aCSF”, containing NaCl, 125; KCl, 2.5; NaHCO3, 24; NaH2PO4, 1.2; HEPES, 5; MgSO4, 1; CaCl2, 2 and glucose, 8. The solution used and temperature of recordings is indicated in the figure legends showing spontaneous AP firing data.

### Recordings of ionic currents

Resting membrane potential was measured in current clamp mode using extracellular solution containing (mM): NaCl, 150 (or equimolar amount of NMDG); KCl, 3,5; HEPES, 10; glucose, 20; CaCl2, 1.2; MgCl2, 2 (as per Lu et al.^88^). TTX (0.5 μM) was added to the aCSF and pipette solution contained (mM): K-gluconate, 120; HEPES, 40; MgCl2, 5; Na2ATP, 2; NaGTP, 0.3.

To record voltage ramp responses in voltage clamp mode to approximate passive membrane permeability to potassium, we used a low-sodium and 0 mM nominal calcium solution that contained (mM): NMDG, 125; NaHCO3, 24; KCl, 2.5; NAH2PO4, 1.2; HEPES, 5; glucose, 8; MgSO4, 1. Pipette solution contained (mM): Cesium methanesulphonate, 120; HEPES, 40; MgCl2, 5; NaATP, 2; NaGTP, 0.3; QX-314, 5; TEA-chloride, 5; 4-Aminopyridine, 1 – in order to block voltage-gated potassium and sodium channels. “Leak” potassium channels are largely unaffected by intracellular cesium^89^.

Voltage ramps as well as voltage-gated calcium currents were recorded using the same Cesium methanesulphonate-based pipette solution and external solution composed of (mM): NaCl, 125; NaHCO3, 24; KCl, 2.5; NAH2PO4, 1.2; HEPES, 5; glucose, 8; MgSO4, 1; CaCl2, 2. To record voltage-gated calcium currents, aCSF additionally contained 0.5 μM TTX, 1 mM TEA-chloride and 100 μM 4-AP.

To measure voltage-gated sodium currents in whole-cell and nucleated patch configurations, we used solutions as described by Milescu et al^90^. Here, external solution contained (in mM): NaCl, 124; KCl, 3; glucose, 30; NaH2PO4, 0.5; NaHCO3, 25; MgSO4, 1; CaCl2, 1.5 with addition of TEA-chloride (5mM) and CdCl2 (50 μM). Pipette solution contained (in mM): Cs-gluconate, 100; NaCl, 4; TEA-chloride, 10; 4-AP, 5; EGTA, 10; CaCl2, 1; HEPES, 10; MgATP, 4; NaGTP, 0.3; Na-phosphocreatine, 4.

Spontaneous synaptic currents were recorded with “low-K^+^ aCSF” and Cesium methanesulphonate-based pipette solution. sEPSCs were recorded while holding the neurons at - 65 mV in gap-free mode; sIPSC were recorded at the potential of 0 mV (reversal potential for AMPA currents).

*In vitro* validation of DREADD receptor function was performed in current-clamp mode using “low-K^+^ aCSF”, K-gluconate-based intracellular solution as described above and with the addition of 5 µM CNO.

For AP frequency quantification, first 3 minutes of recordings were omitted; in voltage clamp recordings, at least 1 minute was allowed after break-in before any recording was performed. All ionic current recordings were conducted at 36°C (± 0.5°C) in order to mimick more closely the physiological neuronal conditions. Basic cell membrane properties such as capacitance and input resistance were calculated based on a membrane test protocol (a brief step of -10 mV from a holding potential of -65 mV). Series resistance (Rs) was typically 10–25 MΩ across experiments. In voltage clamp recordings, whole-cell capacitance compensation was applied and Rs was compensated 50-60%; the compensation was readjusted before each protocol. Voltage protocols applied are shown in data figure insets. In current-clamp experiments, pipette capacitance neutralization and bridge-balance were used. In experiments where voltage-gated sodium currents were measured, a liquid junction potential (LJP) of 8 mV was corrected on-line; in other experiments, LJP was corrected off-line. In voltage clamp experiments, cells whose membrane resistance changed by >50% or series resistance changed by >20% between start and end of recording were excluded from analysis. An in-house software was developed for the automated analysis of the action potential (AP) waveforms.

### Quantitative PCR

The animals were sedated with isoflurane and sacrificed via cervical dislocation 3 weeks after injection of shRNA AAVs to the preoptic area. The whole brain was prepared and stored in cold DPBS (Gibco). The brain was cut with the help of a mouse brain matrix and the whole POA was extracted and transferred to an Eppendorf tube, which was subsequently filled with TRIzol reagent (Ambion). RNA was extracted using the TRIZOL (Cat# 15596026, Ambion) and ROTI Phenol/Chloroform/Isoamyl alcohol (Cat# A156, Carl Roth) protocol. The preoptic area tissue was transferref from TRIzol solution to a glass mortar and manually disrupted with a pestle. Subsequently, disrupted tissue was suspended in ROTI Phenol/Chloroform/Isoamyl. The samples were centrifuged at 14000 rpm in tabletop centrifuge at 4°C for 10 minutes. The resulting aqueous phase was transferred to a spin column for purification (Cat# R1013, Zymo Research) and the eluted RNA was stored at -80°C until further analysis.

600 ng of total RNA were used for first-strand cDNA generation with SuperScript III Reverse Transcriptase (Cat# 10368252, ThermoFisher Scientific) using oligodT primers according to the manufacturer’s instructions. The resulting cDNA was diluted to a concentration of 600 ng/µL. cDNA was analysed by quantitative PCR using the following primers specific for the *Scn3a* transcript. *Ube2l3* and *Tubb3* served as housekeeping genes. Primer sequences are listed in the following table.

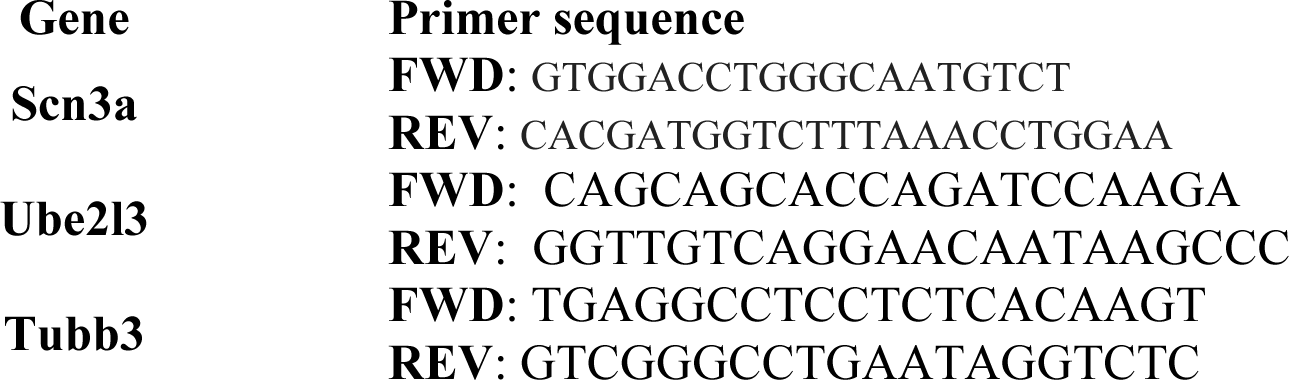

qPCR amplification reactions (15 µL) contained 7.5 µL FastStart Essential DNA Green Master Mix (Cat# 06402712001, Roche), 5 µL RNase free water (Qiagen), 1 µL cDNA, and 1.5 µL of forward and reverse primer. Reactions were run on a Roche LightCycler 96 System (Roche Diagnostics). Controls without RT were included to control for traces of genomic DNA. No template controls were included to check for contamination and non-specific amplification. The resulting CT values were exported as text files and imported into Microsoft Excel for further analysis. The acquired data were analysed by an approach described in Vandesompele et al. 2002 and Hellemans et al. 2007^91,92^. Data are expressed as relative gene expression ratios. All samples were measured in triplicates.

### Microendoscopy of VMPO^LepR^ neurons in awake behaving mice

Experiments were performed in 8-week-old male LepR-Cre+/- mice. Each mouse underwent two sequential stereotaxic surgeries, one for injections of an adeno-associated viral (AAV) vector expressing GCaMP6f, and a second one performed 7 days later to implant a GRIN lens attached to a baseplate.

For AAV injection, mice were deeply anesthetized with 2% isoflurane at a flow rate of 0.5 L/min and placed in a stereotactic frame (Kopf Instruments, Tujunga, CA). Body temperature was maintained at 37°C with a heating pad (Hot-1, Alascience, Scientific Instruments). Ophthalmic ointment (Bepanthen, Bayer Vital GMBH, Germany) was applied to the eyes to prevent drying. Upon deep anesthesia, mice underwent bilateral craniotomies at two anteroposterior locations, using a high-speed rotary micro drill (Stereotaxic Drill, Kopf Instruments). The following stereotaxic coordinates were used: 0.2 mm AP ± 0.4 mm ML, and 0.5 mm AP ± 0.4 mm ML. Then, a glass pipette filled with GCaMP6f delivered the virus into the VMPO (5 mm DV). In each injection site, 200 nL of a 1:3 virus dilution in saline solution was injected, using a NanoJet microinjector (World Precision Instruments) at a rate of 20nL/min. After each injection, the pipette was left at the injection site for 10 min to avoid backflow, and then slowly withdrawn. The skin was then sutured with nylon suture thread (Dafilon, Braun, Germany).

Implantation GRIN lens and baseplate. Seven days after the first surgery, mice were again anesthetized as previously described, and a 0.6x7 mm gradient refractive index (GRIN) lens with an integrated baseplate for the nVista miniaturized head-mounted microscope (‘Miniscope’, Inscopix, USA) was slowly inserted to the brain (60 µm/min) on one of the hemispheres previously injected. The following stereotaxic coordinates were used: 0.35 mm AP, 0.4 mm ML, 5 mm DV. Then, the baseplate was fixed to the skull using a self-curing adhesive resin (Super-Bond, Sun Medical, Japan) and a light cured composite resin (Gradia®-Direct Flo, GC Corporation, Japan). The surface of the lens was then covered with a plastic basecap (Inscopix, USA), and the skin was sutured with nylon suture thread (Dafilon, Braun, Germany).

6-7weeks after implantation, the pre-acclimation recordings were performed. To dock the Miniscope to the baseplates, mice were briefly anesthetized with isoflurane (2%). Upon docking, the Minsicope was locked to the baseplate with a small screw. Mice recovered from anesthesia for at least 30 minutes, at RT, in a plexiglass box (25x25 cm) with bedding. Recording of calcium signals were first performed at RT, at 5 different focal points. At each focal point calcium transients were recorded for 2 minutes, at 20 Hz, with a maximum resolution of 1280 × 800 pixels, and an LED power of 1-1.1 mW/mm2. After this, mice were moved to the heating chamber that was maintained at near 36°C (±2°C). Recordings at high temperature started 5 minutes after mice were transferred into the heating chamber, using the same parameters as low temperature recordings. Once the pre-acclimation recording session finished, mice were temperature-acclimated for 30 days, as described in the previous sections. Post-acclimation recordings were performed following the same procedure as for pre-acclimation recordings described above. In control experiment, mice were recorded at identical timepoint and in identical conditions, but maintained at RT between recording sessions (30 days).

After recordings, mice were anaesthetized with an intraperitoneal injection of narcoren, trans- cardially perfused with 4% PFA in PBS, and post-fixed in 4% PFA for 48 hours at 4°C. After this, the implant was removed, the brain dehydrated in 30% sucrose overnight at RT, and cut in 30um sections using a sliding microtome (Hyrax S50 and KS34, Zeiss). Four series were generated for each mouse’s VMPO, and one of these series was mounted and imaged with EPI fluorescence microscope (10x, DM6000, Leica) to assess virus expression and implant location within the MPOA.

Raw recordings were first pre-processed using the Inscopix Data Processing Software (IDPS version 1.6.0.3225). For each experimental session, video-recordings obtained at 22 and 36 C were merged and processed together. The Timeseries module of the IDPS toolbox was used to merge the two videos. Raw imaging data was cropped to accommodate only the desired region of interest and to remove the lens boundary artifact. Recordings were then filtered using the Spatial Filter module that removes the low and high spatial frequencies, (measured as the number of oscillations per pixel) thus effectively reducing out-of-focus background fluorescence. We used a trial-and-error method to identify the filter cut-off values and found that the best low cut-off value was 0.005 and the high cut-off value was 0.090. The file was then motion corrected using the Motion Correction module of IDPS. The mean image of the video file was considered as the global reference. The maximum translational value for a pixel was also estimated using the trial-and-error method and it varied between the ranges of 20 and 40 for recording from different mice. These files were then exported as .tiff image stacks and were used to extract calcium transients using EZcalcium^93^ extraction and analysis toolbox based on the CaImAn pipeline^94^. For this, we first performed manual ROI detection using the ROI Detection module to generate the fluorescence (ΔF/F) and the deconvolved neural spiking values. The deconvolution was performed using the Markov chain Monte Carlo^95^ method which is a fully-Bayesian deconvolution method and we used the Rise and Decay autoregression method to estimate the calcium indicator dynamics. We considered 0.9 as the merge threshold above which two neurons sharing a correlation coefficient would be merged into a single ROI. Calcium signals arising from each ROI were visually inspected and curated using the ROI Refinement module in EZcalcium. The data was then saved as a MATLAB data (.mat) file. Additionally, ROI masks were created for each file. These masks were then used for manual matching of neurons across different recording sessions. Raw ΔF/F values obtained from the EZcalcium module were then used to compute the baseline (22°C) z-score, (BZ) for each neuron, given by:

BZi = (xi-xib) / sib Where,

xi = fluorescence value during the baseline period xib = mean of values from the baseline period

sib = standard deviation of values from the baseline period

Post-stimulus (36°C) z-scores PZ were computed as a function of the BZ, given by, PZj = (xj-xib) / sib

Where,

xj =fluorescence value of a neuron after stimulus

xib = mean of values of the xj neuron from baseline period

sib = standard deviation of the xj neuron values from baseline period.

The z-score computation was performed using a custom python code and was then exported as a

.xlsx file for further analysis.

### Human brain tissue *in situ* hybridization

RNAscope fluorescent in situ hybridization (FISH) was performed on formalin-fixed, paraffin-embedded (FFPE) human brain sections covering the VMPO (tissue blocks obtained from the Edinburgh Brain Bank in collaboration with Professor Colin Smith). The tissue was sectioned at 5 μm and mounted onto Fisher SuperFrost Plus glass slides (Fisher Scientific). Multiplex FISH was performed using the Leica RX Fully Automated Research Stainer (Leica) and the RNAscope LS multiplex fluorescent reagent kit (Advanced Cell Diagnostics, Bio-Techne) with Opal fluorophore reagent pack detection (Akoya BioSciences, Inc.). All slides were counterstained with DAPI and coverslipped with ProLong Diamond antifade mountant (ThermoFisher Scientific). The sections were hybridized with human-specific probes to detect mRNA transcripts for PACAP (ADCYAP1,#582508), LEPR-tv1 (long isoform, #410378-C2), LEPR-alltv (all isoforms, #410388), vGLUT2 (SLC17A6, #415678), PTGER3 (#488438) and OPN5 (#1058668-C2) (Advanced Cell Diagnostics, Bio-Techne). The slides were scanned with the Olympus VS200 slidescanner using (Olympus) a 20× air objective (0.8 NA) and a DAPI/CY3/CY5 filter set. Images were prepared with the Olympus OlyVIA software, and signal intensity levels were adjusted to match across staining/slides.

Luxol Fast Blue (LFB) and Hematoxylin-Eosin (HE) staining of myelinated fibers (blue) and cell bodies (purple) was performed on human FFPE brain sections to facilitate correct anatomical annotation. Standard staining protocol included: Deparaffination, LFB (Solvent Blue 38/ethanol/acetic acid, SigmaAldrich) incubation overnight followed by lithium carbonate/hematoxylin/acetic alcohol/lithium carbonate/eosin (Sigma Aldrich/Merck) incubation steps, dehydration in xylene and mounting with Pertex.

### Data and Statistical analysis

Data was analyzed using ImageJ (version 1.53c, NIH, USA), Olympus OlyVIA software, R and RStudio (version 1.2.5033), Python (version 3.7.6), Microsoft Excel, Igor Pro (version 6.37), MATLAB (version R2021a). Statistical tests were performed using R or GraphPad Prism (V5.00 and V6.00; GraphPad software, USA). N numbers in each figure legend are displayed in the format “N/n”, with N being the number of mice and n the total number of cells recorded. Results are presented either as mean ± standard error of the mean (s.e.m.) or as median and interquartile range. Distribution of data was assayed using the KS normality test, the D’agostino and Pearson omnibus normality test and the Shapiro-Wilk normality test. The difference between two groups was tested using a Two-Sample T-test or the non-parametric Mann-Whitney U test. For multiple group testing with ANOVA or Kruskal-Wallis Test, either Tukey’s HSD, Dunn’s or Sidak’s multiple comparisons tests were used as a post-hoc tests.

In endoscopic imaging experiments, we used non-parametric Kolmogorov-Smirnov test and compared the z-scores of individual cells at 22°C and 36°C to estimate if units increased, decreased, or did not change their activity upon acute increase of external temperature; to statically assess the impact of acclimation on temperature sensitivity, we compared averaged z-scores using two-sided Kolmogorov-Smirnov test or Wilcoxon signed-rank test. P-values < 0.05 were considered statistically significant. Star, ‘*’, signifies *P* ≤ 0.05, ‘**’ *P* ≤ 0.01, and ‘***’ *P* ≤ 0.001. Details on the statistical methods applied are included in the figure legends.

## Data availability

Individual data points are represented throughout all figures. Upon peer-reviewed publication, the associated data will be provided as Source Data Files, with all data that is presented in each main and Extended Data figure included in subfolders named correspondingly in the Source Data folder that will then be found on the HeiData server of Heidelberg University (https://heidata.uni-heidelberg.de/) under the following address: https://doi.org/10.11588/data/MRCFI2

## Code availability

All original code for electrophysiological data analysis will be deposited on Github and will be publicly available upon publication; for reviewers this code is available for download from this site: https://www.dropbox.com/sh/7bqb9gm2qzfssij/AABx58mFhPWsaZ8AiWcY0ytsa?dl=0 The MATLAB and Python codes used for endoscopic imaging data analysis can be accessed at the following Github link: https://github.com/AcunaLabUHD/AcunaLab_Miniscope_Siemens Any additional information required to reanalyze the data reported in this paper is available from the lead contact upon request.

---

**Extended Data Fig. 1.**
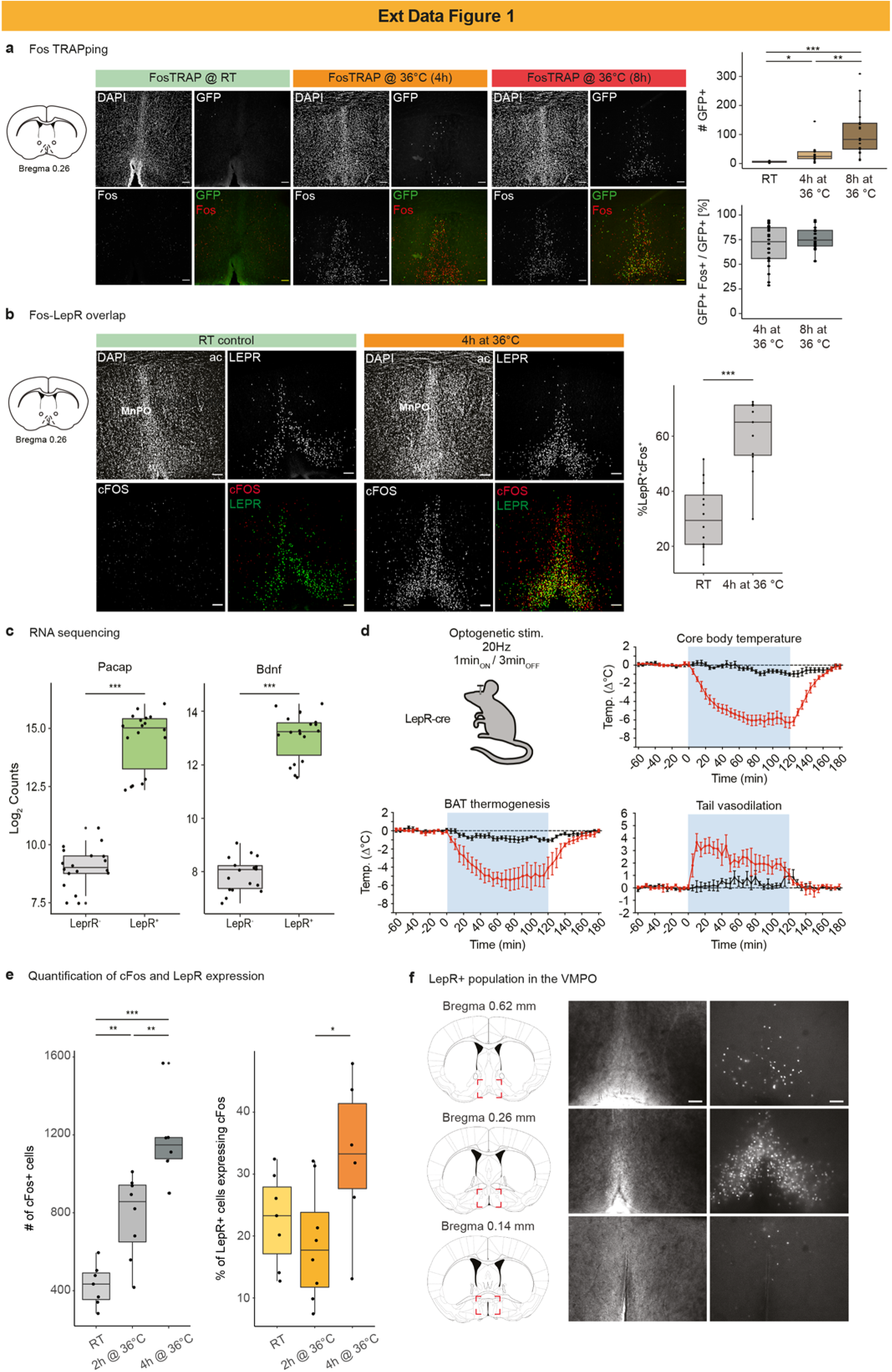
VMPO neurons responding with a delay to a heat stimulus overlap with LepR-positive neurons and are receptive to becoming activated by heat acclimation. a, Left: Representative images using fluorescent confocal imaging of brain sections revealing the VMPO of FosTRAP2;HTB mice that received z-4-hydroxytamoxifen (4-OHT; 50mg/kg i.p.) at room temperature (TRAP@RT) and after 2 hours of a 4-hour (TRAP@36°C (4h)) or 8-hour (TRAP@36°C (8h)) exposure to 36°C, stained with antibodies against HTB-GFP, cFos and DAPI. Scale bar: 100 μm. Right top: Increasing the time of exposure to 36°C increases the number of TRAPped GFP-positive cells in the VMPO. Kruskal-Wallis test, P <0.0001; Dunn’s multiple comparisons test, *P = 0.0110 (TRAP@RT: TRAP@36°C (4h)), ***P < 0.0001 (TRAP@RT : TRAP@36°C (8h)), **P = 0.0047 (TRAP@36°C (4h) : TRAP@36°C (8h)). n = 7 sections / 3 animals for TRAP@RT, n = 22/5 for TRAP@36°C (4h) and n = 18/4 for TRAP@36°C (8h). Right bottom: TRAPped neurons were found to highly overlap with endogenously expressed Fos protein in animals exposed to warmth; n = 22/5 for TRAP@36°C (4h) and n = 18/4 for TRAP@36°C (8h). Boxplots show median and interquartile range. b, Representative images using fluorescent confocal imaging of warm-stimulated (4 hours) and RT control mouse brain sections of LepR-Cre;HTB mice, revealing the rostral POA and MnPO and stained with antibodies against GFP (indicating LepR expression) and cFos; shown are the individual black and white images for DAPI, GFP and cFos and the 2-color composite images (GFP: green; cFos: red) revealing partial overlap of LepR and cFos as a consequence of challenging mice with warm temperatures and quantified in the graph shown in the right panel (displayed as % of all LepR-positive neurons and quantified from 11 and 10 sections of N = 2 animals for RT and 36°C, respectively; two-tailed T-test, p <0,0001, boxplot shows median and interquartile range; scale bars: 100 μm). c, Expression of Pacap (Adcyap) and Bdnf transcripts assessed by bulk mRNA sequencing of FACS sorted LepR^+^ and LepR^-^ cells obtained from POA tissue isolated from LepR-Cre;HTB mice. DESeq2 software package was used for differential expression analysis; LepR^+^: LepR^-^ ***P < 0.0001; n=18 (Wald test p<0,0001, boxplot shows median and interquartile range); each data point represents expression of the respective gene in each sample plotted as a log2 normalized value. d, rAAV(DJ)-DIO-ChR2-mCherry was injected and optic fiber implanted in the POA of LepR-Cre mice for optogenetic stimulation. Core body temperature, BAT and tail temperature of ChR2-expressing and control mice before, during (blue shading) and after blue light stimulation (20 Hz, 10 msec pulses, 1min ON/3min OFF) were measured. n=4 mice per group. Data represent mean ± s.e.m. e, Right: Quantification of the number of neurons expressing cFos (visualised by immunohistochemistry) in VMPO of LepR-Cre;HTB kept at room temperature (RT), and stimulated at 36°C for 2 or 4 hours and displayed as % of all cFos-positive neurons. Kruskal-Wallis test, P < 0.0001; Dunn’s multiple comparisons test, **P = 0.0047 (RT : 2h @ 36°C), ***P < 0.0001 (RT : 4h @ 36°C), **P = 0.0034 (2h @ 36°C : 4h @ 36°C). n = 7 sections / 2 animals for RT, 8 sections / 3 animals for 4h @ 36°C, and 6 sections / 2 animals for 8h @ 36°C. Left: Based on the same tissue sections, quantification of the absolute number cFos-positive cells. Kruskal-Wallis test, P = 0.0442; Dunn’s multiple comparisons test, P = 0.7224 (RT : 2h @ 36^C), P = 0.1709 (RT : 4h @ 36°C), *P = 0.0381 (2h @ 36°C : 4h @ 36°C). Boxplots show median and interquartile range. f, The extent of preoptic LepR+ neuron population. Representative images of the 250 µm acute slices from LepR-Cre;HTB mice used for *ex vivo* electrophysiological experiments. The majority of neurons for all VMPO electrophysiological recordings were recorded in slices shown in the middle panel (approx. bregma 0.26 mm). Scale bar: 100 μm. Neuronal activity was recoded in brain slices under fast synaptic transmission blockade and using “high-K^+^ aCSF” except for panels (c) and (e) where “low-K^+^ aCSF” and a 33°C bath temperature was used.

**Extended Data Fig. 2.**
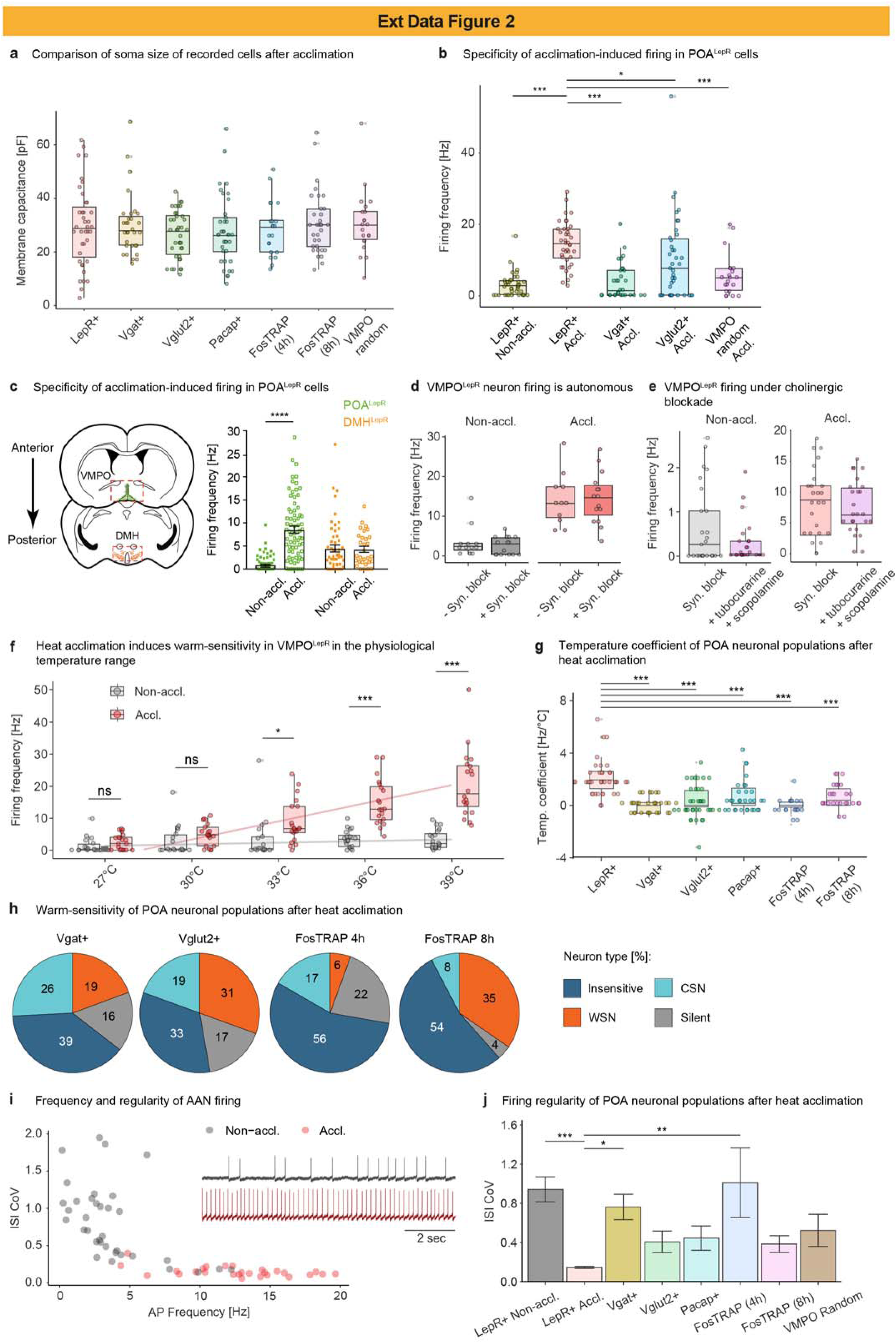
Heat acclimation-induced upregulation of tonic warm-sensitive AP firing is most robustly detected in the VMPO^LepR^ population. a, Comparison of membrane capacitances of acclimated neurons from various VMPO populations recorded in acute slices. Within all the recorded neuronal populations, cells in similar soma size range were selected for measurements. b, Comparison of spontaneous AP frequency at 36°C in acclimated VMPO^LepR^ population to AP frequencies observed in other VMPO neuronal populations tested. On average, acclimated VMPO^LepR^ neurons had a significantly higher firing frequency compared to acclimated VMPO^Vgat^ neurons, acclimated VMPO^Vglut2^ neurons as well as randomly sampled VMPO acclimated neurons. Non- acclimated VMPO^LepR^ neurons plotted for reference. One-way ANOVA, P < 0.0001; Sidak’s multiple comparison test, ***P < 0.0001 (LepR+ Accl. : LepR+ Non-accl.), ***P < 0.0001 (LepR+ Accl. : Vgat+ Accl.), *P = 0.0362 (LepR+ Accl. : Vglut2+ Accl.), ***P < 0.0001 (LepR+ Accl. : VMPO random Accl.). n = 40/7 (LepR+ Accl), n = 40/6 (LepR+ Non accl), n = 31/3 (Vgat+ Accl.), n = 39/4 (Vglut2+ Accl.) and n = 21/3 (VMPO Random Accl.) cells. c, Cartoon depicting the location of VMPO^LepR^ neurons (green) and DMH^LepR^ neurons (orange) recorded in whole-cell patch clamp configuration to assess specificity of acclimation-induced increases in AP firing frequency. Firing frequency (Hz) was recorded in both neuronal populations, in brain slices prepared from acclimated and non-acclimated animals (n=35/5 per group). Kruskal-Wallis test, P<0.0001; Dunn’s multiple comparisons test, p<0.0001 (Non-Accl. : Accl.POA LepR). No change in the average AP firing frequency was observed in DMH^LepR^ neurons. d, AP firing rates at 36°C in both non-acclimated and acclimated VMPO^LepR^ neurons in *ex vivo* brain slices are comparable in the presence and absence of the fast synaptic transmission blockers (CNQX 10 µM, APV 50 µM and Gabazine 5 µM). n = 12/2 (Non-accl. –Syn. block), n = 15/3 (Non-accl. +Syn. block), n = 11/2 (Accl. –Syn. block), n = 15/3 (Accl. +Syn. block) cells. e, Addition of the acetylcholine receptor antagonists tubocurarine (10 µM) and scopolamine (10 µM) to the external solution containing fast synaptic transmission blockers (CNQX, APV and Gabazine; 10, 50 and 5 µM respectively) slightly (but insignificantly) reduced AP firing in non-acclimated VMPO^LepR^ neurons but does not modulate AP firing in acclimated VMPO^LepR^ cells *ex vivo*. n = 25/5 (Non-accl. / syn. block), n = 25/4 (Non-accl. / syn. block+tubocurarine+scopolamine), n = 25/6 (Accl. / syn. block), n = 25/4 (Accl. / syn. block+tubocurarine+scopolamine). Data recorded at 33°C. f, Firing frequencies of VMPO^LepR^ at sub-physiological and physiological temperatures: regression analysis (represented here by grey and red regression lines) of non-acclimated and acclimated VMPO^LepR^ based on firing rates measured at 33°C, 36°C and 39°C (previous data shown here and in main Fig. 1e). The extension of these regression lines into the colder temperature range predicted that firing of acclimated and non-acclimated neurons should become indistinguishable at ∼29.1°C (crossing point of the red and grey regression line), which was subsequently confirmed experimentally, by recording firing rates at 27°C and 30°C. Data partially overlapping with Fig 1e. ns: not significant. g, Warm sensitivity (measured by the temperature coefficient Tc) of tested VMPO neuronal populations after heat acclimation. One-way ANOVA, P < 0.0001; Tukey’s multiple comparison test, ***P < 0.0001 (LepR+ : Vgat+), ***P < 0.0001 (LepR+ : Vglut+), ***P < 0.0001 (LepR+ : Pacap+), ***P < 0.0001 (LepR+ : FosTRAP 4h), ***P < 0.0001 (LepR+ : FosTRAP 8h). n = 38/7 (LepR+), n = 31/3 (Vgat+), n = 36/4 (Vglut+), n = 31/3 (Pacap+), n = 18/2 (FosTRAP 4h) and n = 26/3 (FosTRAP 8h) cells. h, Distribution of *ex vivo* recorded temperature-insensitive, cold-sensitive (CSN, temperature coefficient < -0.6 Hz/°C), warm-sensitive (WSN, temperature coefficient ≥ 0.75 Hz/°C) and silent neurons within the acclimated VMPO neuronal populations demarcated by the expression of Vgat (n = 31/3) and Vglut2 (n = 36/4) as well as “warm TRAPped” neurons for either 4h (n = 18/2) or 8h (n = 26/3). Compare with Fig. 1d. i, Spontaneous activity pattern in representative non-acclimated and acclimated VMPO^LepR^ differs not only by frequency but also regularity of action potential firing as evidenced by the different inter spike interval coefficient of variation (ISI CoV, a measure of AP firing regularity, is the standard deviation of the interspike interval (ISI) divided the mean ISI). Analysed recordings were performed at 36°C bath temperature. j Interspike interval coefficient of variation (ISI CoV) for the indicated neuronal populations obtained from heat acclimated mice. Neuronal activity was recoded in brain slices under fast synaptic transmission blockade and using “high- K+ aCSF” except for panels (c) and (e) where “low-K+ aCSF” and 33°C bath temperature were used.

**Extended Data Fig. 3.**
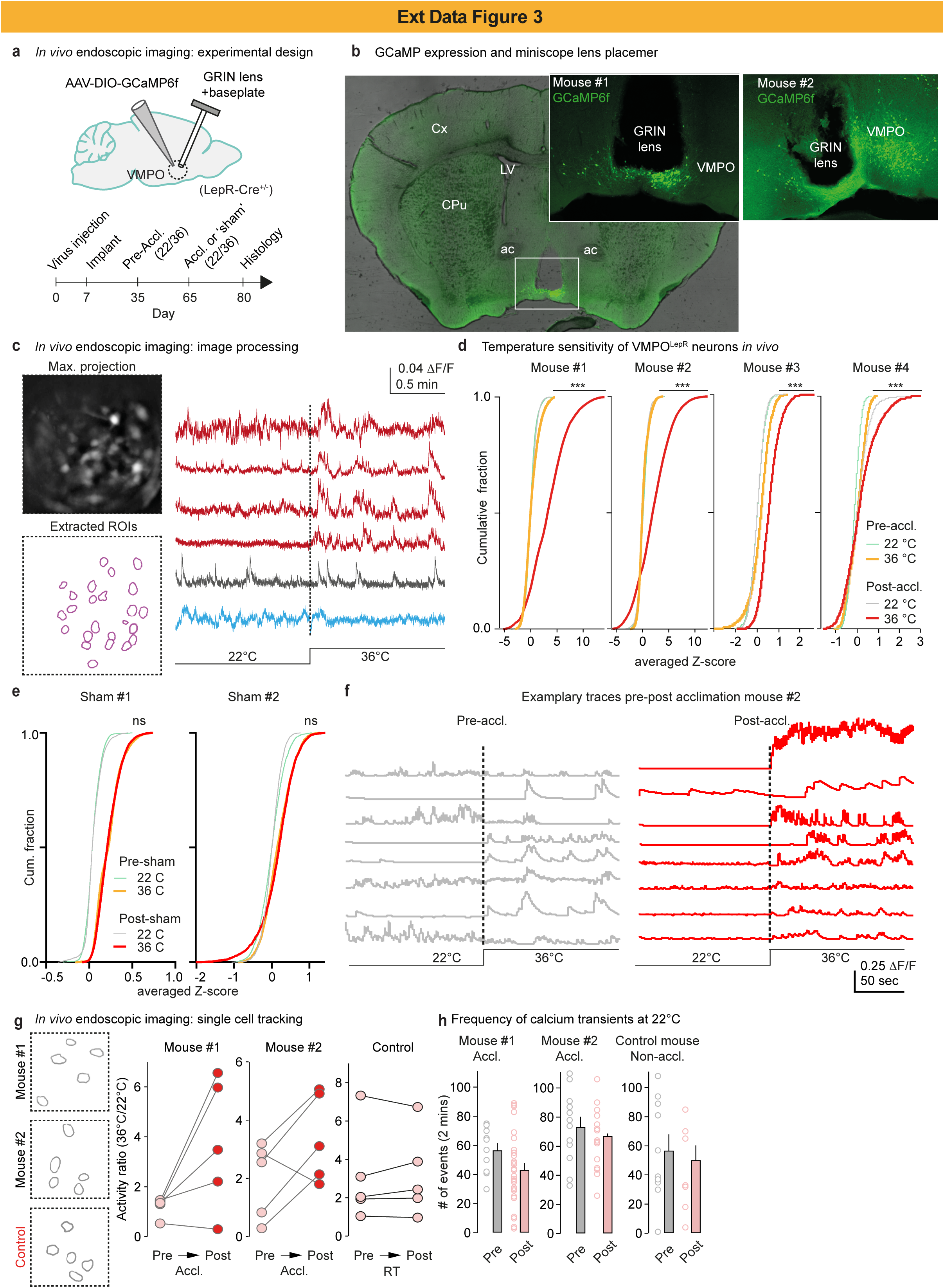
Microendoscopy reveals acclimation-induced VMPO^LepR^ warm responsiveness *in vivo*. a, Top: schematic of experimental configuration indicating AAV-mediated GCaMP6f delivery into VMPO^LepR^ neurons and GRIN lens implantation. Bottom: experimental timeline. b, Representative images showing GCaMP6f expression in VMPO^LepR^ neurons, and location of the GRIN lens implant used to gain visual access to analyze VMPO activity. c, Left: maximal projection image from a representative imaging session and location of the regions of interest (ROI) extracted for analysis. Right: calcium dynamics from 6 representative neurons recorded from a non-acclimated mouse at RT (22°C) and at 36°C ambient temperature. Note that some cells increase (red traces), decrease (blue trace) and don’t change (grey trace) activity upon temperature increase. d, Cumulative distribution plots of the activity (averaged z-scores) of all extracted neurons in 4 mice upon increasing ambient temperature acutely from RT (22°C) to 36°C, before and after acclimation for 30 days at 36°C. Mann-Whitney U test, ***P < 0.0001 for cumulative fraction at 36°C Pre vs Post acclimation. e, Cumulative distribution plots of the activity (averaged z-scores) of all extracted neurons in 2 time-matched control mice upon increasing ambient temperature acutely from RT to 36°C, before and after sham acclimation. The time-matched controls did not undergo acclimation to temperature but stayed at at 22°C for 30 days in between recording sessions. Mann-Whitney U test, P = 0.2273 (Sham #1) and P = 0.3624 (Sham #2) for cumulative fraction at 36°C Pre vs Post sham acclimation. f, Exemplary traces of calcium dynamics (DF/F) in 8 randomly picked cells of mouse #2 from panel d at 22°C and 36°C before heat acclimation (Pre-accl.), and other 8 randomly picked cells from the same mouse at 22°C and 36°C after heat acclimation (Post-accl.). g, Representative VMPO^LepR^ that could be reliably tracked and recorded before and after acclimation (30 days in between recording sessions) in 2 acclimated mice, and in a time-matched RT control mouse (recorded at the same time intervals but without undergoing heat acclimation). Activity of single cells upon changes in ambient temperature from 22°C to 36°C. Cells were longitudinally recorded before (pre) and after (post) temperature acclimation. h, Summary plots of calcium transients (number of events detected during a 2 mins recording session) in two mice (left, middle) before and after heat acclimation with the mouse kept at 22°C during recordings and a non-acclimated control mouse with matched inter-recording interval in between recording sessions (30 days) without heat acclimation (right); related to panel h. N=4 acclimated animals, N=2 time-matched non-acclimated animals.

**Extended Data Fig. 4.**
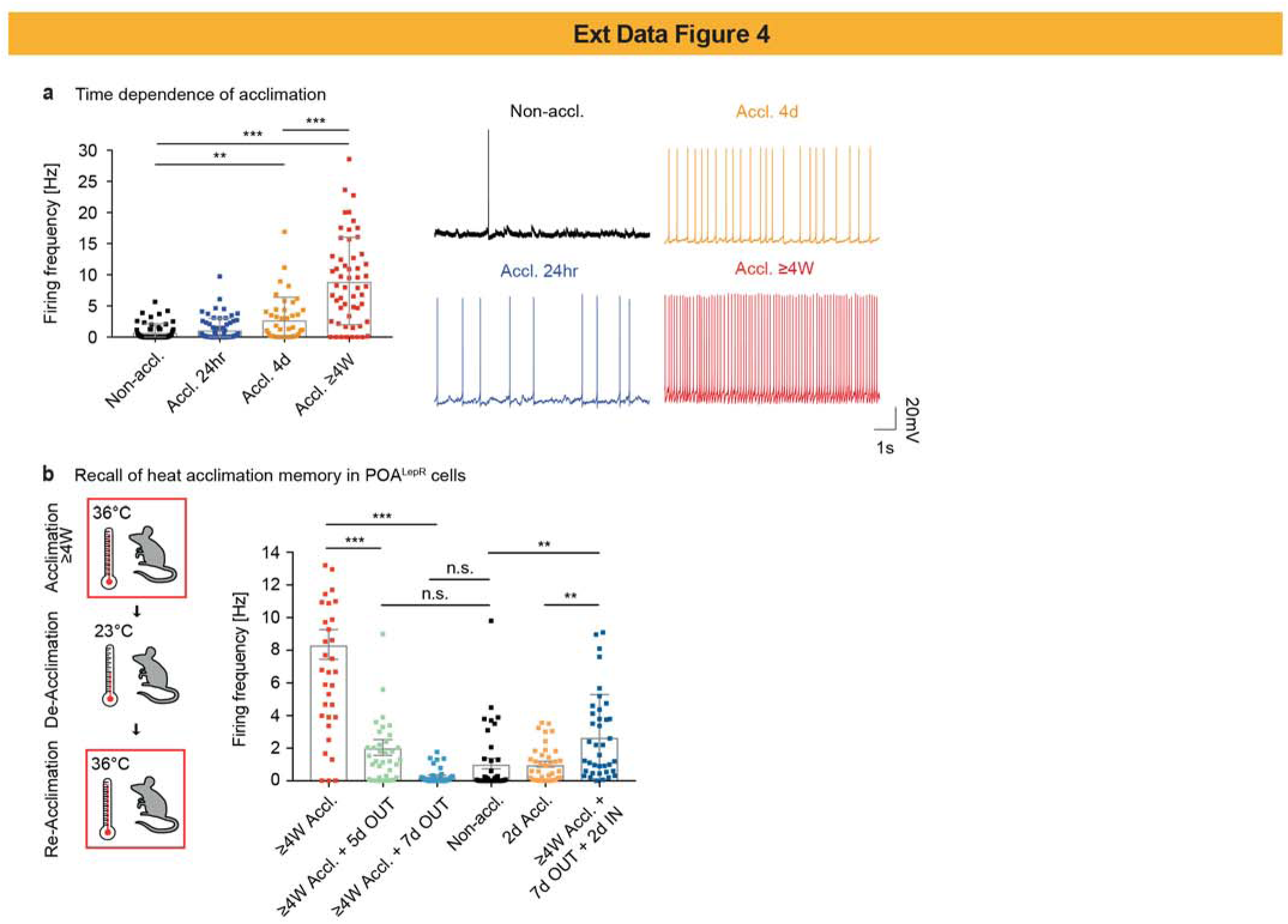
Kinetics of VMPO^LepR^ acclimation, deacclimation and reacclimation. a, Left: AP firing frequencies of VMPO^LepR^ neurons recorded *ex vivo* from non-acclimated mice as well as mice acclimated for 24 hours, 4 days and 4 weeks (full acclimation). Kruskal-Wallis test, P <0.0001; Dunn’s multiple comparisons test, **P = 0.0062 (Non-accl. : Accl. 4d), ***P < 0.0001 (Non-accl. : Accl. ≥4W), ***P = 0.0005 (Accl. 4d:Accl. ≥4W). n = 42/5 per group. Right: Representative traces of AP firing patterns as a function of heat acclimation duration, recorded in VMPO^LepR^ neurons. Brain slices were recorded at 33°C bath temperature. b, AP firing frequency (Hz) was measured in VMPO^LepR^ neurons subsequent to subjecting LepR- Cre;HTB mice to different acclimation, de-acclimation and re-acclimation periods: Non-acclimated control (Non-accl., black), two-day acclimation (2d Accl., orange), full acclimation (≥ 4 weeks Accl., red), five or seven days of de-acclimation after full acclimation (≥ 4 weeks Accl. + 5d OUT, green and ≥4 weeks Accl. + 7d OUT, light blue, respectively) or re-acclimation after removing fully (4-5 weeks) acclimated animals for 7 days from the 36°C acclimation chamber to RT and re-acclimating them for only 2 days at 36°C (≥ 4 weeks Accl. + 7d OUT + 2d IN, dark blue). Subsequent to full acclimation (4- 5 weeks), AP firing returned to baseline after 7 days of de-acclimation. Subsequent re-acclimation for only 2 days, substantially elevated AP firing to high levels, much higher than short 2-day acclimation of naïve animals could accomplish. One-way ANOVA, P < 0.001; Sidak’s multiple comparisons test, ***P < 0.0001 (≥ 4W Accl. : ≥ 4W Accl. + 5d OUT), ***P < 0.0001 (≥ 4W Accl. : ≥ 4W Accl. + 7d OUT), **P = 0.0061 (2d Accl. : ≥ 4W Accl.+ 5d OUT + 2d in), **P = 0.0061 Non-accl. : ≥ 4W Accl.+ 5d OUT + 2d in), **P = 0.004 (2d Accl. : ≥ 4W Accl.+ 7d OUT + 2d IN), ***P = 0.0002 (Non-accl. : ≥ 4W Accl.+ 7d OUT + 2d IN). n= 38/3 cells per group. Spontaneous neuronal activity was recoded in brain slices without synaptic blockade, using “low-K^+^ aCSF” and at 33°C bath temperature.

**Extended Data Fig. 5.**
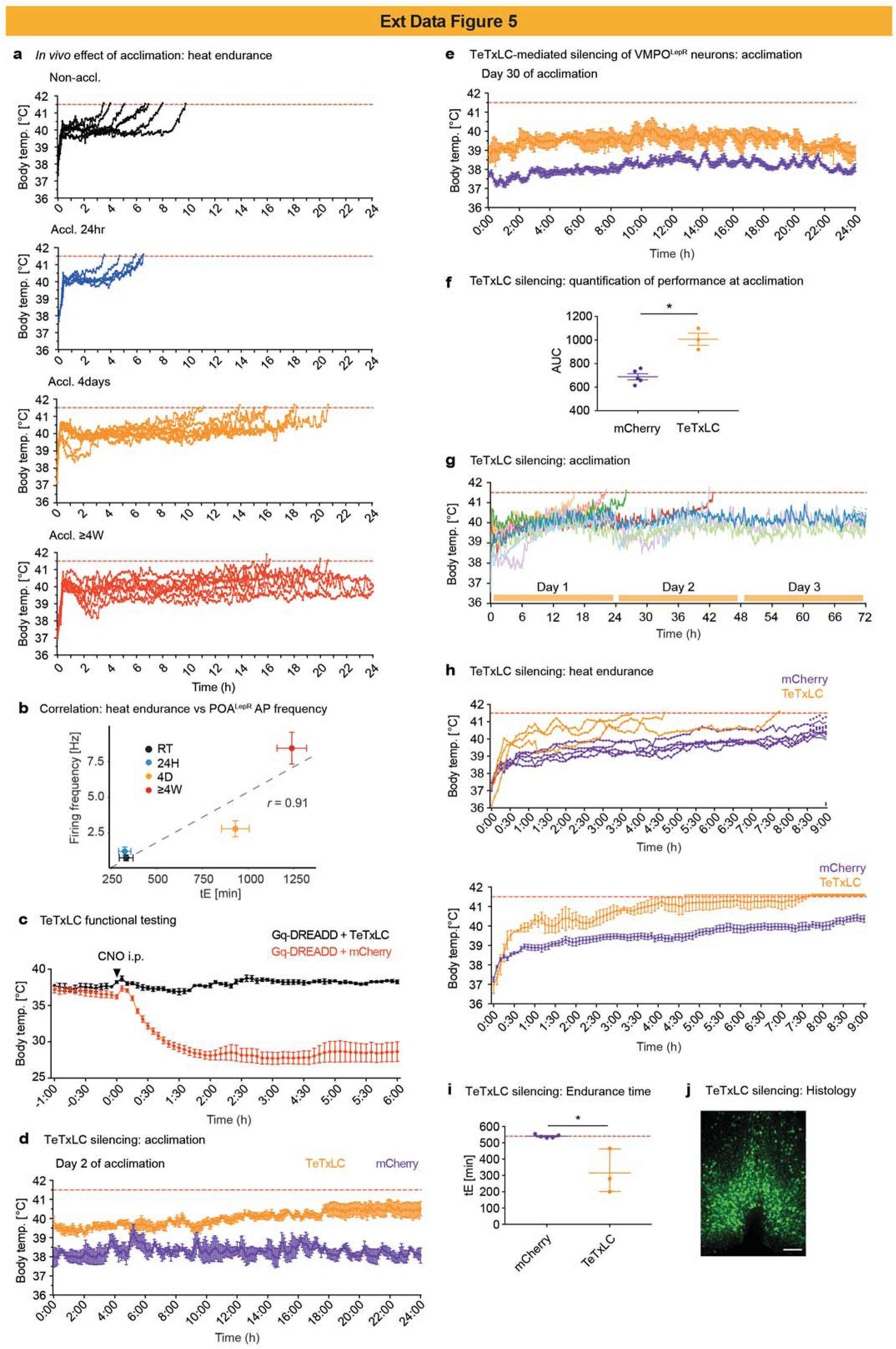
Heat acclimation-induced heat tolerance is blocked by TeTxLC-mediated VMPO^LepR^ silencing. a, Heat endurance assay: Body temperature of individual non-acclimated (Non-Accl., black, N = 7) animals, 24hr-acclimated (Accl. 24hr, blue, N = 5) animals, 4 days-acclimated (Accl. 4days, orange, N = 8) animals, and ≥ 4 weeks acclimated (Accl. ≥ 4W, red, N = 7) acclimated mice during 24-hour heat endurance assay (39°C ambient temperature). Animals that reached the cut-off temperature of 41.5°C, demarcated by the dashed red line, were discontinued from the assay. b, Correlation plot between heat endurance time (tE) and average firing frequency of POA^LepR^ neurons after varying duration of acclimation. Pearson’s (r) correlation coefficient between the two parameters is shown. c, TeTxLC functionality was tested *in vivo* by measuring body temperature following i.p. injection of CNO in mice where VMPO^LepR^ neurons expressed either Gq-DREADD or Gq-DREADD+TeTxLC (N = 4 per group). While animals that did not express TeTxLC displayed a decrease in body temperature following CNO administration, co-expression of TeTxLC completely abrogated the response. d, e, Average body temperature of TeTxLC- and mCherry (control)-infected animals at day 2 (d) and at day 30 (e) of heat acclimation, showing that TeTxLC-animals are hyperthermic throughout the acclimation phase. Note that during heat acclimation 6 of the 9 TeTxLC-animals dropped out (but none of the ctrl animals). f, Quantification of the area under the curve (AUC) calculated for the two groups for the last day of acclimation. Mann-Whitney U test, *P = 0.0286. N = 3 mice for TeTxLC and N = 5 mice for mCherry control. g, Body temperature traces of individual TeTxLC-silenced LepR-Cre animals are shown for the first 3 days of heat acclimation (36°C). 6 out of 9 animals with silenced VMPO^LepR^ neuron outputs failed during the first two days of acclimation and reached the Tcore cut-off of 41.5°C (demarcated by dashed red line). h, TeTxLC-silenced animals that completed the 30-day acclimation cycle (N = 3) were tested side-by-side with acclimated mCherry-injected control animals (N = 5) in the heat endurance assay. Top: Body temperature traces of individual mice. Bottom: average body temperature traces for the TeTxLC-expressing and mCherry-expressing control groups. i, Quantification of the endurance time (tE) in the 9-hour (540 min) heat endurance assay for the two groups. Mann-Whitney U test; *P = 0.0262 (mCherry : TeTxLC). N = 3 for TeTxLC and N = 5 for mCherry control mice. j, Representative image of VMPO^LepR^ neurons labelled with that is EGFP co-expressed with TeTxLC; Size bar = 250 μm.

**Extended Data Fig. 6.**
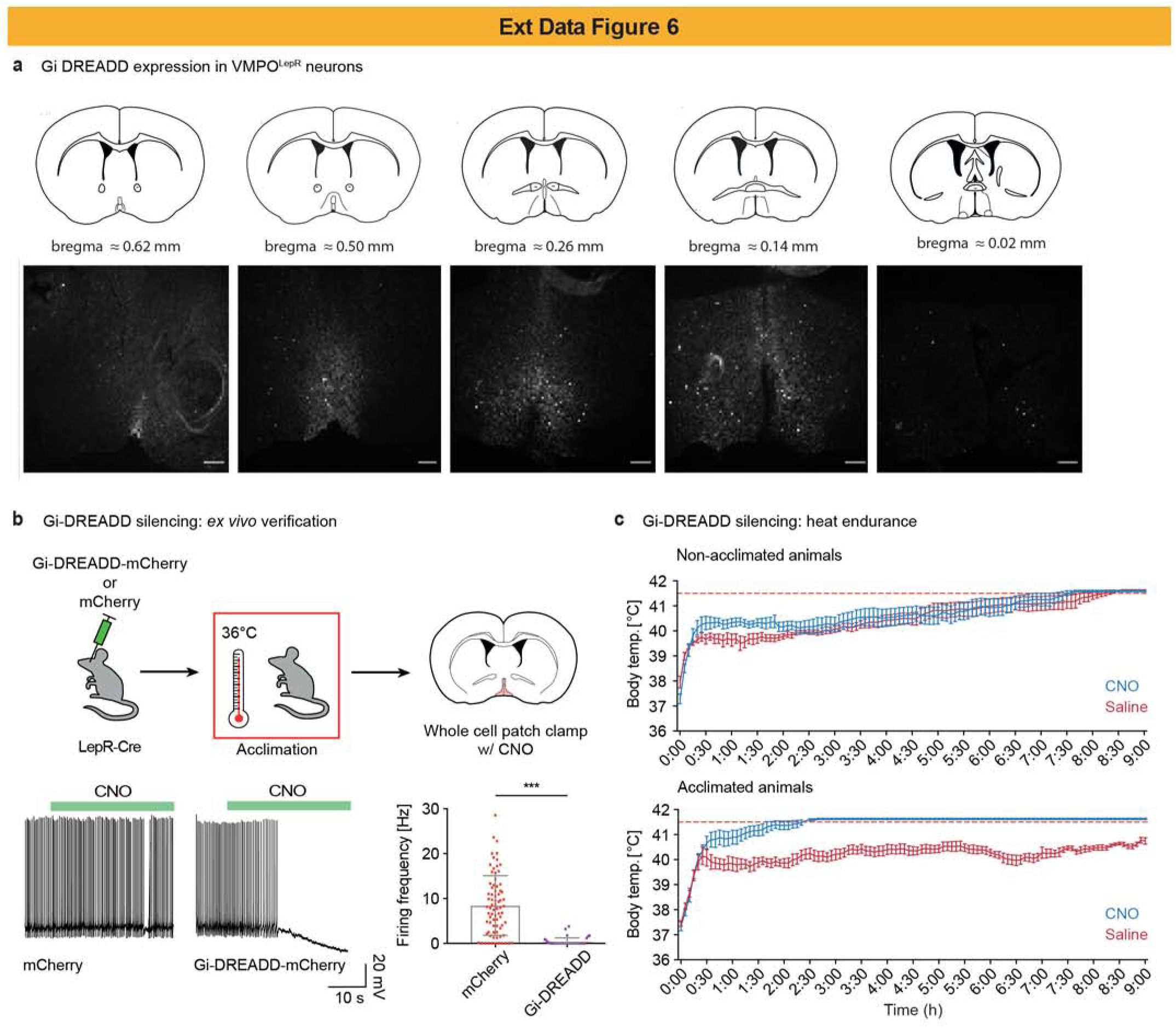
Gi-DREADD-driven inhibition of acclimation-induced VMPO^LepR^ activity prevents heat tolerance. a, Schematic drawings representing different anatomical positions along the rostral caudal axis of the preoptic hypothalamic region with the 3 middle drawings (approx. bregma = 0.5 mm to bregma = 014) indicating the center of the VMPO region (top) with corresponding typical fluorescent images depicting the extent of virally (AAV) delivered Cre-dependent Gi-DREADD expression in a LepR-Cre mouse (bottom). b, Top: Schematic showing the protocol used for *ex vivo* verification of CNO triggered, Gi-DREADD mediated inhibition of VMPO^LepR^ following heat acclimation. Bottom left: Representative electrophysiological traces showing the effect of CNO on the firing pattern of acclimated VMPO^LepR^ neurons injected with either Cre-dependent Gi-DREADD-mCherry AAV or only a Cre-dependent mCherry control AAV. Bottom right: average (mean ± s.e.m.) tonic AP firing frequency of acclimation- induced VMPO^LepR^ cells in the presence of 5 µM CNO. Mann-Whitney U test, ***P < 0.0001. n = 35/3 cells per group. c, Heat endurance assay: Average (mean ± s.e.m.) body temperature of non-acclimated (top) or acclimated (bottom) Gi-DREADD-positive and CNO-injected animals during the heat endurance assay. The same animals injected with saline instead of CNO were also plotted for comparison. N = 8 mice for the non-acclimated and N = 7 mice for the acclimated condition.

**Extended Data Fig. 7.**
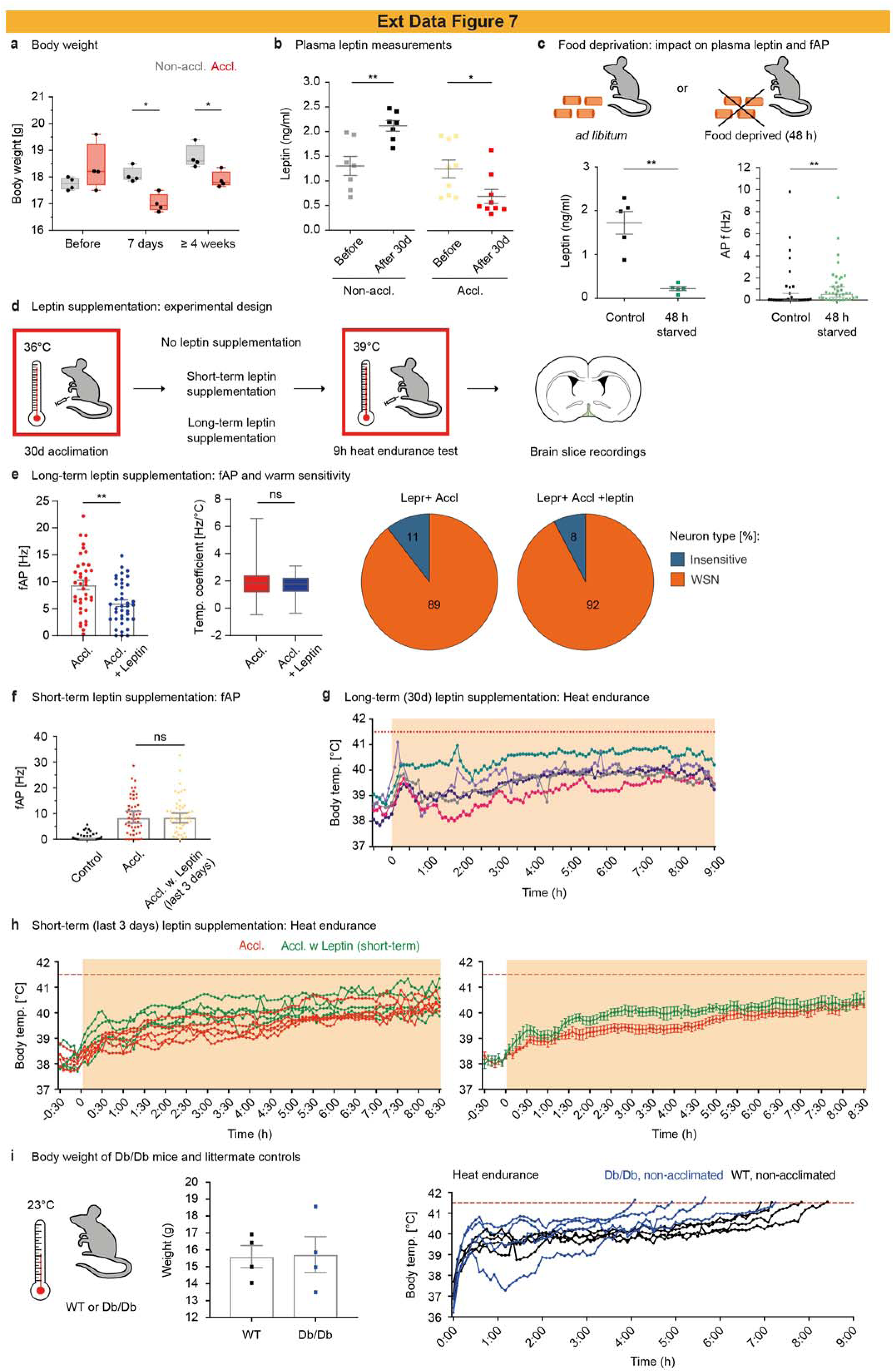
Effect of leptin signaling on VMPO^LepR^ activity and performance on heat acclimation and heat endurance. a, Body weight of acclimated animals decreased and remained significantly lower compared to that of non-acclimated counterparts during the entire acclimation period. Two-way ANOVA (effect of acclimation time * treatment), P = 0.0046; Tukey’s multiple comparisons test; *P = 0.0177 (7 days, Non-accl. : Accl.), *P = 0.0366 (≥4 weeks, Non-accl. : Accl.). N=4 per group. b, Blood plasma leptin measurements in non-acclimated and acclimated animals over the course of 30 days. Mann-Whitney U test, **P = 0.0070 for the non-acclimated condition and *P = 0.0142 for the acclimated condition. N = 7 for Non-accl. and N = 9 for Accl. c, Leptin content in the blood as well as frequency of action potential firing of VMPO^LepR^ neurons were tested upon food-depriving LepR-Cre;HTB animals for 48 h. Ad libitum-fed mice served as control group. Mann-Whitney U test, **P = 0.0079 for Leptin concentration and **P = 0.0050 for fAP. d, LepR-Cre;HTB animals were twice daily injected intraperitoneally with leptin during the last three days of acclimation (short term) or during the entire 30-day acclimation period (long-term). Control groups received injections containing saline. Following acclimation, animals were tested in 9-hour heat endurance assay. Following the assay, the activity of VMPO^LepR^ neurons was recorded in brain slices *ex vivo*. e, Effect of long-term supplementation of leptin during the heat acclimation process on tonic VMPO^LepR^ activity. Left: long-term leptin treatment slightly reduced fAP (recorded at 36°C) in acclimation- induced VMPO^LepR^. Unpaired two-tailed t-test, **P = 0.0035. n = 38/5 (Accl.) and n = 39/3 (Accl. + Leptin) cells. Middle: long-term supplementation of leptin did not have an effect on warm-sensitivity of VMPO^LepR^ neurons, measured by temperature coefficient. n = 38/5 (Accl.) and n = 39/3 (Accl. + Leptin) cells. Right: Distribution of temperature-insensitive, and warm-sensitive (WSN) within acclimated VMPO^LepR^ control group (n = 38/5 cells) and VMPO^LepR^ group after 30 days of leptin supplementation (39/3 cells). Recordings were performed using “high-K^+^ aCSF” and in the presence of synaptic blockers CNQX, AP-V and gabazine. f, Effect of short-term supplementation of leptin (last three days of heat acclimation) on VMPO^LepR^ activity. Short-term leptin treatment did not have any impact on fAP in acclimated VMPO^LepR^; non- acclimated VMPO^LepR^ group was plotted for visual comparison. n = 51/5 (Accl.) and n = 51/5 (Accl. + Leptin 100 nM in aCSF) cells. g, Heat endurance assay: Body temperature traces of LepR-Cre;HTB animals during the heat endurance assay (39°C) following long-term supplementation of leptin during 30d of heat acclimation at 36°C. N = 5 mice. h, Heat endurance assay: Body temperature traces of individual (left) LepR-Cre;HTB animals following short-term supplementation of leptin (last three days of heat acclimation) during 39°C heat challenge. Control group consisted of acclimated animals injected with saline. Group averages are presented in the right panel. N = 5 animals each. i, Db/db animals were pair-fed with littermate control mice for 1 week and kept at room temperature (22°C/23°C) before undergoing the 9-hour heat endurance assay. Middle: body weight was comparable between the two groups prior to the assay. Right: Both control and Db/Db mice reached the body temperature cut-off of 41.5°C prior to the conclusion of the 9-hour period. N = 4 animals per group. Tonic neuronal activity in panel (f) was recoded in brain slices without synaptic blockade, using “low- K+ aCSF” and at 33°C bath temperature.

**Extended Data Fig. 8.**
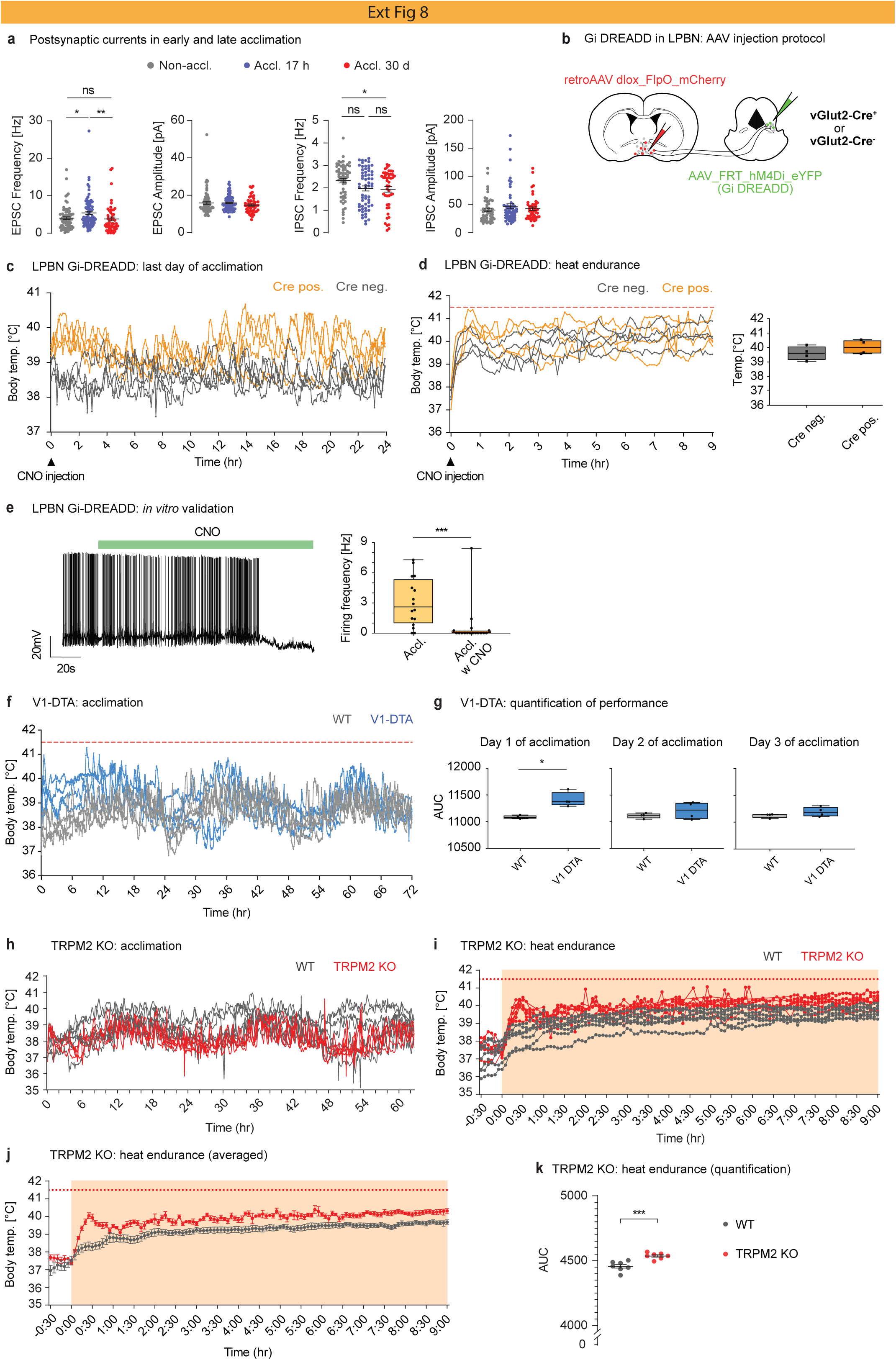
Thermo-afferent pathways via the PBN are required during the initial phase of heat acclimation but appear to become obsolete at late acclimation stages. a, Spontaneous excitatory (EPSC) and inhibitory (IPSC) postsynaptic currents in VMPO^LepR^ neurons recorded in slices of non-acclimated, short-term acclimated (17 hours) and long-term acclimated (30 days) LepR-Cre;HTB mice. Left: EPSC frequency was found to be upregulated after 17 hours of acclimation compared to the non-acclimated condition, and to decrease again to baseline levels after long-term acclimation. Kruskal-Wallis test, P = 0.0017; Dunn’s multiple comparisons test, *P = 0.0328 (Non-accl. : Accl. 17h), **P = 0.0024 (Accl. 17h : Accl. 30d). n = 74/4 (Non-accl.), n = 82/4 (Accl. 17h) and n = 51/4 (Accl. 30d) cells. 2^nd^ from left: EPSC amplitude did not change between the conditions tested. 3^rd^ from left: IPSC frequency was found to decrease over the course of acclimation. One-way ANOVA, P = 0.0345; Tukey’s multiple comparison test, *P = 0.0497 (Non-ccl.:Accl. 30d). n = 57/3 (Non-accl.), n = 60/3 (Accl. 17h) and n = 46/3 (Accl. 30d) cells. 4^th^ from left: IPSC amplitude did not change between the conditions tested. b, Strategy for Gi-DREADD delivery into glutamatergic LPBN neurons that target VMPO using the Vglut2-Cre mouse line. c, Gi-DREADD mediated inhibition of PBNèPOA projection neurons renders Cre-positive animals (orange), but not Cre-negative controls (grey), slightly hyperthermic when CNO was injected at the very end of the 30-day acclimation phase. d, CNO-mediated inhibition of these PBNèPOA projection neurons during the heat endurance assay did not perturb heat tolerance. Left: Body temperature traces of individual Cre-positive (orange) and Cre-negative (grey) mice during the 9 hour heat endurance assay. Right: Boxplot showing the average body temperature of mice expressing Gi-DREADD and AAV-injected controls during the assay. e, Representative electrophysiological traces showing the effect of CNO on the firing pattern of Gi-DREADD-expressing PBN neurons recorded from an acclimated mouse *ex vivo*. Right panel: average AP firing frequency (n=17) of control and CNO treated cells. Mann-Whitney U test; ***P < 0.0001. n = 17/2 cells each. Boxplots show median and interquartile range. f, Body temperature of individual V1-DTA ablated and wildtype littermate control animals (N = 4 per group) during the first three days of heat acclimation. g, Area under the curve (AUC) calculated from body temperature recordings for three consecutive days of acclimation (Day 1 = 0-24 hr, Day 2 = 24-48 hr and Day 3 = 48-72 hr) for the V1-DTA ablated and control mouse groups. Mice lacking major peripheral somatosensory thermal signals were hyperthermic only initially but adapted to acclimation temperatures similar to control animals but with a delay of 1 to 2 days. Mann-Whitney U test; *P = 0.0286. N = 4 animals per group. h, Body temperature traces of individual TRPM2-KO and control animals during the first 3 days of acclimation. N = 7 mice each. i, j, Individual (i) and average (j) body temperature traces TRPM2-KO and control animals during the heat endurance assay. N = 7 mice per group. k, Quantification of the area under curve (AUC) for TRPM2-KO and control animals during the heat endurance assay. TRPM2 KO animals did not reach the 41.5°C threshold but had a significantly elevated body temperature throughout the assay compared to control mice. Unpaired two-tailed t-test, ***P = 0.0006. N = 7 mice per group.

**Extended Data Fig. 9.**
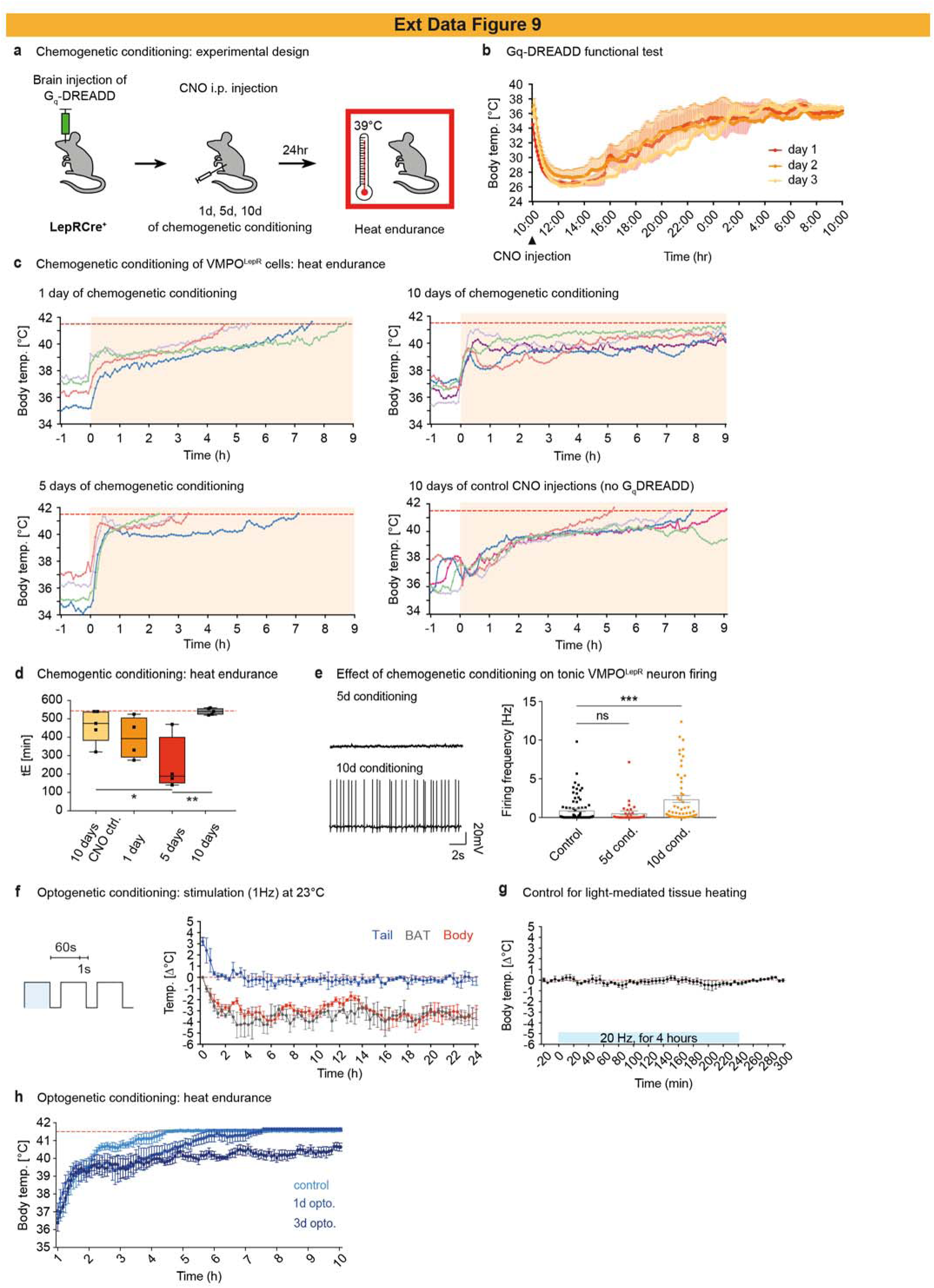
Long-term activation of VMPO^LepR^ neurons by chemogenetic (Gq-DREADD) or optogenetic (ChR2) conditioning is sufficient to induce heat tolerance in the heat endurance assay. a, Cartoon showing experimental paradigm: viral injection of Cre-dependent Gq-DREADD viral particles into the rostral POA of LepR-Cre mice. VMPO^LepR^ neurons were chemogenetically activated (conditioned) by daily injection of CNO (0.3 mg/kg i.p.) for 1, 5 or 10 consecutive days. CNO injections were terminated 24 hours prior to heat endurance assay. b, Chemogenetic conditioning of VMPO^LepR^ cells via Gq-DREADD animals produced significant decrease in body temperature (hypothermia) that is protracted for up to 10 hours after CNO injection and could be repeated over multiple consecutive days. Traces represent group average (mean ± s.e.m.) for each day of CNO injection. N = 4 animals. c, Body temperature traces of individual chemogenetically conditioned mice during the heat endurance assay. All animals that were chemogenetically conditioned for 10 days passed the heat endurance assay for more than 9 hours. Animals that reached the cut-off temperature of 41.5°C, demarcated by the dashed red line, were discontinued from the assay. d, Boxplots (median and interquartile range) showing the endurance times (tE) before a body temperature of 41.5°C was reached, corresponding to data shown in (c). Note that the maximum tE attainable for all animals was at 9 hours (540 min, demarcated by red dashed line). All animals that were chemogenetically conditioned for 10 days reached the maximum tE, suggesting that the heat tolerance potential of these mice was underestimated in this assay. Kruskal-Wallis test, P = 0.0370; Sidak’s multiple comparison test, *P = 0.0312 **P = 0.0034. N = 5 for Control (animals that received saline injections for 10 days), N = 4 for “1 day” (a single CNO injection), N = 4 for “5 day” (5 days of CNO injections) and N = 5 for “10 day” (10 days of CNO injections). e, Left: Representative traces of AP firing patterns of two VMPO^LepR^ neurons recorded 24 hours after chemogenetic conditioning for 5 or 10 days. Right: average AP firing frequency of VMPO^LepR^ neurons from non-stimulated control LepR-Cre;HTB animals and from animals chemogenetically conditioned for 5 or 10 days. Kruskal-Wallis test; P < 0.0001; Dunn’s multiple comparisons test, ***P < 0.0001 (Control: 10d cond.). n = 42/4 cells per group. Neuronal activity were performed without synaptic blockade, using “low-K^+^ aCSF” and at 33°C. f, Average body-, brown adipose- (BAT-) and tail temperature of LepR-Cre mice stereotactically injected with viral particles into the rostral POA to express ChR2 in a Cre-dependent manner and stimulated with blue light at a low frequency of 1 Hz at room ambient temperature (23°C). g, Optogenetic control experiment: in the absence of ChR2, light stimulation of the POA/VMPO of up to 20 Hz continuously for 4 hours did not affect body temperature in freely moving mice. h, Average body temperature (mean ± s.e.m.) during heat endurance of LepR-Cre mice expressing ChR2 in VMPO^LepR^ neurons that were either not optogenetically conditioned (control, light blue trace), conditioned for 1 day (1d opto, blue trace) or 3 days (3d opto, dark blue trace). Note that all mice, including control animals, were optically stimulated with light pulses at 1 Hz during the heat endurance assay. N = 4 animals each.

**Extended Data Fig. 10.**
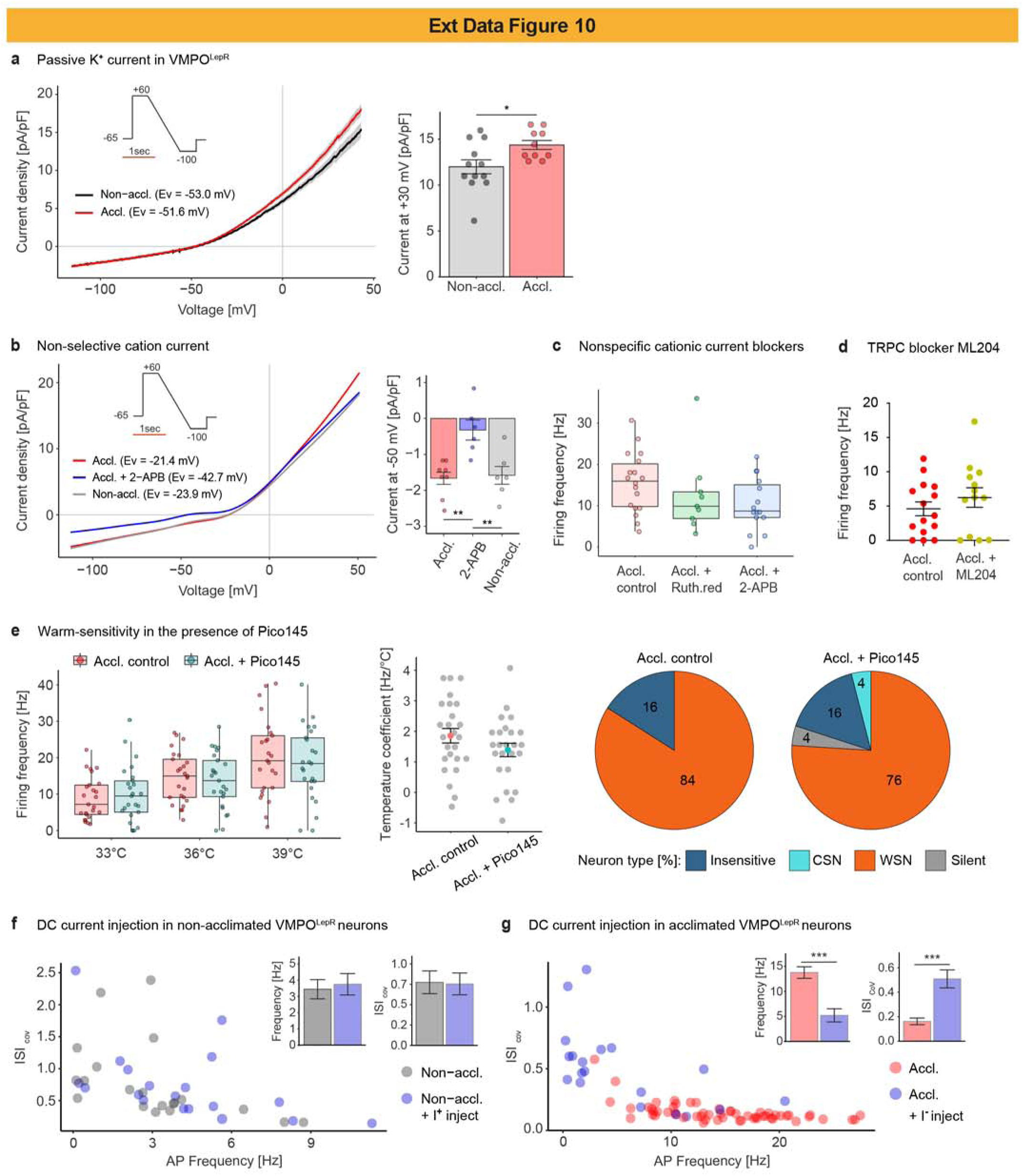
Characterization of background cation currents and their effect on VMPO^LepR^ neuron activity. a, Left: Passive K^+^ transmembrane conductance was slightly enhanced in acclimated VMPO^LepR^ compared to non-acclimated controls. Average reversal potentials (EV) are stated. Inset shows the hyperpolarising ramp protocol used. Right: Quantification of current amplitude at +30 mV based on presented traces. Unpaired two-tailed t-test, *P = 0.0208. n = 12/2 (Non-accl.) and n = 10/2 (Accl.) cells. Data represents mean ± s.e.m. b, Left: Passive, nonspecific cation transmembrane current was similar between non-acclimated and acclimated VMPO^LepR^ groups, but reduced by 2-APB (100 µM) in the acclimated condition. Traces represent group averages; average reversal potentials (EV) are stated. Inset shows the hyperpolarising ramp protocol used. Right: Quantification of current amplitude (mean ± s.e.m.) at -50 mV based on presented traces. One-way ANOVA, P = 0.0016; Tukey’s multiple comparison test, **P = 0.0013 (Accl. : Accl.+2-APB), **P = 0.0052 (Non-accl. : Accl.+2-APB). n = 8/1 (Accl.), n = 6/1 (Accl.+2-APB) and n = 6/1 (Non-accl.) cells. c, Inhibition of non-selective cation currents using the non-specific TRP channel blockers Ruthenium red (10 µM) and 2-APB (100 µM) did not have a significant effect on tonic firing frequency of acclimated VMPO^LepR^ neurons. n = 10/5 (Accl. control), n = 9/1 (Accl. + Ruth.red) and n = 15/2 (Accl. + 2-APB) cells. Boxplots represent median and interquartile range. d, Inhibition of TRPC channels with the blocker ML204 (10 µM) did not majorly affect the firing frequency of acclimated VMPO^LepR^ neurons. n = 15/1 (Accl. control), n = 13/1 (Accl. + ML204) cells. Data represent mean ± s.e.m. e, Blockade of TRPC1/4/5 channels did not affect the firing pattern nor warm-sensitivity of acclimation- induced VMPO^LepR^ neurons. Left: Boxplots (median and interquartile range) showing the frequency of AP firing in acclimated VMPO^LepR^ neurons without (red) and with the selective TRPC1/4/5 channel antagonist Pico145 (10 nM) at three bath temperatures. n = 25/5 (Accl. control), n = 25/3 (Accl. + Pico145) cells. Middle: The average (mean ± s.e.m.) temperature coefficient was comparable between VMPO^LepR^ acclimated control group and VMPO^LepR^ neurons recorded in the presence of Pico145. Right: Distribution of temperature-insensitive, cold-sensitive (CSN), warm-sensitive (WSN) and silent neurons within acclimated VMPO^LepR^ control group (n = 25/5 cells) and VMPO^LepR^ group recorded in the presence of Pico145 (25/3 cells). f, Depolarisation of non-acclimated VMPO^LepR^ neurons to approximate average membrane potential of acclimated VMPO^LepR^ neurons (-45 mV, requiring the injection of current I+ between +20 pA to +70 pA) did not lead to an increase of firing frequency or increased regularity of firing (represented by the coefficient of variation of interspike interval, ISICoV). n = 22/8 (Non-accl.) and n = 20/2 (Non-accl. + I+) cells. Barplots represent mean ± s.e.m. g, Hyperpolarisation of acclimated VMPO^LepR^ neurons to match the average membrane potential of non-acclimated VMPO^LepR^ neurons (approx. -55 mV, requiring the injection of current I- between -50 to -100 pA) decreased both firing frequency and regularity of firing. Unpaired two-tailed t-test, ***P = 0.0001 for both AP frequency and ISICoV). n = 20/7 (Accl.) and n = 20/2 (Accl. + I-) cells. Barplots represent mean ± s.e.m. Brain slice recordings were conducted at 36°C bath temperature (unless indicated otherwise). In panel (e), “high-K^+^ aCSF” and synaptic blockers were used. In panel (d), “low-K^+^ aCSF” and 33°C bath temperature were used.

**Extended Data Fig. 11.**
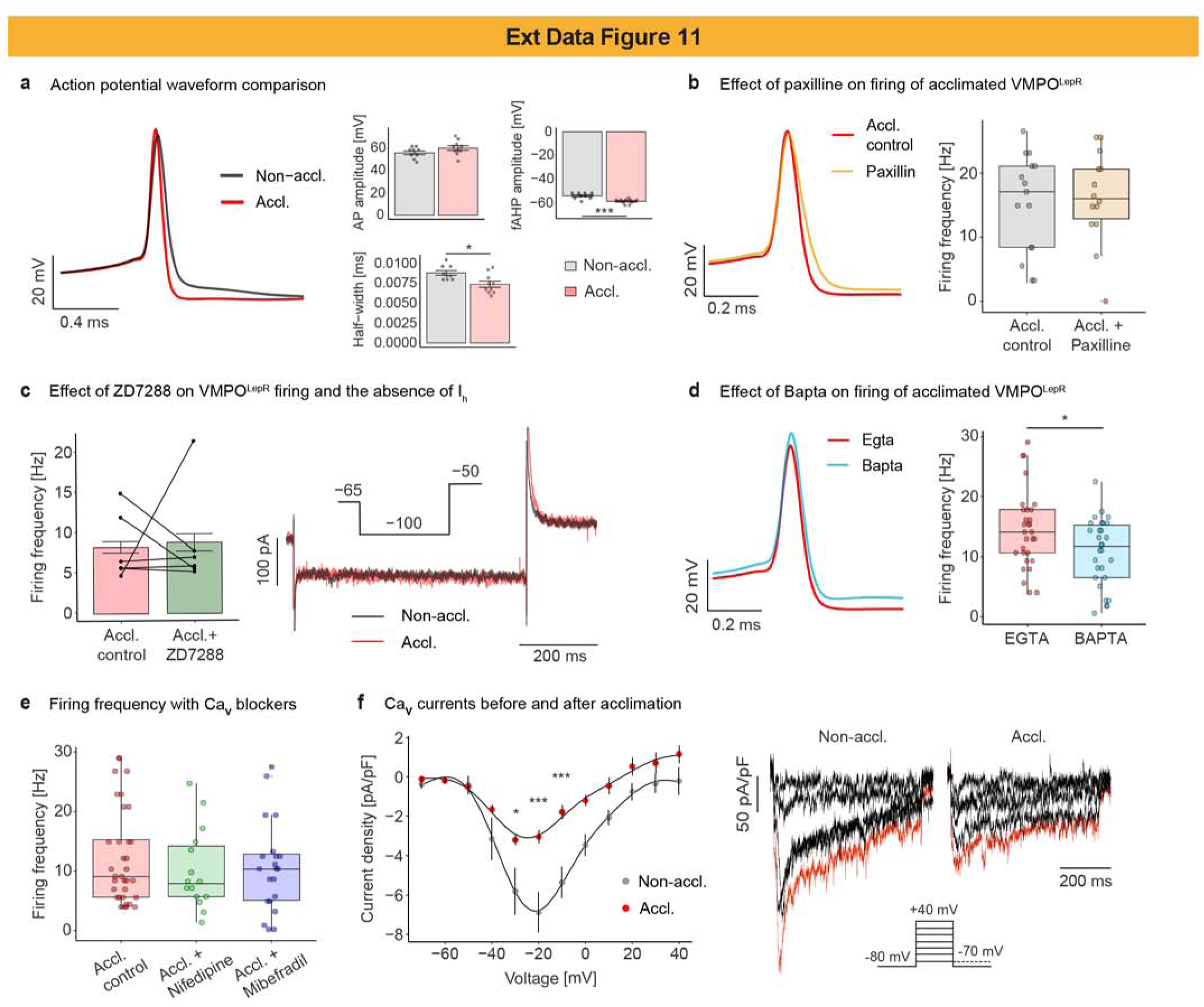
Characterization of action potential waveform and active, voltage-gated conductances in VMPO^LepR^ neurons. a, Left: Alignment of averaged action potential waveforms recorded from non-acclimated (n = 9/4 cells) and acclimated (n = 10/5 cells) VMPO^LepR^ neurons. Right: Average amplitude of action potentials was similar between non-acclimated and acclimated VMPO^LepR^ neurons. AP half-width was found to be shorter in VMPO^LepR^ cells after acclimation (Unpaired two-tailed t-test, *P = 0.0109). Fast afterhyperpolarization (fAHP) was found to have a significantly larger amplitude in the acclimated condition compared to non-acclimated controls (Unpaired two-tailed t-test, ***P = 0.0005). Barplots represent mean ± s.e.m. b, Left: Paxilline (10 µM) reduced fast AHP in acclimated VMPO^LepR^ neurons (yellow trace, n = 5/1 cells, averaged and aligned AP waveforms are shown). Right: Despite reduced fAPH, Paxilline had no impact on the frequency of AP firing in acclimated VMPO^LepR^ cells. n = 15/9 cells for Accl. control and n = 15/1 cells for Accl. + Paxilline. Boxplots represent median and interquartile range. c, Left: HCN channel blocker ZD7288 did not affect firing frequency in acclimated VMPO^LepR^ neurons (n = 6, paired recordings). Barplots represent mean ± s.e.m. Right: Example traces recorded from a non-acclimated and an acclimated VMPO^LepR^ cell using a hyperpolarising step protocol (shown in inset). Hyperpolarization activated currents were not detected in either condition. d, Left: Exchanging EGTA for BAPTA in the pipette solution had a reducing effect on the AHP in acclimated VMPO^LepR^ cells (blue trace, n = 6/2 cells; averaged and aligned AP waveforms are shown). Right: BAPTA slightly reduced the frequency of firing in acclimated VMPO^LepR^ neurons (Unpaired two-tailed t-test, *P = 0.0230; n = 30/10 cells for EGTA and n = 29/4 cells for BAPTA). Boxplots represent median and interquartile range. e, CaV channel blockers Nifedipine (10 µM) and Mibefradil (1 µM) did not significantly alter AP firing frequency of acclimated VMPO^LepR^ neurons (n = 17/8 cells for Accl. control, n = 15/2 cells for Accl. + Nifedipine and 21/2 for Accl. + Mibefradil). Boxplots represent median and interquartile range. f, Voltage-gated Ca^2+^ (CaV) currents recorded in non-acclimated and acclimated VMPO^LepR^ neurons suggested the overall CaV density to be reduced in acclimated VMPO^LepR^ compared to non-acclimated controls. Left: averaged (mean ± s.e.m.) CaV current-voltage relationship obtained for acclimated and non-acclimated VMPO^LepR^ neurons. Two-way ANOVA (effect of acclimation + voltage), P < 0.0001; Tukey’s multiple comparison test, *P = 0.0342 (-30 mV), ***P < 0.0001 (-20 mV), ***P = 0.0001 (- 10 mV). n = 7/2 (Non-accl.) and n = 10/2 (Accl.) cells. Right: Example traces of recorded CaV currents; inset shows voltage step protocol used. Brain slice recordings were conducted at 36°C bath temperature with the exception of panel (c) where “low-K^+^ aCSF” and 33°C bath temperature were used.

**Extended Data Fig. 12.**
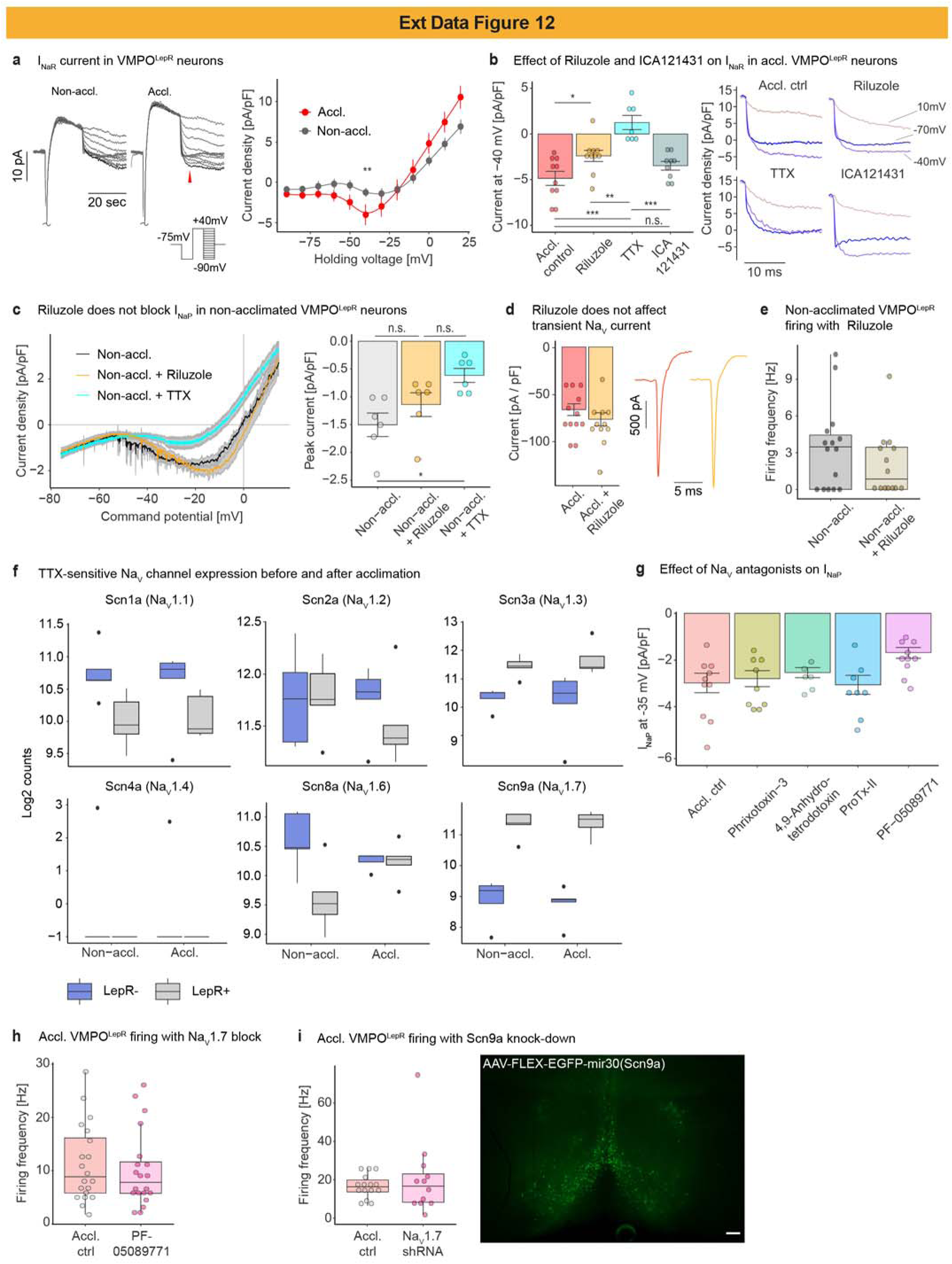
NaV channel current characteristics and gene expression in VMPO^LepR^ neurons. a, Left: example traces of resurgent NaV currents recorded in acclimated and non-acclimated VMPO^LepR^ neurons. Inset: voltage step protocol used to record the resurgent current. Right: Current-voltage relationship for VMPO^LepR^ resurgent NaV current (mean ± s.e.m.). Two-way ANOVA (effect of acclimation * voltage), P < 0.0001; Tukey’s multiple comparison test, **P = 0.0041 (Non-accl.:Accl. at -40 mV). n = 10/2 (Non-accl.), n = 11/2 (Accl.) cells. b, Left: Riluzole reduced and ICA121431 did not affect the TTX-sensitive resurgent NaV current present in acclimated VMPO^LepR^ cells. One-way ANOVA, P = <0.0001; Tukey’s multiple comparison test, *P = 0.0409 (Accl. ctrl : Riluzole), ***P < 0.0001 (Accl. ctrl : TTX), **P = 0.0033 (Riluzole : TTX), ***P = 0.0002 (ICA121431 : TTX). n = 9/2 (Accl. ctrl), n = 10/2 (Riluzole), n = 7/2 (TTX), n = 9/2 (ICA121431). Right: Example traces of resurgent NaV currents at different potentials (-70 mV / blue, - 40 mV / violet and +10 mV / beige) recorded in the acclimated condition and in the presence of Riluzole (10 µM), TTX (1 µM) or ICA121431 (200 nM). c, Left: Traces of INaP (mean ± s.e.m.) in non-acclimated VMPO^LepR^ neurons whereby TTX reduced the current but Riluzole did not have a significant effect. Right panel: quantification of INaP at -35mV based on presented traces. One-way ANOVA, P = 0.015; Tukey’s multiple comparison test, *P = 0.0117 (Non-accl. : Non-accl.+TTX). n = 6/2 (Non-accl.), n = 6/2 (Non-accl.+Riluzole) and n = 6/2 (Non- accl.+TTX) cells. |d, Quantification of peak amplitude (mean ± s.e.m.) and example traces of transient NaV currents recorded in acclimated VMPO^LepR^ neurons (based on the initial depolarising step used in resurgent NaV current recording protocol shown in (a)) with and without Riluzole. n = 12/2 (Accl.) and n = 11/2 (Accl. + Riluzole). Riluzole (10 µM) was found to not affect the amplitude of transient NaV currents. e, Firing frequency of non-acclimated VMPO^LepR^ cells was not affected by Riluzole. n = 15/5 (Non- accl.) and n = 14/1 (Non-accl. + Riluzole). Boxplots represent median and interquartile range. f, Expression analysis of TTX-sensitive NaV channels after Fluorescence-Activated Cell Sorting (FACS) of VMPO LepR^+^ and LepR^-^ cells obtained from non-acclimated and acclimated LepR- Cre;HTB mice; mRNA sequencing results of pooled cells are plotted as normalized log2 values (N = 5 samples for each condition). Boxplots represent median and interquartile range. g, Quantification of INaP amplitude (mean ± s.e.m.) at -35 mV recorded in acclimated VMPO^LepR^ neurons in the presence of the selective NaV channel blockers Phrixotoxin-3 (100 nM), 4,9-Anhydrotetrodotoxin (50 µM), ProTx-II (30 nM) and PF-05089771 (150 nM). One-way ANOVA, P = 0.0471; Tukey’s multiple comparison test, P = 0.0676 (Accl.:PF-05089771). n = 10/2 (Accl.), n = 9/2 (Phrixotoxin-3), n = 6/2 (4,9-Anhydrotetrodotoxin), n = 8/2 (ProTx-II) and n = 10/2 (PF-05089771) cells. h, Neither PF-05089771 (left panel) nor NaV1.7 knock-down via shRNA AAV (middle panel) affected AP firing frequency in acclimated VMPO^LepR^ neurons. n = 20 cells each for PF-05089771; n = 15 vs n = 11 cells for NaV1.7 shRNA. Boxplots represent median and interquartile range. Right panel shows a representative image of the AAV9-pCAG-FLEX-EGFP-mir30(Scn9a) viral construct expressed in LepR-cre mouse VMPO; scale bar 100 um. Brain slice recordings were conducted at 36°C bath temperature with the exception of panel (h) where “low-K^+^ aCSF” and 33°C bath temperature were used.

**Extended Data Fig. 13.**
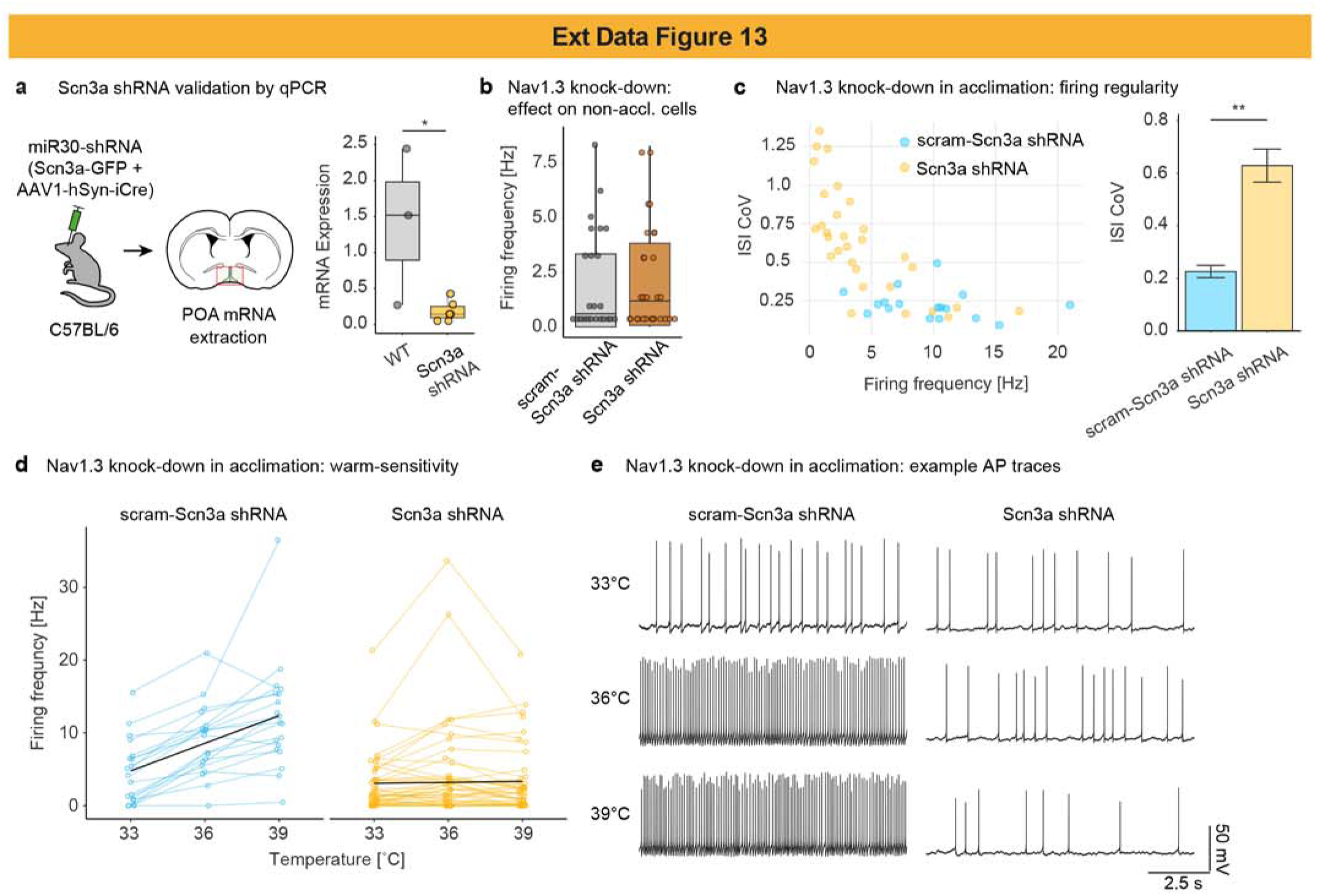
Validation and electrophysiological characterization of Nav1.3 knock- down. a, Left: in order to test the efficacy of shRNA against the NaV1.3 mRNA, Cre-dependent shRNA-carrying AAVs were co-injected together with AAV encoding the Cre recombinase into the POA of C57/BL6 mice. Following 3 weeks of virus expression, mRNA was extracted from the POA tissue. Non-injected C57BL/6 mouse POA tissue served as control. Right: boxplot (median and interquartile range) of relative NaV1.3 mRNA expression normalized to the housekeeping genes Tubb3 and Ube2l3 in mouse POA. Unpaired two-tailed t-test, *P = 0.0226. N = 3 (WT) and N = 6 (Scn3a shRNA) mice. b, Firing frequencies of non-acclimated VMPO^LepR^ neurons expressing either scrambled shRNA (scram-Scn3a shRNA; n = 27/2) or functional shRNAs against Nav1.3 (Scn3a shRNA; n = 27/3) at 36 °C. c, Left: plot showing AP firing frequency vs regularity of firing in VMPO^LepR^ neurons expressing either functional shRNAs against NaV1.3 / Scn3a mRNA or scrambled control. Right: quantification of the firing regularity between the two groups. Unpaired two-tailed t-test, *P = 0.0226. n = 30/5 (Scn3a shRNA) and n = 17/3 (scram-Scn3a shRNA) cells. d, Firing frequencies of acclimated VMPO^LepR^ neurons expressing either scrambled shRNA (scram- Scn3a shRNA; n = 19/3) or functional shRNAs against NaV1.3 (Scn3a shRNA; n = 47/7) at the three indicated bath temperatures. Individual cells are plotted in color; black lines represent linear regression for each group. Slope=temperature coefficient (TC) = 1.2651 for scram-Scn3a shRNA and TC = 0.0474 for Scn3a shRNA, demonstrating that NaV1.3 knock-down significantly reduced warm sensitivity of acclimated VMPO^LepR^. e, Example traces of spontaneous warm-sensitive activity of acclimated VMPO^LepR^ expressing either scrambled or functional shRNAs against NaV1.3 recorded at 33°C, 36°C and 39°C.

**Extended Data Fig. 14.**
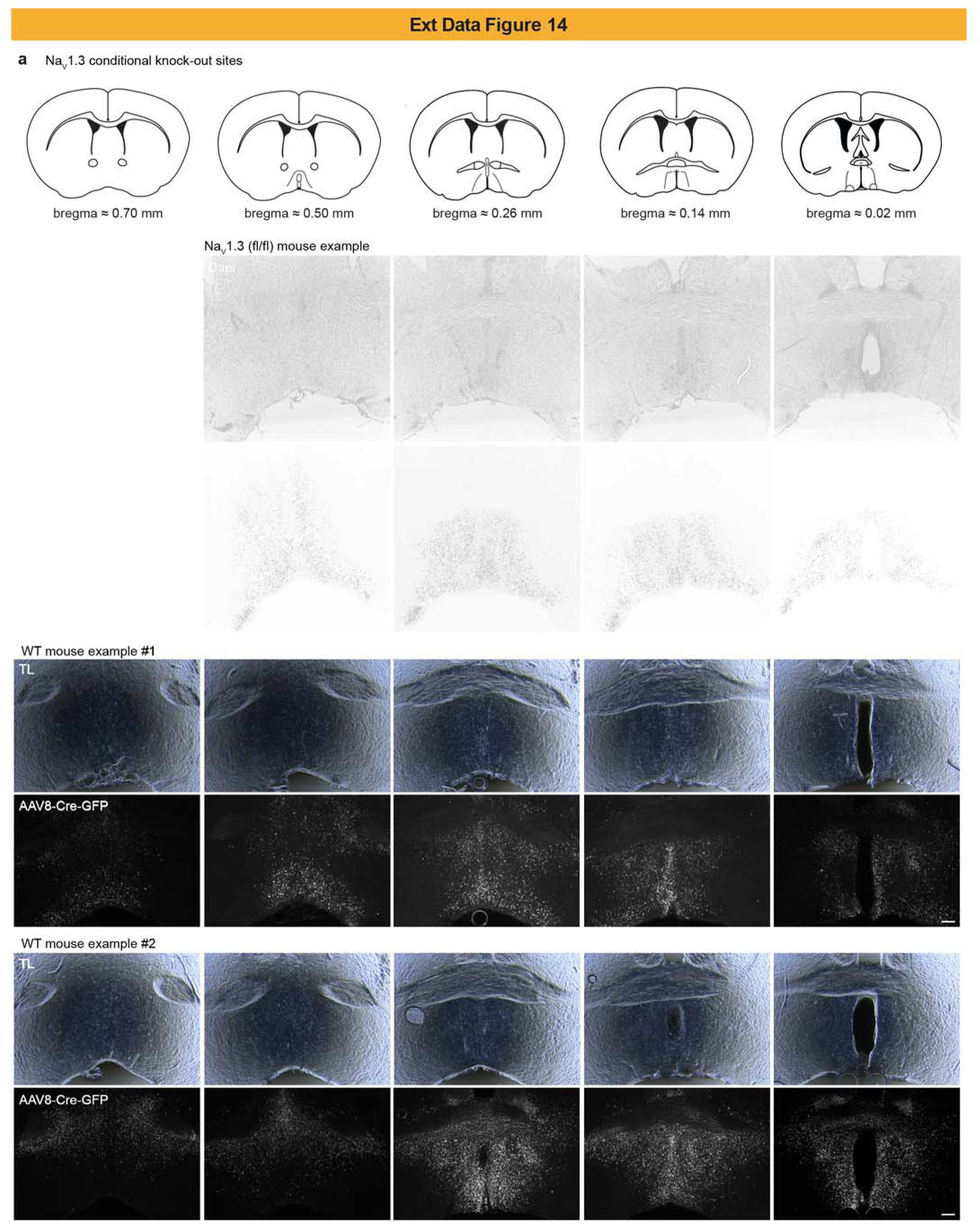
Histological Validation of AAV-Cre expression in NaV1.3^fl/fl^ and wildtype mice. a, Images showing the site of virally-delivered Cre recombinase by GFP expression in NaV1.3^fl/fl^ and wildtype control mice. Cartoons on top show antero-posterior coordinates of the mouse brain, reflecting the stereotactic injection sites. For wildtypes the DIC images are shown above the fluorescent images (TL = transmitted light); Scale bars = 100 μm.

**Extended Data Fig. 15.**
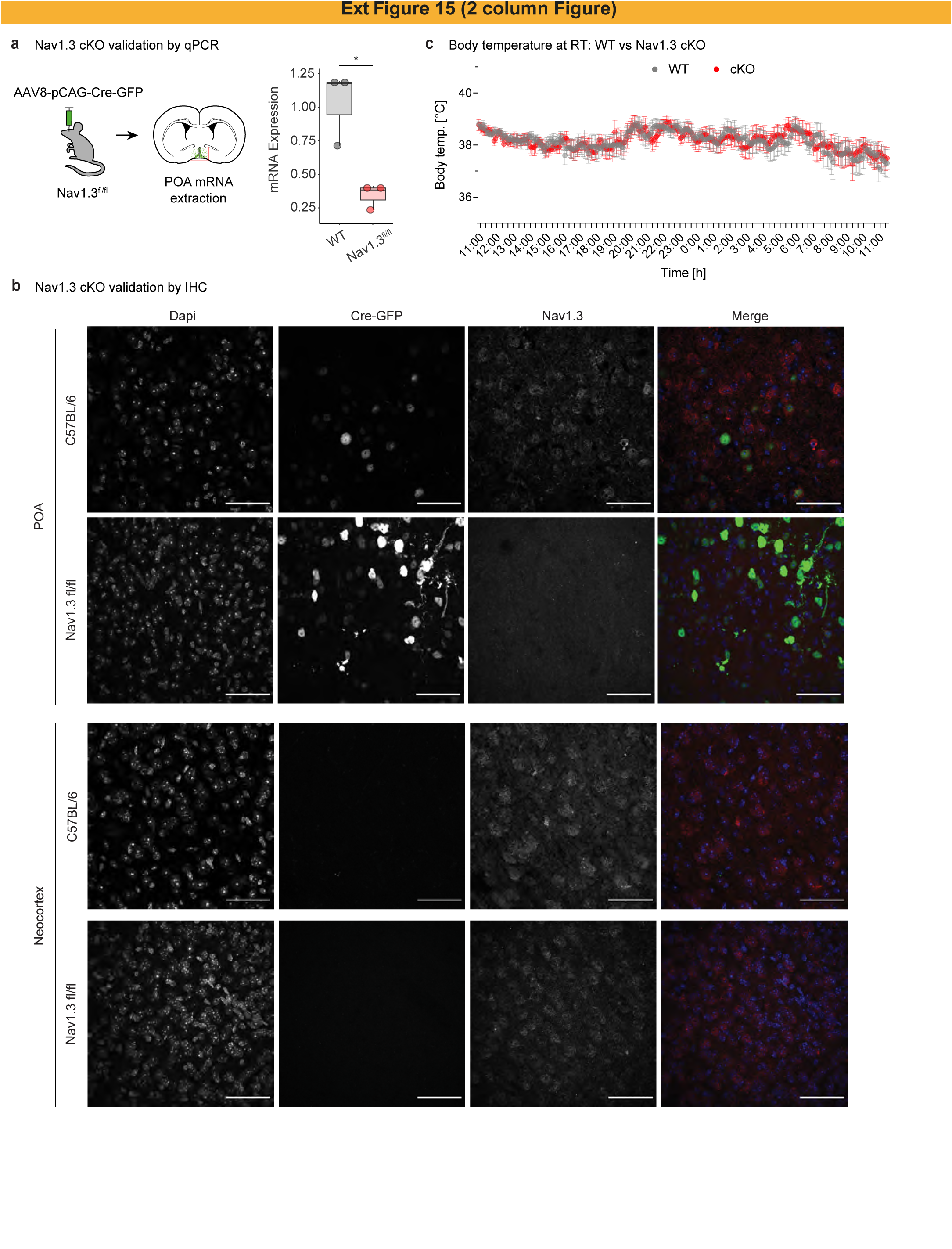
Verification and characterization of NaV1.3 conditional Knock-out. a, Left: in order to test the efficacy of Scn3a conditional knock-out (cKO), Cre recombinase-carrying AAVs were injected into the POA of Nav1.3^fl/fl^ mice and C57BL/6 wildtype (WT) mice. Following 3 weeks of virus expression, mRNA was extracted from the POA tissue. Right: boxplot (median and interquartile range) showing qPCR to assess relative NaV1.3 mRNA expression normalized to the housekeeping genes Tubb3 and Ube2l3. Unpaired two-tailed t-test, *P = 0.0147. N = 3 Cre-injected wildtype (WT) and N = 3 Cre-injected Nav1.3^fl/fl^ (NaV1.3 cKO) mice. b, Average (mean ± s.e.m.) body temperature measured telemetrically of Nav1.3 cKO and control mice at room temperature. N = 9 mice. c, Immunohistochemistry in brain slices using antibodies detecting GFP and Nav1.3 and counterstained with DAPI. Merged images show GFP in green and Nav1.3 in red. 60x images show staining signal in POA and neocortex of AAV-Cre injected C57BL/6 wildtype (WT) and AAV-Cre injected NaV1.3^fl/fl^ (NaV1.3 cKO) mice. Shown are representative images of mice stereotaxically injected with AAV8-Cre-GFP into the POA (but not into the neocortex, a brain region that therefore served as internal control). Scale bar = 50 μm.

**Extended Data Fig. 16.**
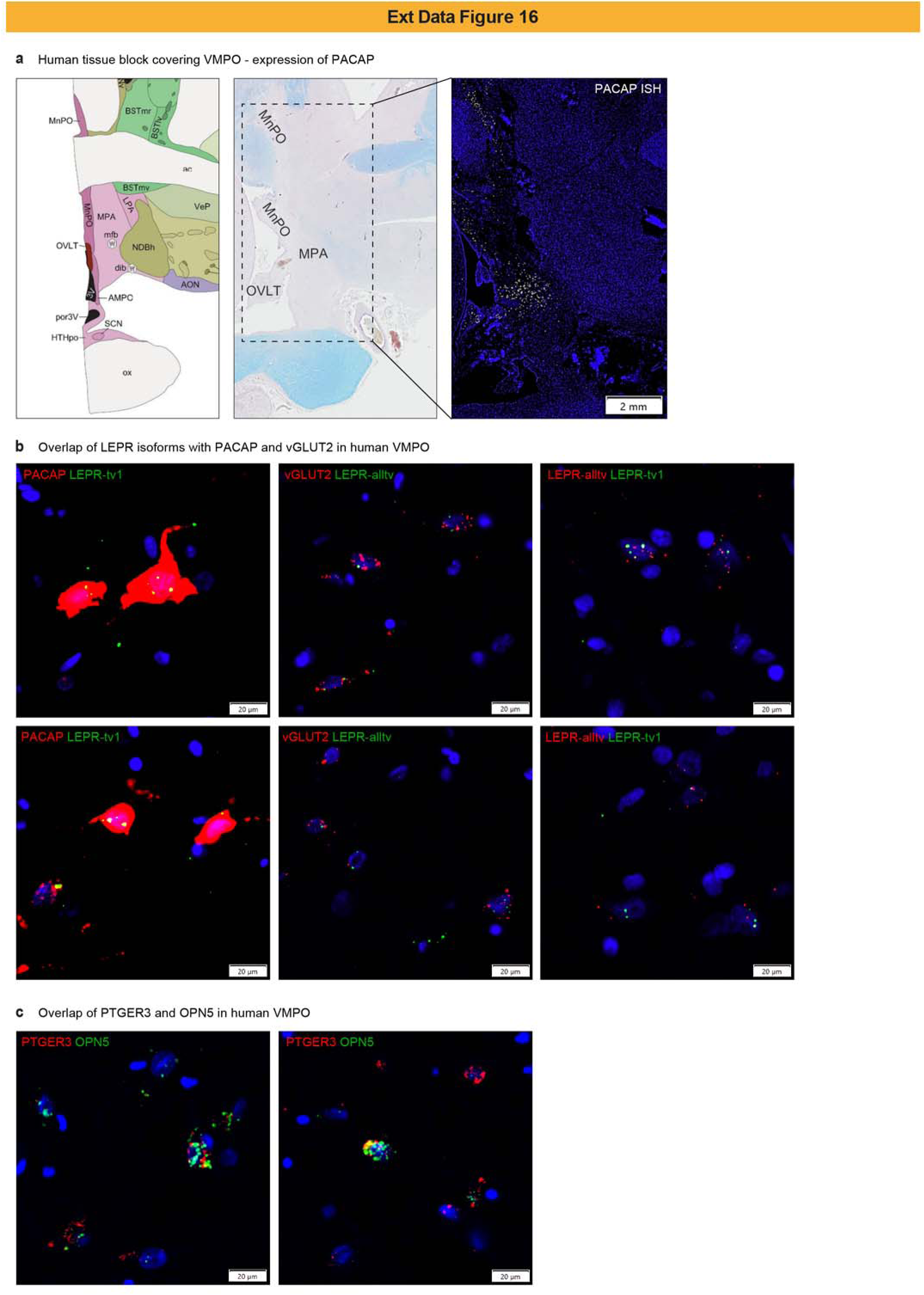
Human VMPO expression analyses of “QPLOT” marker genes LEPR, vGLUT2, PACAP, PTGER3 and OPN5. a, Left: Allen Brain Atlas annotation of human VMPO. Middle: human tissue block covering preoptic areas MnPO/MPA/OVLT (LFB/HE stain). Right: ISH of the VMPO brain section showed in the middle panel stained for PACAP (ADCYAP1). b, Additional examples of RNAscope ISH showing LEPR co-expression in human VMPO. Two different LEPR ISH probes (see Methods) were co-labeled with PACAP (ADCYAP1) and vGLUT2 (SLC17A6). Co-expression of mRNA is shown in yellow. c, RNAscope ISH showing co-expression (yellow) of other thermoregulatory neuronal markers, the Prostaglandin E Receptor 3 (PTGER3) and Opsin 5 (OPN5), in human VMPO. Images were taken from 1 tissue section/staining from a single human donor.

